# The Tung Tree (*Vernicia Fordii*) Genome Provides A Resource for Understanding Genome Evolution and Oil Improvement

**DOI:** 10.1101/2019.12.17.877803

**Authors:** Lin Zhang, Meilan Liu, Hongxu Long, Wei Dong, Asher Pasha, Eddi Esteban, Wenying Li, Xiaoming Yang, Ze Li, Aixia Song, Duo Ran, Guang Zhao, Yanling Zeng, Hao Chen, Ming Zou, Jingjing Li, Fan Liang, Meili Xie, Jiang Hu, Depeng Wang, Heping Cao, Nicholas J. Provart, Liangsheng Zhang, Xiaofeng Tan

## Abstract

Tung tree (*Vernicia fordii*) is an economically important woody oil plant that produces tung oil containing a high proportion of eleostearic acid (∼80%). Here we report a high-quality, chromosome-scale tung tree genome sequence of 1.12 Gb with 28,422 predicted genes and over 73% repeat sequences. Tung tree genome was assembled by combining Illumina short reads, PacBio single-molecule real-time long reads and Hi-C sequencing data. Insertion time analysis revealed that the repeat-driven tung tree genome expansion might be due to long standing long terminal repeat (LTR) retrotransposon bursts and lack of efficient DNA deletion mechanisms. An electronic fluorescent pictographic (eFP) browser was generated based on genomic and RNA-seq data from 17 various tissues and developmental stages. We identified 88 nucleotide-binding site (NBS)-encoding resistance genes, of which 17 genes may help the tung tree resist the *Fusarium* wilt shortly after infection. A total of 651 oil-related genes were identified and 88 of them were predicted to be directly involved in tung oil biosynthesis. The fewer phosphoenolpyruvate carboxykinase (PEPC) genes, and synergistic effects between transcription factors and oil biosynthesis-related genes may contribute to high oil content in tung seeds. The tung tree genome should provide valuable resources for molecular breeding and genetic improvement.

## Introduction

Tung tree (*Vernicia fordii*), a woody oil plant native to China, is widely distributed in the subtropical area. Tung tree taxonomically belongs to the Euphorbiaceae family, along with several other economically important species including cassava (*Manihot esculenta*), castor oil plant (*Ricinus communis*), rubber tree (*Hevea brasiliensis*) and physic nut (*Jatropha curcas*). Species commonly referred to as tung trees include three major subspecies (*V. fordii*, *V. montana*, and *V. cordata*), of which *V. fordii* is the most widely cultivated species due to wide geographic distribution, medium stature for easy plantation management, and high-quality oil production. Tung trees have been planted for tung oil production or ornamental purpose for more than 1000 years in China [1]. Tung trees have been widely distributed in 16 Chinese provinces and many countries after they were introduced into America, Argentina, Paraguay and other countries for plantation and tung oil production at the beginning of the 20th century [1] (Figure S1).

Tung seeds contain 50%−60% tung oil, which is composed of approximately 80% α-eleostearic acid (α-ESA), a type of unusual fatty acid. As the major component in tung oil, α-ESA has three conjugated double bonds (9 cis, 11 trans, 13 trans), and thus is easily oxidized. Due to its excellent characteristics, tung oil has been widely used as a drying ingredient in paints, varnishes, coating and finishes since ancient times [2]. Tung oil also can be used for synthesizing thermosetting polymers and resins with superior properties [3, 4], and has been proposed as a potential source of biodiesel [5–7]. Tung oil was one of the chief exports until 1980s, and then declined due to the development of chemical coatings. Interestingly, tung oil has been attracted world-wide attention in recent years due to production security, environmental concerns, and negative effect of synthetic chemical coatings on human health [8–10]. New technologies have been developed to improve the performance of tung oil-based coatings [3,11,12].

As an oil crop, economic traits involved in fatty acid biosynthesis and oil accumulation are the targets of improved breeding efficiency for tung tree. However, identification of important genes, gene families and marker loci associated with oil content, fatty acid composition, and fruit yield has been hampered due to a lack of genetic and genomic information. Only a few functional genes, mainly involved in the formation and regulation of fatty acids such as *fatty acid desaturase* (*FAD2*, *FAD3*, *FADx*) and *diacylglycerol acyltransferase* (*DGAT*), have been investigated to date [13–17].

In the present study, we report the sequencing and assembly of *V. fordii* genome, which was achieved by combining whole-genome shotgun sequencing of Illumina short reads and real-time (SMRT) long reads on a Pacific Biosciences (PacBio) platform. We also used a Hi-C map to cluster the majority of the assembled contigs onto 11 pseudochromosomes. We conducted evolutionary comparisons and comprehensive transcriptome analysis of genes involved in oil biosynthesis to elucidate the genetic characteristics of oil synthesis and genetic difference as compared to other plant species.

## Results

### Genome sequencing, assembly and validation

The self-bred progeny ‘VF1-12’ of *V. fordii* cv. Putaotong was used for genome sequencing (File S1). The genome of *V. fordii* was estimated to be 1.31 Gb in size with a low heterozygous rate of 0.0976% (Tables S2 and S3; File S2; Figure S4). After removing low-quality reads, we obtained a total of 177.68 Gb of high quality data, including 160.21 Gb of Illumina sequencing data and 187.47 Gb of SMART data, corresponding 135.73 × coverage of the tung tree genome (Table S4; Figure S5). The assembled tung tree genome was 1.12 Gb covering 85% of the estimated genome size, and contained 34,773 contigs with a maximum length of 544.11 Kb and 4,577 scaffolds with a maximum length of 5.09 Mb (**Table 1**; Table S5). Among them, 3,333 contigs and 29,721 scaffolds were more than 2 Kb in length (Table S5). After Hi-C data assessment and assembly, 1.06 Gb (95.15%) of the genome sequences were anchored onto 11 pseudochromosomes, with maximum clustered sequence lengths, minimum clustered sequence lengths and scaffold N50 of 120.57 Mb, 63.43 Mb and 87.15 Mb, respectively (Table 1; Tables S6−S11; **Figure 1**; Figures S6 and S7).

**Figure 1.**
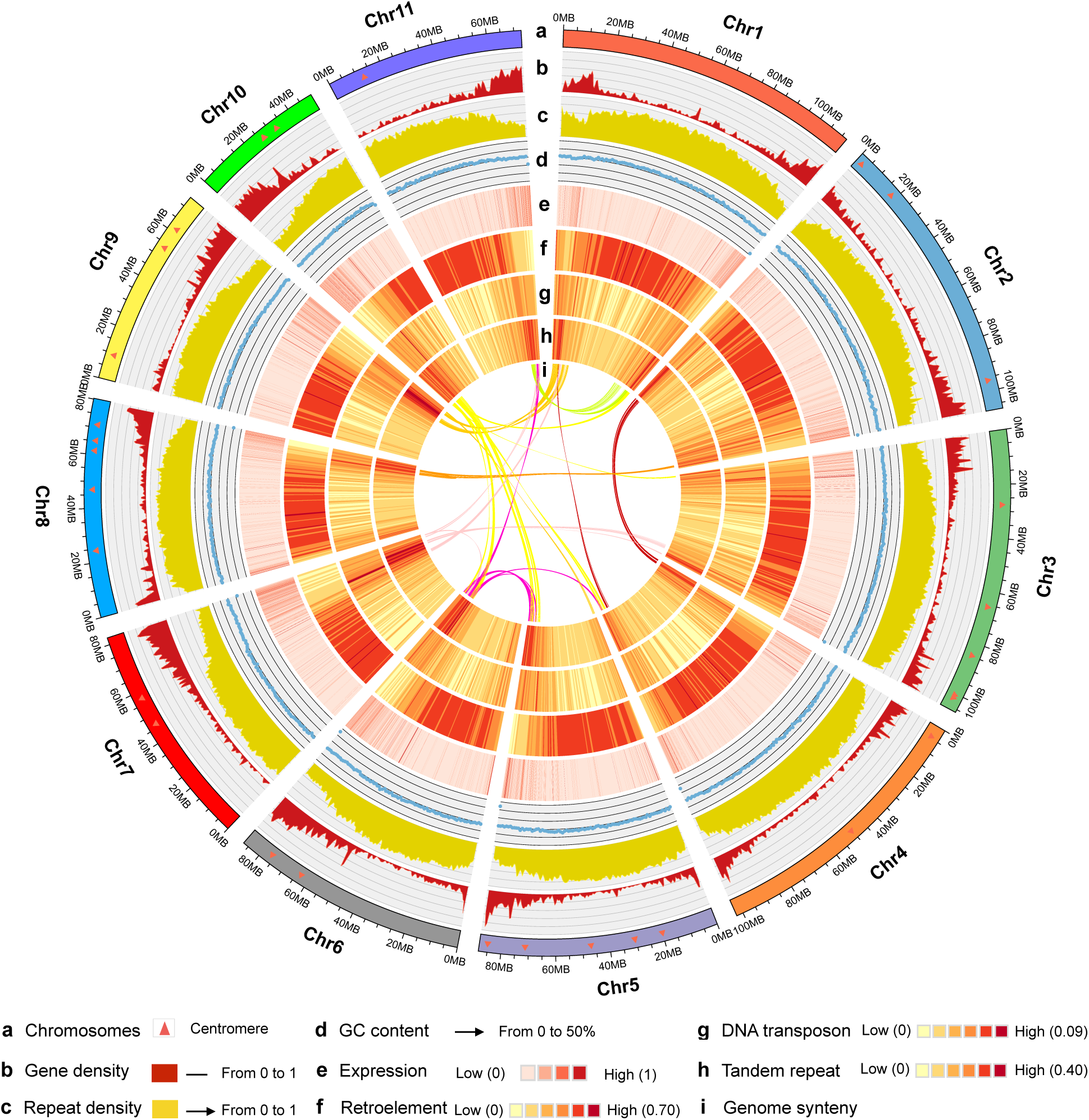
The genomic landscape of tung tree. The features from outside to inside are pseudochromosomes (**a**), gene density (0−1) (**b**), repeat density (0−1) (**c**), GC content (0%−50%) (**d**), expression (0−1) (**e**), retroelement (0−0.70) (**f**), DNA transposon (0−0.09) (**g**), tandem repeat (0−0.40) (**h**), genome synteny (**i**); Intra-genome collinear blocks with gene pairs numbering more than 20 are highlighted with arcs in the middle of the diagram. Circos was used to construct the diagram. All distributions were drawn using a window size of 1 Mb with the exception of expression, which was drawn using a window of 50 Kb. Chr, chromosome.

The CEGMA prediction indicated that 87.9% complete elements and 95.97% partial elements in tung tree genome could be hit for the 248 most conserved genes (Table S12). The BUSCO analysis showed that 1,379 (95.7%) of BUSCO genes were complete, of which 1338 (92.9%) and 41 (2.8%) were single-copy and duplicated, respectively (Table S13). RNA-seq data showed that 90.36%, 96.83% of flower samples 1 and 2 unigenes, 95.35%, 95.50%, and 96.48% of seed samples 1–3 unigenes showed good alignments to the assembled tung tree genome with mapping rate > 90%, respectively (Tables S14−S19). Furthermore, 88.3% to 95.6% of the reads from the five samples could be mapped to our genome assembly (Table S20). The validation results suggested that our tung tree genome assembly was of high quality in this study.

### Genome annotations

In total, 28,422 genes were predicted with an average transcript length of 3,785 bp, average CDS length of 1,034 bp, average exon number of 4.85 per gene, average exon length of 213 bp, and average intron length of 714 bp (Table 1; Table S21; Figure S8). The GC content was 31.93% across the genome, 41.91% in coding sequences and 31.16% in intron regions (Table 1; Tables S22−S24). BUSCO analysis showed that 1290 complete BUSCOs (89.6%) could be searched of all BUSCO groups, indicating that most of the gene models were complete (Table S25).

Among the total 28,422 genes, 23,143 genes (81.4%) were functionally annotated. Tremble, Swissprot and NR allowed the annotation of 79.6%, 63.8%, and 81.1% of all genes, respectively (Table S26). Gene ontology (GO) annotation revealed that 12,581 genes could be grouped into three categories with 65.97% in molecular function (GO:0003674), 20.1% in cellular component (GO:0005575), and 58.52% in biological process (GO:0008150) (Figure S9). We were able to use kyoto encyclopedia of genes and genomes (KEGG) to annotate 6835 genes to 235 pathways, of which oil biosynthesis and metabolism-related glycerolipid metabolism (ko00561), fatty acid biosynthesis (ko00061), fatty acid elongation (ko00062), and fatty acid degradation (ko00071) were of particular interest in this paper (Table S27).

In addition, we identified several types of non-coding RNAs in tung tree genome, including 465 microRNA (miRNA) genes, 740 transfer RNA (tRNA) genes, 116 ribosomal RNA (rRNA) genes, and 1414 small nuclear RNA (snRNA) genes (Table S28).

### Gene family evolution and phylogeny

A total of 22,991 tung tree genes clustered into 15,038 gene families including 8,865 gene families shared by all eight species, and 635 tung tree-unique families and 5,431 tung tree-specific unclustered genes (Table S29). GO annotation of the tung tree-specific families showed that the genes involved in macromolecule metabolic processes (GO:0043170), cellular macromolecule metabolic processes (GO:0044260) and protein metabolic processes (GO:0019538) were highly enriched (Table S30; Figure S10). A total of 933 genes could be annotated using KEGG database, of which 586 genes were mapped to KEGG pathways. We observed KEGG enrichment in translation (110), carbohydrate metabolism (61), biosynthesis of other secondary metabolites (42), amino acid metabolism (44), folding, sorting and degradation (44), signal transduction (43), biosynthesis of other secondary metabolites (42), and environmental adaptation (36) (Table S31). We identified 11,985 gene families that were shared among the five Euphorbiaceae species (Figure S11A). The tung tree shared 13,408, 13,387, 13,519, and 13,216 gene families with *J. curcas, H. brasiliensis, M. esculenta,* and *R. communis*, respectively, of which 9,778, 6,643, 7,980, and 10,675 gene families had a one-to-one orthologous relationship (Figure S11A). Additionally, compared with *A. thaliana, P. trichocarpa,* and *V. vinifera*, 3,421 gene families were found to be specific to Euphorbiaceae (Figure S11B).

A phylogenetic tree was generated based on a total of 2,085 single gene families among the eight species (**Figure 2A**; Figure S12). We estimated that *V. fordii* and *J. curcas* diverged around 34.55 million years ago (Mya) (Figure 2A). These data indicate that *V. fordii* is more closely related to *J. curcas* than *M. esculenta, R. communis,* and *H. brasiliensis* in Euphorbiaceae family, which is consistent with their phylogenetic classification based on morphological characteristics.

**Figure 2.**
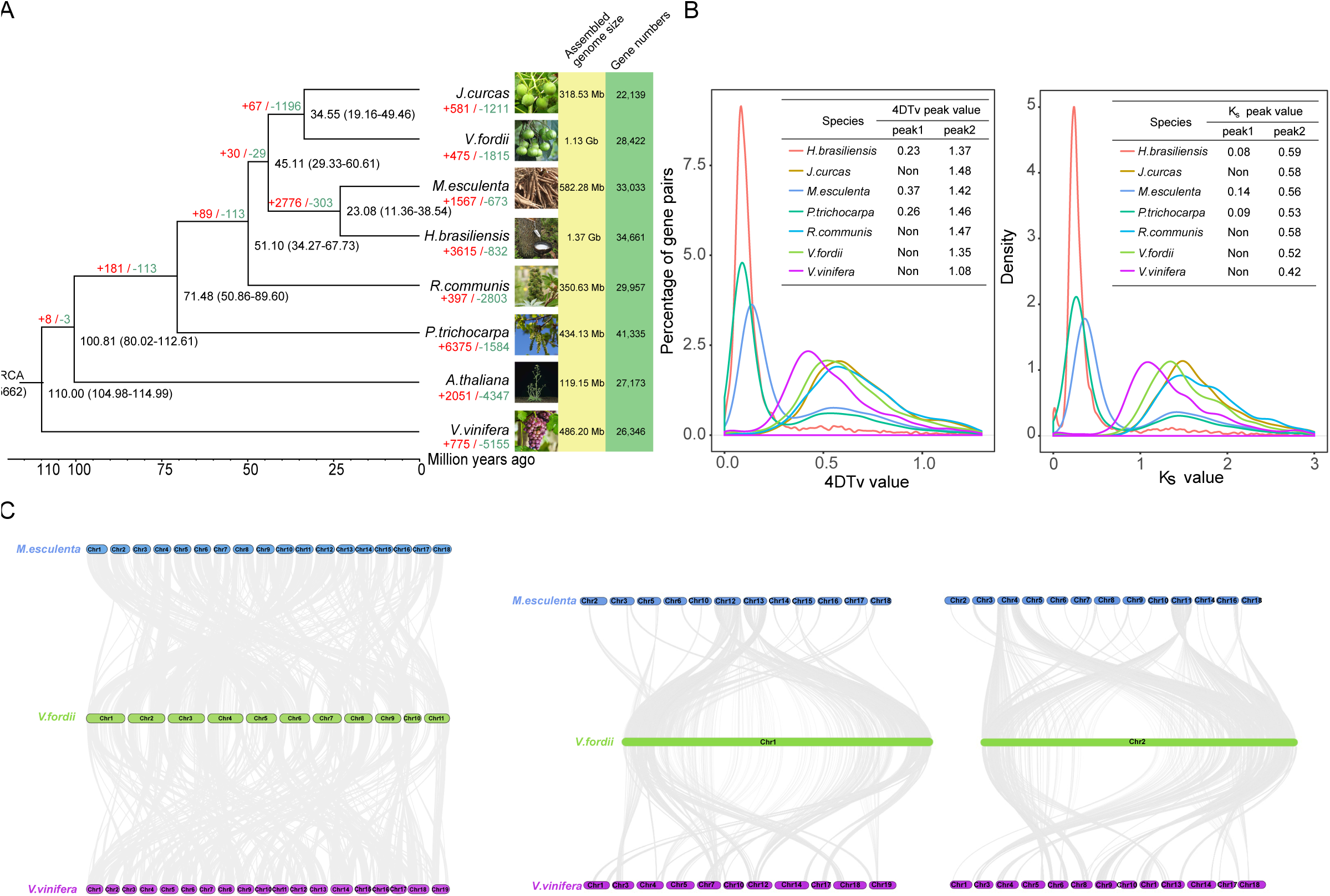
Evolution of tung tree genome. **A**. Phylogenetic tree of tung tree and 7 other plant species based on orthologues of single-copy gene families. The number in parentheses at each branch point represents the divergence time (Mya). The number at the root (15,662) represents the number of gene families in the common ancestor. The value above each branch indicates the number of gene family expansion/contraction at each round of genome duplication after divergence from the common ancestor. Bootstrap value for each node is 100. **B**. Density distribution of 4DTv and Ks for paralogous genes. The peak value is shown in the inset. “non” means no peak value. **C**. Collinear relationship of *V. fordii*, *M. esculenta* and *V. vinifera*. Gray line connects matched gene pairs. The chromosomes of tung tree, cassava and grapevine were assigned with green, blue and purple, respectively. The annotated genes were clustered into gene families among eight sequenced whole genomes including *A. thaliana, P. trichocarpa, V. vinifera* and five Euphorbiaceae species *i.e*. *V. fordii, R. communis, M. esculenta, H. brasiliensis, and J. curcas.*

The expansion and contraction of gene families occur since plants are subjected to selection pressure during their evolution, thereby playing significant roles in plant phenotypic diversification [18]. Expansion and contraction analysis on 15,662 shared gene families based on the phylogenetic tree produced 475 expanded gene families encompassing 1,612 genes, and 1,815 contracted families in tung tree as compared to other plant species (Figure 2A). Of the 1,612 expanded genes, 839 could be annotated using the GO database. GO annotation revealed highly enriched genes related to macromolecule metabolic processes (GO:0043170), cellular macromolecule metabolic processes (GO:0044260), and nucleotide binding (GO:0000166) (Table S32; Figure S13).

The Ka/Ks ratio, also called ω or dN/dS, represents the number of non-synonymous substitutions per non-synonymous site (Ka) to the number of synonymous substitutions per synonymous site (Ks), indicating selective pressure acting on a protein-coding gene in genetics. The values of Ks and Ka substitution rates and the Ka/Ks ratio were estimated in each homologous cluster. A total of 586 positively selected genes (PSGs) in tung tree genome were identified, of which 475 were annotated using Swissprot functions (Table S33). GO annotation revealed that the PSGs related to pigment metabolic processes (GO:0042440), mitochondrial membrane (GO:0031966) and nuclear part (GO:0044428) are highly enriched (Table S34; Figure S14).

### Whole genome duplication and collinearity

All of the seven species showed peak 2 with the values ranging from 1.08 to 1.48 for 4DTV analysis, and 0.42 to 0.59 for Ks analysis (**Figure 3**). However, peak 1 was only observed in *V. fordii*, *J. curcas* and *R. communis* (Figure 2B). The results suggest that only an ancient genome triplication event (*i.e.*, γ event shared by core eudicots) and no recent independent whole-genome duplication (WGD) events occurred in the subsequent ∼ 34.55 Mya evolutionary history in the tung tree lineage.

**Figure 3.**
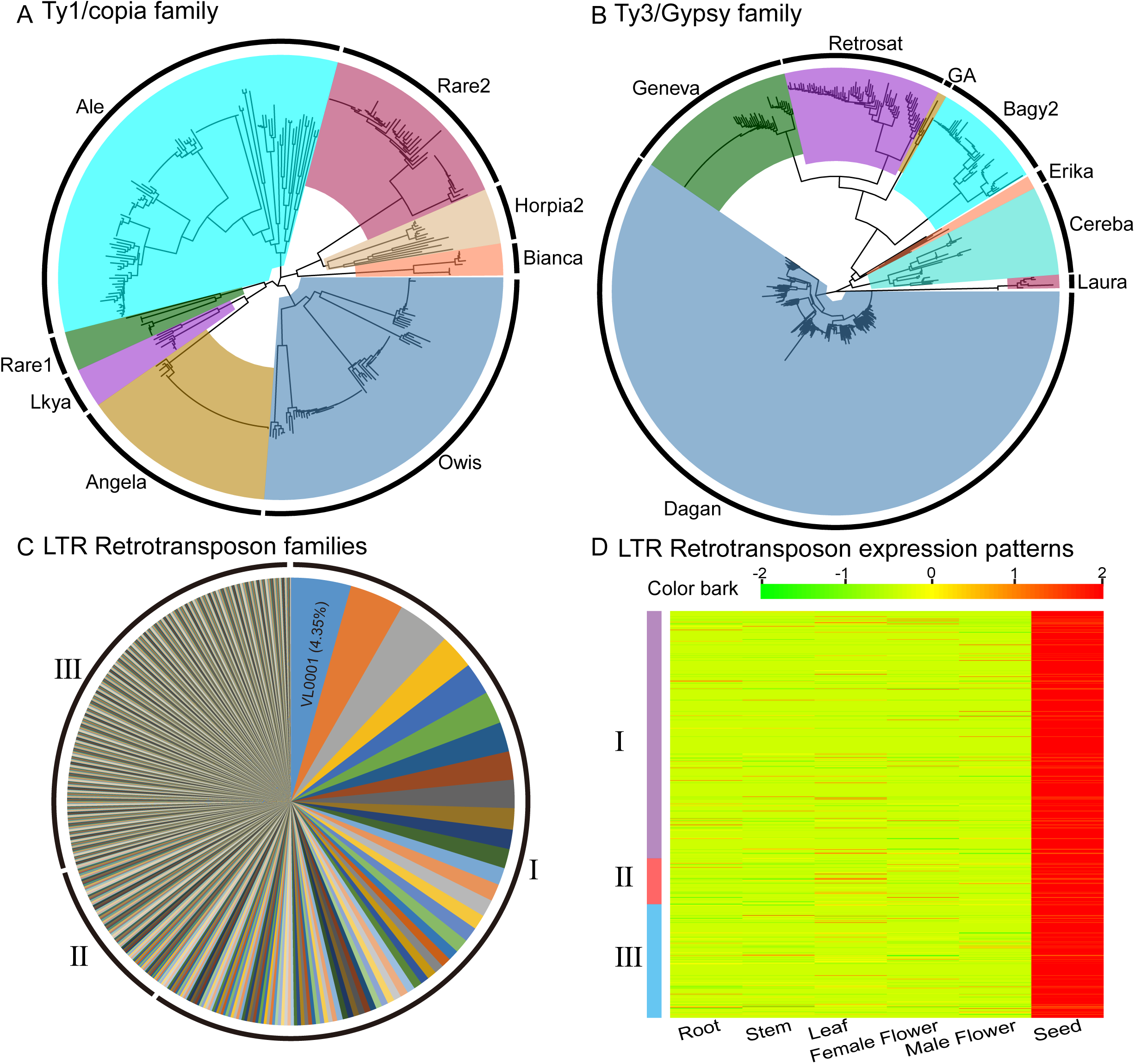
Analysis of the LTR Retrotransposons in the tung tree genome. **A**. The neighbor-joining tree based on 347 *Ty1/copia* sequences; **B**. The neighbor-joining tree based non 622 *Ty3/gypsy* sequences. **C**. Proportions of LTR Retrotransposon families by copy number in the tung tree genome. **D**. Heat map of expression patterns of 701 LTR Retrotransposons. All aligned sequences correspond to the RT domains without premature termination codon. LTR family names and their proportion are indicated. I, II, and III indicate high-copy families (>= 5 intact members), median-copy families (2–4 intact members) and single-copy families, respectively.

Plotting collinear regions of tung tree with itself showed that only 122 syntenic blocks containing 2,010 collinear gene pairs were identified in the tung tree genome (Figure 1; Table S35). A total of 3,559 genes comprised the collinear gene pairs, accounted for only 12.52% of tung tree genes, which is similar with *V. vinifera* (13.91%) and much lower than *M. esculenta* (33.86%) (Tables S36 and S37). The low collinear rate of tung tree genome suggests that a minority of the tung tree genome was duplicated during its evolution, which is consistent with the finding that the tung tree did not undergo a recent WGD event.

The tung tree genome generally showed a one-to-one and one-to-two syntenic relationships with *V. vinifera* (one duplication) and *M. esculenta* (two duplications), respectively (Figure 2C). Tung tree genome shared a total of 694 syntenic blocks, containing 22,133 collinear gene pairs with *M. esculenta*, and 589 syntenic blocks containing 14,570 collinear gene pairs with *V. vinifera* (Figure 2C; Figures S15 and S16). Most collinear regions between tung tree and *M. esculenta* revealed that one chromosome in tung tree corresponded to two chromosomes in *M. esculenta* (Figure 2C; Figure S17). For instance, VfChr1 in tung tree corresponded to MeChr12 and MeChr13 in cassava, and, similarly, VfChr2 to MeChr4 and MeChr11, VfChr3 to MeChr7 and MeChr10, VfChr5 to MeChr1 and MeChr2, as well as VfChr6 to MeChr1 and MeChr5, respectively. These results indicate that that VfChr1, VfChr2, VfChr3, and VfChr5 of tung tree might be formed by fragmentation and recombination of ancestral chromosomes. The collinear regions between tung tree and *V. vinifera* did not exhibit the remarkable corresponding chromosome relationships, in contrast to those between tung tree and *M. esculenta*.

### Repeat-driven genome expansion

Tung tree had larger genome size than physic nut and castor bean, which was mainly attributed to repeat expansion in tung tree genome. Repetitive element analysis showed that tung tree genome harbored the greatest repeat content (73.34%) among the five sequenced Euphorbiaceae species (Table S40), which was slightly higher than the rubber tree (71%) [19], and much higher than the castor oil plant (50.33%) [20], Physic nut (49.8%) [21], and cassava (less than 40%) [22]. The repeat sequences were distributed at both ends of each tung tree chromosome (Figure 1). We identified 66,3931 simple sequence repeats (SSRs) in the tung tree genome. The annotated SSRs were mostly mononucleotide (39.62%) and dinucleotide (13.38%) (File S3). Retroelements comprised the majority (51.89%) of the tung tree genome, of which 50.77% belonged to long terminal repeat (LTR) retrotransposons (Table S41). Of the repeat sequences, two types of LTR retrotransposons, *Ty1/Copia* (84,180 in number) and *Ty3/Gypsy* (284,597 in number) were most abundant, accounting for 15.13% and 53.46%, respectively (Figure 3A and B; File S3; Table S41). The *Ty1/Copia* and *Ty3/Gypsy* were ∼ 0.53 Gb of total length, occupying 50.31% of the assembled tung tree genome.

Kimura analysis showed that two LTR retrotransposons (*Ty1/Copia* and *Ty3/Gypsy*) and DNA transposons were almost simultaneously amplified, with similar peaks for amplification bursts (Figure S18). Insertion time analysis of intact LTR retrotransposons indicated that both of *Ty1/Copia* and *Ty3/Gypsy* experienced multiple bursts over the last 3-4 Mya and they were younger than other unclassified transposable elements (File S3; Figures S19 and S20). In addition, median-copy families and high-copy families were younger than single-copy families (Figure S21). In light of our analysis, the dramatic expansion in tung tree genome size might be due to long standing LTR retrotransposon bursts and lack of efficient DNA deletion mechanisms. VL0001 was the largest *Ty3/Gypsy* family with 130 copies, accounting for 7.54% of the high-copy families and 4.35% of LTR retrotransposons (Figure 3C; Table S42).

Based on our RNA-seq data, 1,738 out of the total 2,991 LTR retrotransposons were expressed across six tissues. *Ty3/gypsy* LTR retrotransposons generally exhibited higher expression levels than *Ty1/Copia* retrotransposons, ranging from 0.71-fold in seed to 4.09-fold in leaf with approximately two-fold higher on average (File S3; Table S44). Among the 1,738 LTR retrotransposons, 701 showed the highest expression level in seeds, of which 60.77% belonged to high-copy families (Figure 3D; File S3). This suggests that abundant high-copy LTR retrotransposons may be more active than other LTR retrotransposons families in developing tung seeds. In addition, 184, 204, 244, 148, and 257 LTR retrotransposons exhibited the highest expression levels in root, stem, leaf, female flower, and male flower, respectively (File S3; Figure S22). Among these LTRs, high-copy LTR families also accounted for the highest proportion in the other five tissues.

### The tung tree eFP browser

A total of 28,422 genes were identified from the tung tree genome, of which 23,143 genes were annotated. The genome-wide gene identification allowed us to investigate gene expression on a large-scale in tung tree. To provide easy access and enable visualization of the expression levels of tung tree genes, flowers and seeds at different developmental stages were sampled for RNA-seq analysis (File S4). Based on RNA-seq data from 17 tung samples, a “Tung Tree eFP Browser” (at http://bar.utoronto.ca/efp_tung_tree/cgi-bin/efpWeb.cgi) was implemented to permit visualization of gene expression patterns with “absolute”, “relative” and “compare” modes in these tissues using the annotated gene IDs (File S4). The search interface generated an “electronic fluorescent pictograph” colored according to transcript abundance data for individual tung tree gene in various tissues/organs. As exemplified (Figure S23), the *VfFADx-1* (Vf11G0298) using linoleic acid (C18:2Δ9,12) as substrates to produce α-ESA (18:3Δ9,11,13) exhibited expression patterns consistent with its role in oil biosynthesis. In addition, the Tung Tree eFP Browser could be used for functional characterization of tung tree gene copies with different expression patterns. For instance, three feruloyl CoA ortho-hydroxylase homologues (Vf03G0652, Vf00G0634 and Vf03G0623) exhibited conservation of function as revealed by similar expression patterns in various tissues/organs (**Figure 4**; Table S51). Among three purple acid phosphatase homologues, the Vf11G0977 displayed neofunctionalization, *i.e*., functional diversification due to its expression in roots compared to the other homologues (Figure 4; Table S51).

**Figure 4.**
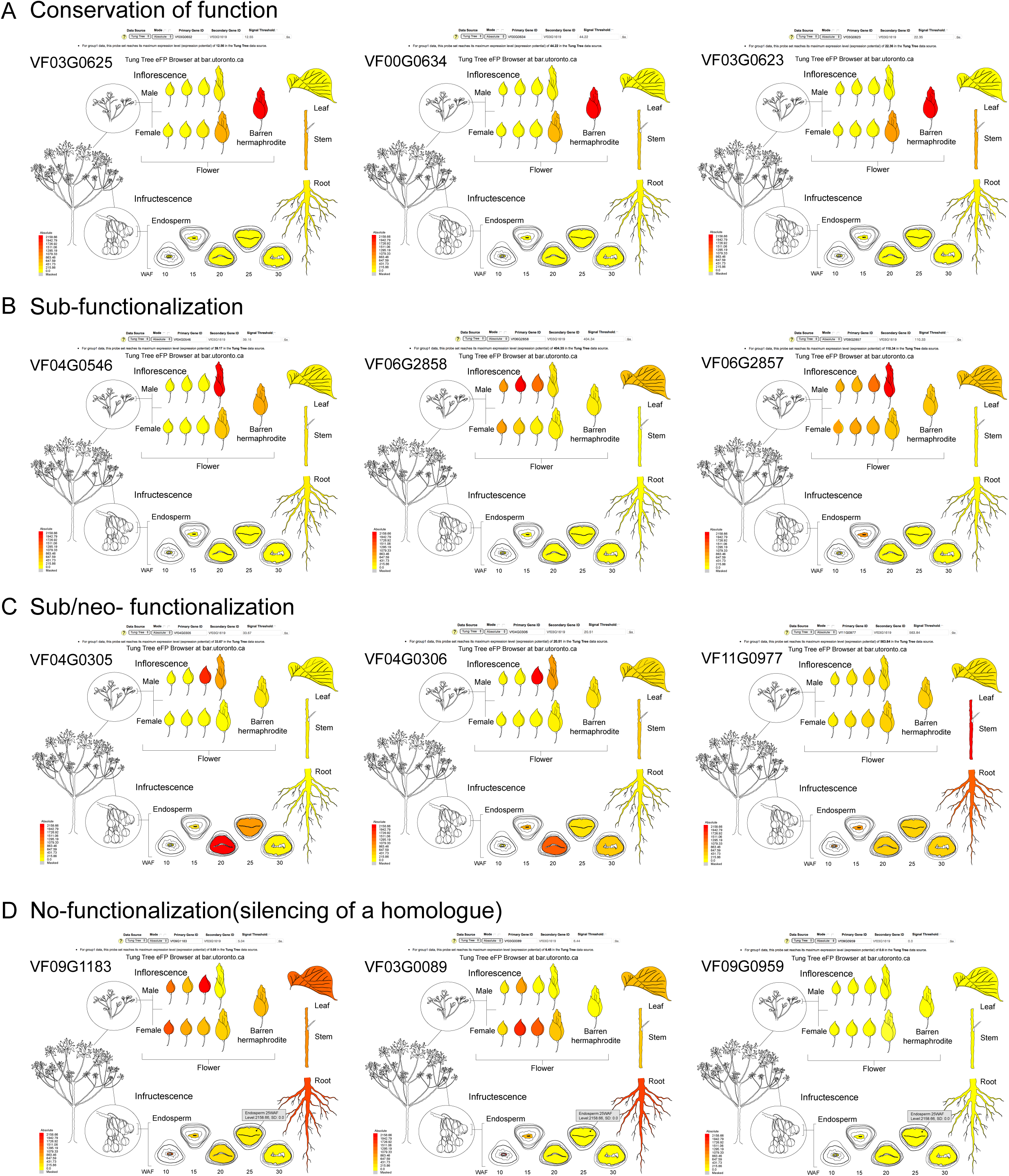
Functional conservation and diversification of tung tree homologs as visualized with the Tung Tree eFP Browser. eFP browser images showing conservation, sub-functionalization, neo-functionalization and non-functionalization of tung tree homologs. In each panel, the expression patterns of three homologs of each gene is shown. In all cases, red represents higher levels of transcript accumulation and yellow represents a lower level of transcript accumulation. From top to bottom, the genes are involved in feruloyl CoA ortho-hydroxylase (from left to right Vf03G0652, Vf00G0634, and Vf03G0623), Protein ECERIFERUM (from left to right Vf04G0546, Vf06G2858, and Vf06G2857), Purple acid phosphatase (from left to right Vf04G0305, Vf04G0306, and Vf11G0977), and Protein LYK5 (from left to right Vf09G1183, Vf03G0089, and Vf09G0959).WAF, week after flowering.

### NBS-encoding resistance genes

Disease resistance is one of the most important traits in tung tree breeding programs. The *V. fordii* is susceptible to wilt (*Fusarium oxysporum*), black spot (*Mycsphaqrella aleuritids*) and twig dieback (*Nectria aleuriidia*). Information on disease resistance-related genes will be helpful for understanding plant resistance mechanisms. Identification and characterization of these genes on a genome-wide scale will provide a basis for improvement of disease resistance in tung tree. Genes encoding nucleotide-binding sites (NBSs) are the largest class of plant disease resistance genes. Based on whether they contain a Toll/interleukin-1 receptor (TIR) domain, NBS resistance genes can be further categorized into two subclasses (TIR and non-TIR) (File S5).

A total of 88 genes with an NBS domain were identified in tung tree, of which 28 (31.82%) were organized in tandem arrays (Table S52; **Figure 5A**; Figure S25). The number of NBS-encoding genes in *V. fordii* was similar to *Z. mays* (107), but remarkably lower than *R. communis* (232), *M. esculenta* (312), *J. curcas* (275), and *H. brasiliensis* (483) (Table S52). The 88 NBS-encoding genes were classified into four subfamilies, including 23 coiled-coil (CC)-NBS, 16 NBS-leucine-rich repeat (LRR), 7 CC-NBS-LRR, and 42 NBS, however they did not form four independent classes in the phylogenetic tree (Figure 5A). Intriguingly, all of the tung tree NBS-encoding resistance genes do not belong to the TIR type (Table S52).

**Figure 5.**
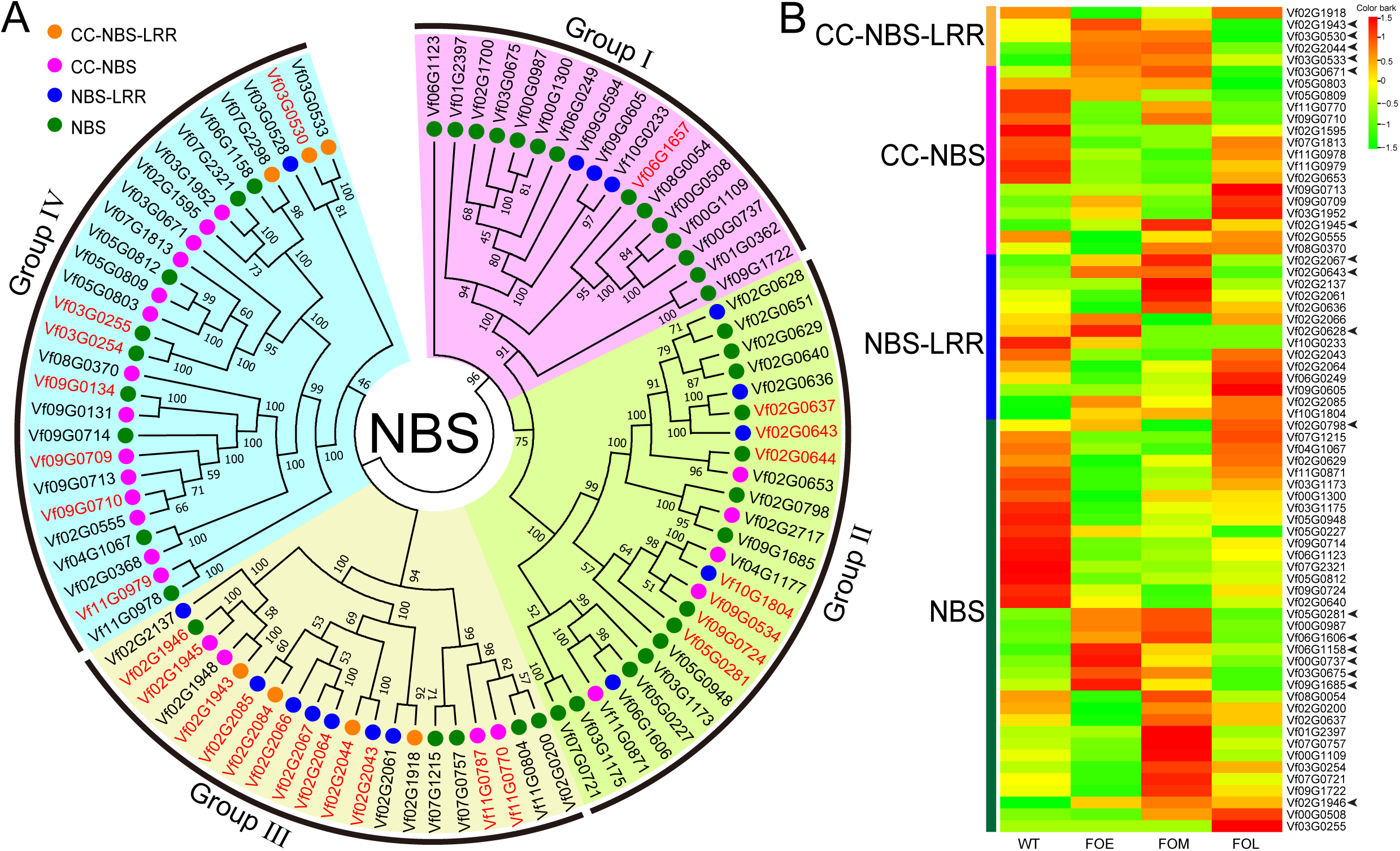
The NBS-encoding genes in tung tree genome. **A**. The maximum-likelihood phylogenetic tree based on 88 tung tree NBS encoding genes; dots in green, blue, pink, and orange indicate NBS subfamily, NBS-LRR subfamily, CC-NBS subfamily, and CC-NBS-LRR subfamily, respectively. Gene IDs in red indicate tandem repeats. **B**. Heat map of expression patterns of tung tree NBS-encoding genes. FOE, FOM, and FOL represents early, middle, and late stage after *F. oxysporum* infection. Different colored arrows indicate NBS genes responding to Fusarium wilt.

The NBS genes were distributed nonrandomly across all 11 chromosomes (Figure S24). More than 85% *NBS* genes were clustered in groups, and clusters were most abundant on chromosomes 2, 9, and 3 (Figure S24). Enrichment of *NBS* genes in these corresponding genomic regions indicated that resistance gene evolution might involve tandem duplication and divergence of linked gene families, as described in other plant genomes such as rubber tree [23] and pear [24]. RNA-seq data showed that the 88 tung tree NBS genes displayed differential expression patterns in roots after *F. oxysporum* infection (Figure 5B; File S5). The expression level of 17 genes including 8 *NBS*s, 3 *NBS-LRR*s, 2 *CC-NBS*s, and 4 *CC-NBS-LRR*s increased at early stage after infection (FOE) and decreased at late stage after infection (FOL) (Figure 5B). These results suggest that these genes may help the tree resist the pathogen shortly after infection.

### Evolution of genes involved in oil biosynthesis

Tung oil is the most important product from tung tree. Tung oil biosynthesis starting from acetyl-CoA involves 18 enzymatic steps with multiple isozymes in each step (**Figure 6A**). The oil is packed in subcellular structures called oil bodies or lipid droplets (Figure 6B; File S6). Tung seed oil droplets formed following the pattern of α-ESA accumulation in the seeds (Figure 6B and C). No visible oil droplet was observed in 10 weeks after flowering (WAF) seeds and small oil droplets were observed in 15 WAF seeds. The number and sizes of oil droplets were dramatically increased in more mature seeds (20, 25, and 30 WAF).

**Figure 6.**
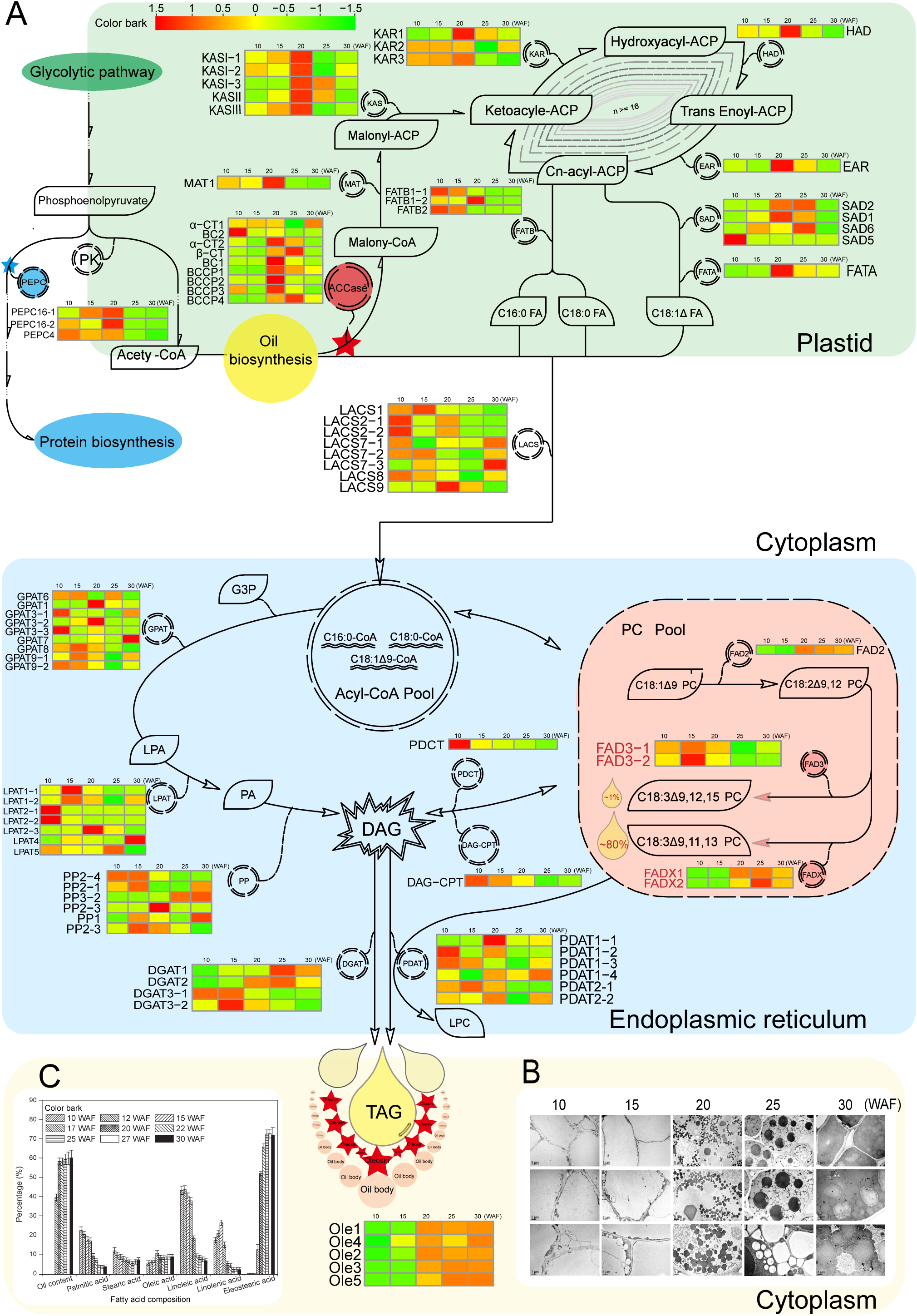
Network of genes involved in tung oil biosynthesis. **A**. Tung oil biosynthesis pathway. Tung oil biosynthesis is catalyzed by 18 enzymatic steps with multiple isozymes in each step. Acetyl-CoA is converted into C16 and C18 fatty acids in the plastid. TAG is synthesized in the endoplasmic reticulum and packed in the oil bodies. The metabolites are described in the black box. The enzymes are circled between two metabolite boxes. The expression levels of oil-biosynthesis genes are presented with the heat map. The scale bar of relative expression levels are shown at the top left. **B**. Oil droplet development in tung tree seeds. **C**. Tung oil and fatty acid accumulation profiles. PEPC, phosphoenolpyruvate carboxylase. PK, pyruvate kinase. ACCase, acetyl CoA carboxylase. α/β-CT, acetyl-coenzyme A carboxylase carboxyl transferase subunit alpha/ beta. BCCP, biotin carboxyl carrier protein. BC, biotin carboxylase. MAT, malonyl-CoA transacylases. KAS, ketoacyl-ACP synthase. KAR, ketoacyl-ACP reductase. HAD, hydroxyacyl-ACP dehydrase. EAR, enoyl-ACP reductase. FAT, fatty-acyl carrier protein thioesterase. SAD, stearoyl-ACP desaturase. FA, fatty acid. LACS, long-chain acyl-CoA synthetase. G3P, glycerol-3-phosphate. GPAT, glycerol-3-phosphate acyltransferase. LPA, lysophosphatidic acid. LPAT, lysophosphatidic acid acyltransferase. PA, phosphatidic acid. PP, phosphatidate Phosphatase DAG, diacylglycerol. PDCT, phosphatidylcholine. DAG-CPT, CDP-choline-diacylglycerol cholinephosphotransferase. PC, phosphatidylcholine. FAD, fatty-acid desaturase. DAGT, diacylglycerol O-acyltransferase. PDAT, phospholipid-DAG acyltransferase. LPC, lyso-phosphatidylcholine. TAG, triacylglycerol. Ole, oleosin. WAF, week after flowering.

Tung oil biosynthesis in the seeds started in mid-June (10 WAF), increased rapidly until 25 WAF with the oil content of 55.42% (Figure 6C), and ended by 30 WAF. Oleic acid (C18:1Δ9) accounted for minor percentage, whereas linoleic acid (C18:2Δ9,12) accounted for the major content (43%) in young seeds (10 and 15 WAF). Both gradually decreased in more mature seeds. Accumulation of linoleic acid and α-ESA (α-C18:3Δ9,11,13) showed opposite patterns in the developing tung seeds (Figure 6C) because linoleic acid is the same substrate for synthesizing α-ESA and α-ALA (α-linolenic acid, C18:3Δ9,12,15). The α-ESA synthesis started after 15 WAF and then increased rapidly up to 72.35% of seed oil following seed ripening (Figure 6C). The α-ALA accumulation was observed in 10 WAF seeds and accounted for minor percentage during the whole developmental stages, although it shares the same substrate with α-ESA. These developmental patterns of α-ESA biosynthesis and oil droplet formation were used as the criteria for selecting seed stages for our transcriptomic analysis.

We annotated 22,419 genes in the tung tree genome and identified 651 genes related to oil biosynthesis (Table S53). Among them, 88 genes were predicted more directly involved in oil biosynthesis (Figure 6A; File S7; Table S54). This study provided far more tung oil-related genes than those deposited in the GenBank databases (29 genes). These genes belonged to 18 families whose expression profiles were described in Figure 6A. The number of tung oil-related genes (88) was within the range of other plant species including 91 genes in *J. curcas*, 84 genes in *R. communis*, 87 genes in *A. thaliana*, 105 genes in *S. indicum*, and 210 genes in *G. Max* (Table S54).

Several key genes important in oil biosynthesis have been studied extensively, including acetyl CoA carboxylase (*ACCase*), *FAD*s, *DGAT*s and *oleosins* (*OLE*s) (Figure 6A). The current study indentified one additional *DGAT3* and two additional *FAD*s besides those reported previously. We also reported for the first time that tung tree genome had six *phospholipid:diacylglycerol acyltransferase* genes (*PDAT*) (Figure 6A).

ACCase and phosphoenolpyruvate carboxykinase (PEPC) are probably the key enzymes determining the metabolic pathways towards oil or protein biosynthesis in the seeds (Figure 6A) [25]. We identified nine *ACCase* genes in tung tree genome with high expression levels in the mid-late developing stages of tung seeds (Figure 6A). There are 10 *ACCase* genes in soybean, and 6-7 genes in other species (Table S54). We also identified three *PEPC* genes in tung tree genome which were expressed in the early developing stages of tung seeds (Figure 6A; Table S54). There are 16 *PEPC* genes in soybean and more *PEPC* genes in other species than tung tree. Comparison of soybean whose seed has high protein content (∼ 40%) and low oil content (∼ 20%), the fewer *PEPC* genes in tung tree genome might be the reason of high oil (∼ 55%) and low protein content (∼ 5%) in tung seeds, probably contributing to carbon flow towards fatty acid biosynthesis in tung seeds.

FAD protein family catalyzes the desaturation of fatty acids [5] and therefore is responsible for polyunsaturated lipid synthesis in developing seeds of oil crops. FAD2 and FAD3 are the main enzymes responsible for the Δ12 linoleic acid and Δ15 linolenic acid desaturation, respectively. We identified one *FAD2*, two *FAD3* and two *FADx* genes in tung tree (Table S54). *FAD2* and *FADx-1* were highly expressed in mid-late stages of developing seeds; whereas *FAD3* was expressed higher in early stages of seeds (Figure 6A). FADx, a divergent FAD2, converts linoleic acid to α-ESA, the major component of tung oil [14], but the evolutionary relationship between FADx and FAD2 is still uncertain. According to the newly generated phylogenetic tree in this study (**Figure 7**), we found FAD2/x clade could be divided into two clades (FAD2 and FADx) in eudicot plants, suggesting that the two clades were due to gene duplication in eudicot ancestors. The eudicot ancestors have γ WGD event, and gene duplication is likely to be retained by the WGD event. Further synteny analysis revealed that FAD2s and FADxs were likely to be generated by WGDs event (Table S56), which corresponded to the γ WGD event shared by core eudicots. Notably, the FADx clade lost many genes in species like the members of Brassicaceae.

**Figure 7.**
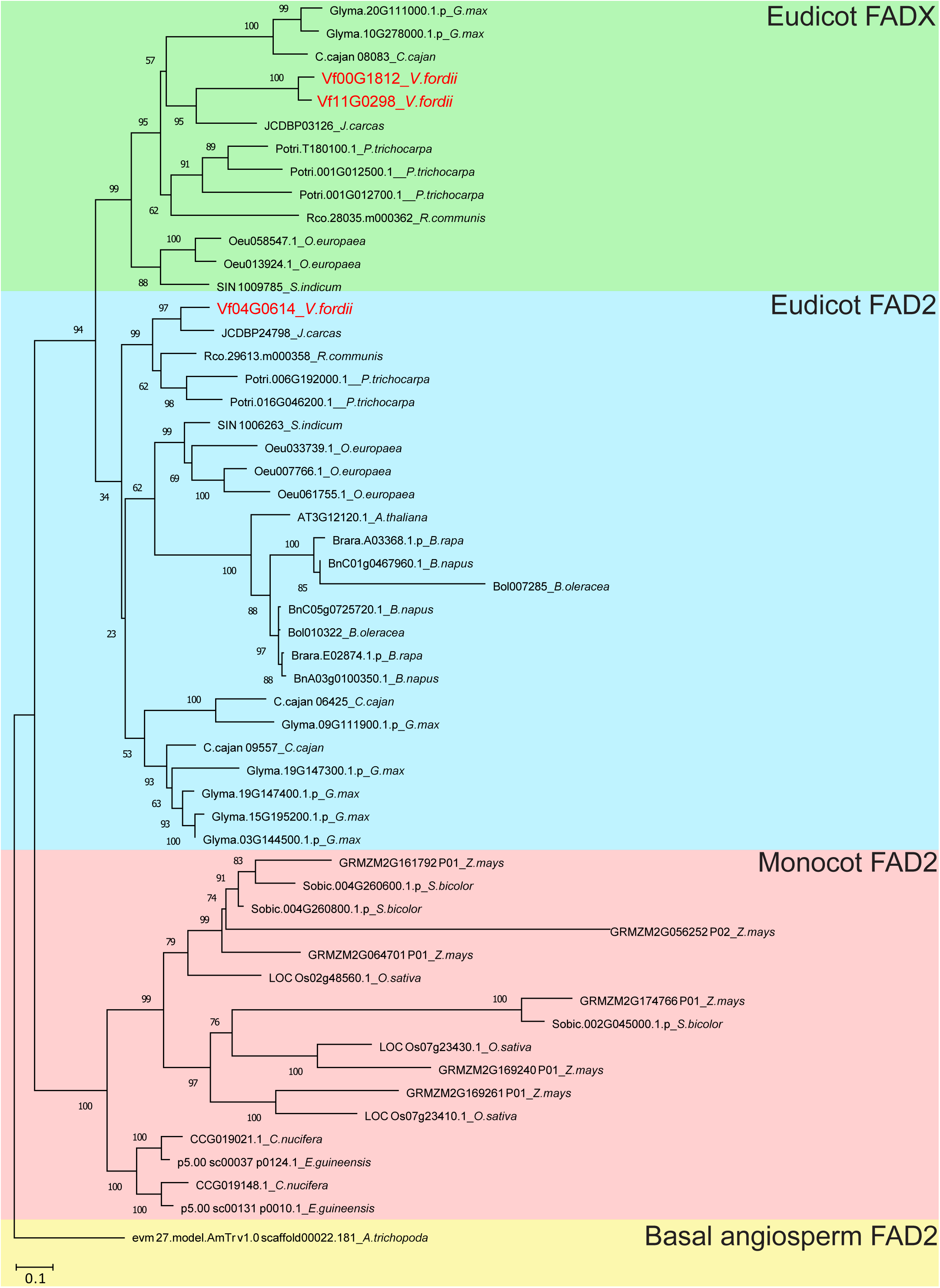
Phylogeny of FAD2 and FADx proteins. A maximum-likelihood phylogenetic tree constructed from protein sequences. The taxon names in the phylogenetic tree are indicated after gene ID. The clades are marked by four different block colors in the tree. The last one (yellow), a basal angiosperm, *A. trichopoda*, used as an outgroup; the monocot FAD2, eudicot FAD2 and eudicot FADx clades are marked in red, blue and green, respectively.

DGAT protein family catalyzes the last step of triacylglycerol (TAG) biosynthesis and is regarded as the rate-limiting step for TAG accumulation. Three DGATs were reported in tung tree in previous studies. *DGAT2* was proposed to be the most important *DGAT* for TAG biosynthesis in tung tree seeds. Our transcripomics study found four *DGAT*s (*DGAT1*, *DGAT2*, and two *DGAT3*) expressed in tung seeds (Figure 6A; Table S55). *DGAT2* was confirmed to be the most abundantly expressed *DGAT* in tung seeds which corresponded to oil accumulation (20–30 WAF), but *DGAT3-1* was the dominant form of *DGAT* in immature seeds (10–15 WAF) and other tissues including stem, root, leaf and female flower (Figure 6A; Table S55).

Recently, it has become obvious that TAG synthesis also can be catalyzed by PDAT. We reported for the first time that there were five PDATs in tung tree genome. *PDAT1-1* and *1-4* were expressed more in mid-late stages of developing seeds but the other three *PDAT* genes were expressed more in the early stages of developing seeds (Figure 6A).

OLEs are the major proteins in plant oil bodies. Genome-wide phylogenetic analysis and multiple sequence alignment demonstrated that the five tung *OLE* genes represented the five OLE subfamilies and all contained the “proline knot” motif (PX5SPX3P) shared among 65 OLE from 19 tree species [26]. We confirmed the five tung tree *OLE* genes coding for small hydrophobic proteins. These five *OLE*s were highly expressed in mid-late stage of developing tung seeds (Figure 6A; Table S55).

A total of eight *long chain fatty aycl-CoA synthetases* (*LACS*) genes were identified in tung tree genome, of which *LACS1* and *2* were more highly expressed in early stage but *LACS7*, *8* and *9* were highly expressed in mid-late stages of developing seeds (Figure 6A). Additionally, 9 *glycerol-3-phosphateacyltransferases* (*GPAT*s), 7 *lysophosphatidic acid acyltransferases* (*LPAT*s), and 6 *phosphatidate phosphatases* (*PP*s) genes were identified in tung tree genome whose expression levels of some genes were higher in early stage rather than late stages of developing seeds and verse visa (Figure 6A; Table S55).

To explore possible synergistic effects among genes in oil accumulation, we performed a weighted correlation network analysis of transcript expression in developing seeds at five stages (FPKM values ≥ 1) (File S8). We identified 10 co-expression modules for each stage sample, among which oil biosynthesis-related genes at 20 WAF were highly enriched in two significant modules (PCC values ≥ 0.8, *P* value ≤ 0.1): MEbrown and MEyellow containing 1,156 and 908 genes, respectively (Tables S57 and S58; Figures S31 and S32). We did not find oil biosynthesis-related genes in other significant modules. In MEyellow and MEbrown modules, 18 and 13 genes were respectively identified to play pivotal roles in fatty acid synthesis and oil accumulation, such as fatty acid synthases (FASs), the upstream rate-limiting enzyme *ACCase* subunits (*α*-*CT*, *BCCP-1*, *BCCP-2*, *BCCP-2*, and *BC-1*), and genes related to TAG assembly like *GPDH*, *LPAT,* etc (**Figure 8**). A number of transcription factors were also identified in the two modules and co-expression networks (Figure 8) including *WRINKLED1* (*WRI1*), *FUSCA3* (*FUS3*), *LEAFY COTYLEDON1* (*LEC1*), and *ABSCISIC Acid INSENSITIVE3* (*ABI3*), which has been reported to facilitate oil accumulation by interacting each other or with oil biosynthesis-related genes [27–31]. We selected four tung tree transcription factors (*FUS3*, *ABI3*, *LEC1-1* and *LEC1-2*) to conduct yeast two-hybrid assay (File S9) and observed that *FUS3* and *LEC1-2* were interacted (Figure S33). The gene co-expression networks indicate that transcription factors and oil biosynthesis-related genes have synergistic effects in oil biosynthesis, which may contribute to high oil content in tung seeds.

**Figure 8.**
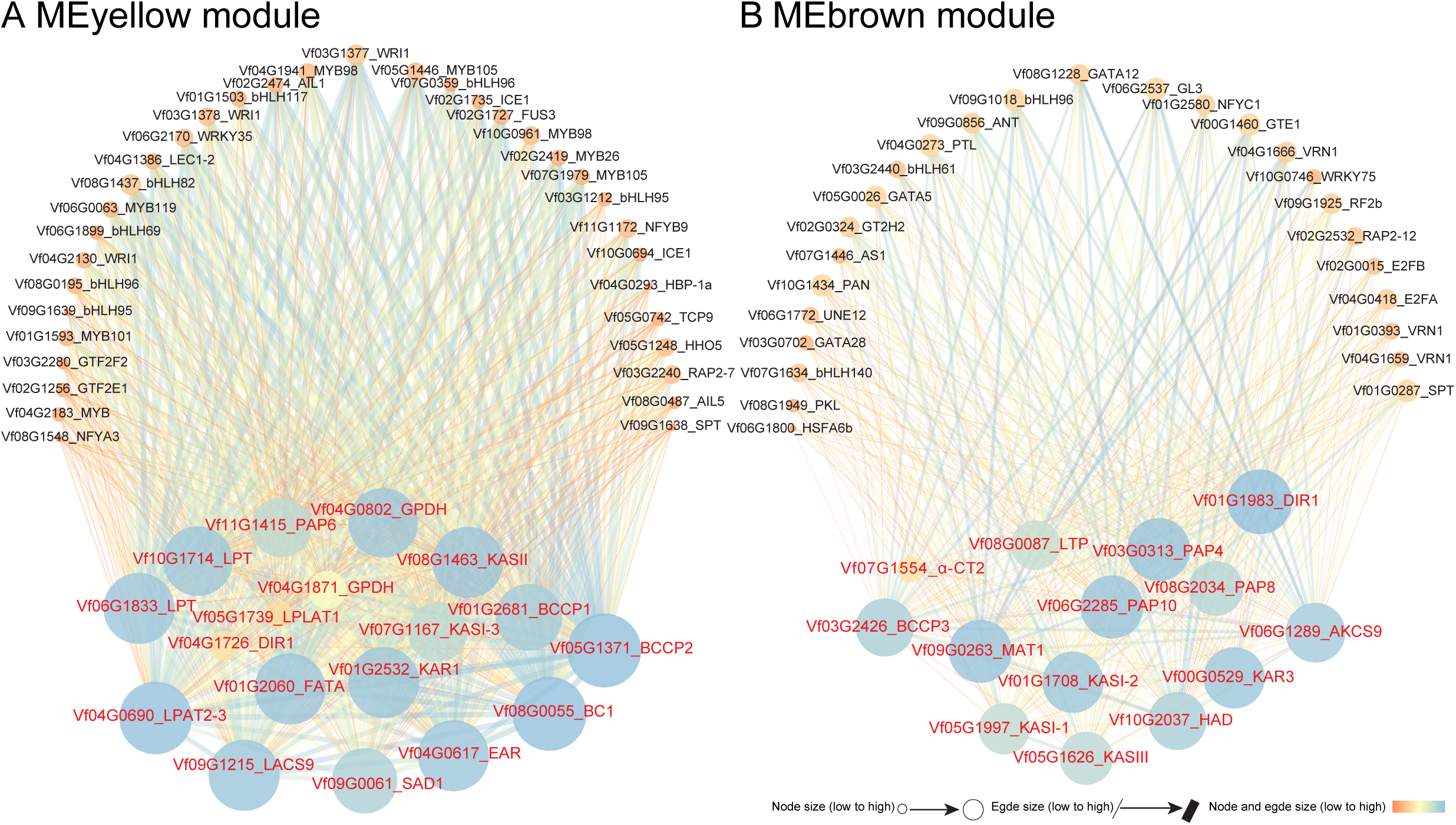
Co-expression networks of tung tree oil biosynthesis-related genes and transcription factors at the transcriptome level. Oil biosynthesis-related genes are colored in red, and their adjacent transcription factors are colored in black.

## Discussion

The whole genomes of an increasing number of plant species have been sequenced due to rapid development of new sequencing technologies in recent years. The genome information provides researchers a useful resource for better understanding plant evolutionary history and exploring important genes to uncover the mechanisms controlling various traits during long-term evolution process. As an economically important tree species, tung tree has been cultivated and utilized for thousands of years. Presently its oil has a great potential for producing environmentally-friendly coatings with low VOCs. However, producing tung oil on an industrial scale is hampered by low yield. Our genome sequencing effort will facilitate the breeding of elite cultivars with yield-related traits including fruit setting rate and seed oil content. In this study, the large amount of repeat sequences and low GC content made the tung tree genome a challenge for WGS strategies using NGS technology even though the tung tree genome was estimated to be extremely low heterozygosity. To overcome the challenge of high repeat content, we generated long reads from 10 kb and 20 kb libraries via PacBio sequencing. Finally, we used the Hi-C map to generate a chromosome-scale assembly of the tung tree genome. The genome sequence covered ∼ 85.50% of the estimated genome size and harbored 28,422 genes. Among the Euphorbiaceae family, rubber tree and cassava instead of tung tree, physic nut and castor bean were found to have undergone a recent WGD event, although they all shared an ancient WGD event. Interestingly, rubber tree and cassava have more genes than the other three species (Figure 2A). The recent WGD event could cause chromosomal rearrangements, fissions or fusions and is one of the reasons resulting in expansion of gene families [18], which may contribute to more gene expansions in rubber tree and cassava than those in tung tree, physic nut and castor bean. The genome sequence of tung tree opens a window to functional and molecular breeding of economically important woody oil plants within the Euphorbiaceae family.

Tung tree had a larger genome size than physic nut and castor bean. In most cases, genome expansions are caused by repeated sequence insertion, like those occurred in tea tree, rubber tree, and Ginkgo (*G. biloba*) [32]. Similar to the three species, *Ty3/Gypsy* families contributed the most to the tung tree genome expansion. Based on our insertion time analysis, we proposed that lack of efficient deleting mechanisms of repeated DNA sequences might have resulted in long-term and continuous LTR retrotransposon bursts and growth, eventually leading to the whole genome size expansion. This is also consistent with the findings in tea tree and *P. abies* [33]. We also found that different LTR retrotransposon families were differentially expressed in various tissues, confirming the retrotransposon activity in the tung tree genome. The eFP Browser has proved to be a useful tool to display gene expression levels visually in several plant species including *A. thaliana*, *P. trichocarpa*, *G. Max*, *S. tuberosum*, *S. lycopersicum*, *C. sativa*, *F. vesca* and other species [34–37]. Based on tung tree genome sequences generated in this study, we created a Tung Tree eFP Browser to display tung tree RNA-seq data from 17 different tissues and stages. This eFP Browser work should facilitate further research in tung tree and other Euphorbiaceae plants.

Plant disease resistance has always been a research hotspot. NBS genes are the largest class of plant disease resistance genes. They confer the capacity for the plant to resist the intrusion of outside pathogens, including bacteria, fungi and virus [38]. The present studies suggested that the TIR domain-containing NBS genes are widely distributed in dicots but not monocots, whereas they are lost in tung tree genome. To date only tung tree and sesame [22] out of dicots have been reported for TIR domain-containing NBS gene loss. This finding provides a new paradigm to investigate the evolution of disease resistance genes. CC is the functional domain of many proteins and CC structure plays an important role in protein-protein interaction [39]. LRR is the signal region in transmembrane domain and loss of it can result in loss of function [40]. In this study, the highest proportion of *CC-NBS-LRR* genes (4/7, 57.14%) responded to *F. oxysporum* infection at early stage, suggesting that CC and LRR domains play more important roles than other domains.

Tung tree is a high-efficient photosynthetic tree with strong photosynthesis rate. Sucrose, the major photosynthesis product, is synthesized in the chloroplast and exported to the sink tissues such as seeds for seed development and metabolite accumulation. Sucrose is converted into hexose phosphate, triose phosphate, phosphoenolpyruvate (PEP), and pyruvate. PEP is a key intermediate metabolite for synthesizing both fatty acids and proteins. PEP is converted into pyruate by pyruate kinase (PK), which is subsequently converted into acyl-CoA and enters fatty acid biosynthesis pathway after ACCase action. On the other hand, PEP is catalyzed by PEPC to produce oxaloacetic acid, which is subsequently used for protein synthesis. Therefore, ACCase and PEPC are probably the keys enzymes determining the metabolic pathway towards oil or protein biosynthesis in the seeds [25]. We identified nine *ACCase* genes in tung tree genome with high expression levels in the mid-late developing stages of tung seeds, which are indicative of their importance in tung oil biosynthesis. There are 10 *ACCase* genes in soybean, and 6-7 genes in other species. We also identified three *PEPC* genes in tung tree genome with high expression levels in the early developing stages of tung seeds. By contrast, there are 16 *PEPC* genes in soybean and more *PEPC* genes in other species than tung tree. Because soybean has more *PEPC* genes and higher protein/ lower oil content in the seed, it is possible that the fewer *PEPC* genes in tung tree diverted less carbon flow towards protein biosynthesis and resulted in high oil/low protein content in tung seeds.

Tung oil is the major economically important product from tung tree. Identification and characterization of all genes involved in tung oil biosynthesis is essential for improving tung oil production and economic value. Many tung oil biosynthetic genes have been identified in our laboratories, including those coding for *diacylglycerol acyltransferases* (*DGAT*) [13, 17], *delta-12 oleic acid desaturase* (*FAD2*) and *delta-12 fatty acid conjugase* (*FADx*) [14], *omega-3 fatty acid desaturase* (*FAD3*) [41], *acyl-CoA binding proteins* [42], *cytochrome b5* [43], *cytochrome b5 reductase* [15], *glycerol-3-phosphate acyltransferase* (*GPAT*) [44], *plastid-type omega-3 fatty acid desaturase* (*TnDES2*) [45], *aquaporin* and *glutaredoxin* [46], and *ß-ketoacyl-ACP synthase* (*KAS*) [47]. Interestingly, we identified an additional *FADx* gene, *FADx-2*, which was probably generated by gene duplication and sub-functionalization based on the different expression patterns of *FADx-1* and *FADx-2* genes. In comparison with *FADx-2*, *FADx-1* was the dominant form responsible for α-ESA synthesis in developing seeds of tung tree. We also identified 9 *ACCases*, 4 *DGAT*s, 7 *FAD*s, 6 *PDAT*s, 5 *OLE*s, 8 *LACS*s, 9 *GPAT*s, 7 *LPAT*s, and 6 *PP*s genes in the tung tree genome. This study provided a more complete picture for genes involved in tung oil biosynthesis. The numbers of tung oil-synthesizing genes are within the range of other species. These suggest that there is no gene expansion in tung tree and the amount and types of oils in various species may not be directly related to the number of genes in oil biosynthesis.

Transcriptomic analysis evaluated the expression profiles of all these genes. Our results indicated that the expression patterns of some of the most important genes were well-coordinated with oil biosynthesis and accumulation in tung tree seeds. Specifically, *DGAT2* was shown to be the most abundantly expressed *DGAT* in tung seeds but *DGAT3-1* was the dominant form of *DGAT* in immature seeds and other tissues including stem, root, leaf and female flower, in agreement with our previous results [13, 17]. *FAD2* and *FADx* were highly expressed in mid-late stages of developing seeds; whereas *FAD3* was expressed higher in early stages of seed, also in agreement with published results [14]. All five *OLE*s were highly expressed in mid-late stage of developing tung seeds, similar results to what we reported previously [26]. Our expression analysis provided novel insights into the potential role of *PDAT*s in tung oil biosynthesis by showing that *PDAT1-1*, *1-4*, and *2-2* were expressed more in mid-late stages of developing seeds but the other three *PDAT* genes were expressed more in the early stages of developing seeds, which were not reported previously. Our gene co-expression analysis revealed that oil biosynthesis-related genes were enriched in two significant modules only at 20 WAF when the seed oil started to accumulate rapidly. The enriched oil biosynthesis-related genes included most of FAS genes, part of TAG biosynthesis genes and some transcription factors. The complete gene co-expression networks provide insights into oil biosynthesis by gene-gene synergistic function.

In conclusion, this study provides whole-genome sequence information, eFP browser, and extensive RNA-seq data. These critical lines of information should be used as valuable resources for functional genomics studies and tree improvement of economically important traits such as oil content and disease resistance in the tung tree.

## Materials and methods

### Plant materials

The self-bred progeny ‘VF1-12’ of the elite *V. fordii* cv. Putaotong was used for whole genome sequencing in this study (File S1). Young leaves were collected from ‘VF1-12’ in the spring for genome sequencing. Young plantlets were used for Hi-C library construction and sequencing. A total of 17 fresh tissues including stems, roots, male flowers, female flowers, and seeds at different developmental stages were collected for RNA-seq. The developing seeds were also used for oil content measurement and fatty acid analysis.

### Whole-genome sequencing, assembly and assessment

The tung tree genome size was estimated by a modified Lander-Waterman algorithm *i.e.*, a formula G = Bnum/Bdepth = Knum/Kdepth [48]. Heterozygosity was estimated by the k-mer distribution and GenomeScope [49]. Nuclear DNA was isolated from fresh leaf tissues by using a DNeasy Plant Mini Kit (Qiagen, CA, USA). A series of DNA libraries were constructed and sequenced with an Illumina HiSeq 2000 sequencing platform (Illumina, CA, USA) (File S10). In addition, SMRTbell template libraries of 20 kb were constructed and sequenced on the PacBio RSII. After removing low-quality reads, the whole genome assembly of tung tree was performed with a hierarchical assembly strategy due to its homozygous genome with highly repetitive sequences (File S11). The genome completeness was assessed by Core Eukaryotic Genes Mapping Approach (CEGMA) [50], Benchmarking Universal Single-Copy Orthologs (BUSCO) analysis [51] and RNA-seq reads mapping [52].

### Hi-C data preparation and contig clustering

The Hi-C library was prepared with the standard procedure described [53]. Raw Hi-C data were generated using HiSeq2500 sequencing platform and then were processed to filter low-quality reads and trim adapters. Clean reads were mapped to the assembled scaffolds by BWA-aln after truncating the putative Hi-C junctions in sequence reads. HiC-Pro software (version 2.7.1) was used to filter the invalid ligation read pairs, including dangling-end and self-ligation, re-ligation and dumped products. Finally the scaffolds were clustered, ordered and orientated onto chromosomes using the valid read pairs by LACHESIS (http://shendurelab.github.io/LACHESIS/).

### Genome annotation

Gene prediction was conducted using *de novo* prediction, homology information and RNA-seq data (File S12). Gene functions were assigned according to the best match derived from the alignments to proteins annotated in SwissProt and TrEMBL databases using Blastp, and the pathway in which the gene might be involved was annotated by KAAS [54]. Motifs and domains were annotated using Inter ProScan (Version 5.2-45.0) [55] by searching against publicly available databases in InterPro [56]. The rRNA, snRNA and miRNA genes were predicted by INFERNAL software using the Rfam database. The rRNA subunits were identified by RNAmmer [57] based on hidden Markkov models (HMMs). The tRNA genes were predicted by tRNAscan-SE [58] with eukaryote parameters. A *de novo* and homology-based approach was used to identify repetitive sequence and transposable elements (TEs) in the tung tree genome.

### Evolutionary analysis

Phylogeny of a total of eight species was constructed based on single-copy gene families by the maximum likelihood (ML) method (File S13). The divergence times were estimated based on all single-copy genes and 4-fold degenerate sites with the program MCMCTree of the PAML package [59]. The neutral evolutionary rate was calculated via Bayes estimation with Markov Chain Monte Carlo algorithm. Gene families which underwent expansions or contractions were identified using the CAFE (Computational Analysis of gene Family Evolution) program [60].The selection pressure of tung tree in the phylogenetic tree was calculated by Codeml. The significance of the identified PSGs was verified using a Chi-square test. WGD events were identified by 4DTv (four-fold synonymous third-codon transversion) and synonymous Ks analysis.

### Data access

The project of tung tree genome sequencing, Hi-C and transcriptome sequencing is registered at NCBI under BioProject accession PRJNA503685 (http://www.ncbi.nlm.nih.gov/bioproject/503685), PRJNA445350 (http://www.ncbi.nlm.nih.gov/bioproject/445350) and PRJNA483508 (http://www.ncbi.nlm.nih.gov/bioproject/483508). The data are publicly available at NCBI under accession number SUB4731026, SRP136294 and SRP155790, respectively.

## Author contributions

XFT and LZ conceived and initiated the study. XFT, LZ, and HPC designed experiments and coordinated the project. XFT, LZ, HXL, MLL, ZL, YLZ, and HC performed the field controlled pollination and managing and sampling experimental materials for genome and transcriptome sequencing. JH and DPW supervised the data generation and analysis. MZ, JJL, FL, JH, and DPW conducted the genome assembly and annotation. MZ and MLX were involved in the WGD event determination. XMY, MZ, and LSZ performed repeat annotation, phylogenetic analysis and expression analysis. AP, EE, WYL, and NJP performed tung tree eFP browser construction. WD, AXS, and LSZ coordinated collinear analysis and phylogenetic analysis. MLL, DR, and GZ performed NBS gene family analysis. HXL, MLL, LZ, and HPC contributed to identification, expression and phylogenetic analysis of oil-related gene families. WD and MLL conducted gene co-expression analysis. MLL performed Yeast two-hybrid assay. WYL and MLL drew all figures of the manuscript. LZ and HPC wrote the manuscript. LZ, HPC, NJP, LSZ, and XFT revised the manuscript. All authors discussed results and commented on the manuscript.

## Acknowledgments

This work was supported by the National Key R&D Program of China (Grant No. 2017YFD0600703), the National Natural Science Foundation of China (Grant No. 31770720), the outstanding youth of the Education Department of Hunan Province (Grant No. 17B279), and the USDA-ARS Quality and Utilization of Agricultural Products National Program 306 through CRIS 6054-41000-103-00-D. Mention of trade names or commercial products in this publication is solely for the purpose of providing specific information and does not imply recommendation or endorsement by the U.S. Department of Agriculture. USDA is an equal opportunity provider and employer.

## Tables

**Table 1 Statistics of tung tree genome assembly and annotation**

## Supplementary material

**File S1 Self-pollination and heterozygosity estimation**

**File S2 Estimation of genome size and heterozygosity**

**File S3 Repeat sequence analysis**

**File S4 Transcriptome sequencing, assembly and eFP browser**

**File S5 Identification and expression of NBS-encoding gene families**

**File S6 Lipid analysis and electron microscopy observation**

**File S7 Oil biosynthesis-related gene family identification and phyleogenetic analysis**

**File S8 Gene co-expression analysis**

**File S9 Yeast two-hybrid assay of transcription factors**

**File S10 Whole-genome shotgun sequencing**

**File S11 Genome assembly and assessment**

**File S12 Gene prediction and functional annotation**

**File S13 Evolutionary analysis**

**Figure S4 The k-mer analysis to estimate the tung tree genome size**

**Figure S5 Distribution of length (A) and quality (B) of Pacbio raw reads**

**Figure S6 Distribution of inserted fragment length for Hi-C library**

**Figure S7 Hi-C linkage density heat map of assembled contigs**

**Figure S8 Cross-species comparison of gene elements between tung tree and other five species**

**Figure S9 Gene GO classification of tung tree genome**

**Figure S10 GO classification of tung tree-specific gene families**

**Figure S11 Venn diagrams of cross-species gene family comparisons**

**Figure S12 Phylogenetic tree of tung tree and seven other plant species**

**Figure S13 GO classification of tung tree expanded gene families**

**Figure S14 GO classification of tung tree PSGs**

**Figure S15 Synteny analysis between *V. fordii* and *M. esculenta***

**Figure S16 Synteny analysis between *V. fordii* and *V. vinifera***

**Figure S17 Collinear relationship of *V. fordii*, *M. esculenta* and *V. vinifera***

**Figure S18 Evolutionary history of TE super-families in tung tree genome**

**Figure S19 The insertion times for intact LTR retrotransposons in tung tree genome**

**Figure S20 Insertion times of *Ty1/Copia*, *Ty3/Gypsy* and other LTR retrotransposon families in tung tree genome**

**Figure S21 Insertion times of various copy number of retrotransposon families in tung tree genome**

**Figure S23 eFP browser view of gene expression pattern in tung tree**

**Figure S24 Chromosomal locations and region duplication for tung tree NBS genes**

**Figure S25 Phylogenetic analysis of NBS-encoding genes**

**Figure S31 The relationship between co-expression module and trait in tung tree**

**Figure S32 Co-expression network analysis of genes in developing seeds in tung tree**

**Figure S33 Yeast two-hybrid assay of transcription factors**

**Table S2 Sequencing data for 500 kb-library used in genome survey**

**Table S3 Sequencing data for 17-mer analysis**

**Table S4 Statistics of sequencing data for tung tree genome**

**Table S5 Statistics of the tung tree genome assembly**

**Table S6 Sequencing data for Hi-C library**

**Table S7 Mapped reads of Hi-C library to tung tree genome**

**Table S8 Statistics of Hi-C sequencing data**

**Table S9 Efficient coverage of Hi-C data to tung tree genome assembly**

**Table S10 Data of tung tree genome after Hi-C assembly**

**Table S11 Scaffold information after Hi-C assembly**

**Table S12 Assessment for completeness of tung tree genome by CEGMA**

**Table S13 Assessment for completeness of tung tree genome by BUSCO**

**Table S14 Coverage of tung tree genome from male flower unigenes**

**Table S15 Coverage of tung tree genome from female flower unigenes**

**Table S16 Coverage of tung tree genome from seed 1 unigenes**

**Table S17 Coverage bof tung tree genome from seed 2 unigenes**

**Table S18 Coverage of tung tree genome from seed 3 unigenes**

**Table S19 Coverage of tung tree genome assembly from merged seed unigenes**

**Table S20 Transcriptomic reads mapped to tung tree genome**

**Table S21 Comparison of gene modules between tung tree and other species**

**Table S22 The GC content across the tung tree genome**

**Table S23 The GC content in coding sequences in the tung tree genome**

**Table S24 The GC content in intron regions in the tung tree genome**

**Table S25 Assessment for completeness of predicted tung tree genes by BUSCO**

**Table S26 Gene functional annotation of tung tree genome**

**Table S27 Pathways based on KEGG annotation in tung tree genome**

**Table S28 Non-coding RNAs in tung tree genome**

**Table S29 Statistics of gene families of *V. forrdi* and other 7 species**

**Table S30 GO analysis of unique gene families in the tung tree genome**

**Table S31 KEGG analysis of unique gene families in the tung tree genome.**

**Table S32 GO analysis of expanded gene families in tung tree genome**

**Table S33 Swissprot annotation of PSGs in tung tree genome**

**Table S34 GO analysis of PSGs in tung tree genome**

**Table S35 Collinearity analysis in tung tree genome**

**Table S36 Collinearity analysis in the *M.esculenta* genome**

**Table S37 Collinearity analysis in the *V. vinifera* genome**

**Table S40 Summary of repeat sequences in tung tree genome**

**Table S41 Annotation of repeat sequences in tung tree genome**

**Table S51 Four types of functional conservation and diversification of tung tree homologs**

**Table S52 Cross-species comparison of NBS-encoding gene family number**

**Table S53 Total oil-related gene families in tung tree genome**

**Table S54 Cross-species comparison of oil-related gene family number**

**Table S55 Expression quantity (FPKM value) and duplication type of 88 important oil genes in tung tree genome**

**Table S56 Colinear gene pairs in poplar (*P. trichocarpa*)**

**Table S57 Co-expression relationship among oil-related genes and transcription factors in yellow module**

**Table S58 Co-expression relationship among oil-related genes and transcription factors in brown module**

