## Supplementary material for "The Tung Tree (*Vernicia Fordii*) Genome Provides A Resource for Understanding Genome Evolution and Oil Improvement": File S1

**File S1: Self-pollination and heterozygosity estimation**

To obtain plants with low heterozygosity for whole genome sequencing, we conducted self-pollination of an elite tung tree (*Vernicia fordii*) cv. Putaotong (Figure S2) at Central South University of Forestry and Technology Germplasm Repository (Yongshun, Hunan). A five-year-old tree was used for control pollination in April, 2012 and a total of 258 seeds were obtained and finally 61 seedlings were produced (Figure S3). The 61 self-bred progenies of ‘Putaotong’ were then used for heterozygosity estimation by 27 SSR markers including 20 genomic SSR markers from Xu et al [1] and 7 EST-SSR markers from Jia et al [2] (Table S1). Consequently, 15 progenies (‘VFPPT1-1’, ‘VFPPT1-4’, ‘VFPPT1-5’, ‘VFPPT1-9’, ‘VFPPT1-12’, ‘VFPPT2-16’, ‘VFPPT2-17’, ‘VFPPT3-2’, ‘VFPPT3-3’, ‘VFPPT4-3’, ‘VFPPT4-6’, ‘VFPPT5-3’, ‘VFPPT5-4’, ‘VFPPT5-5’and ‘VFPPT5-7’) were found to harbor most homozygous loci (25 in each plant) and preliminarily estimated to be low heterozygosity. The ‘VF1-12’ was used for genome survey and whole genome sequencing.

**Supplementary Figures and Tables**

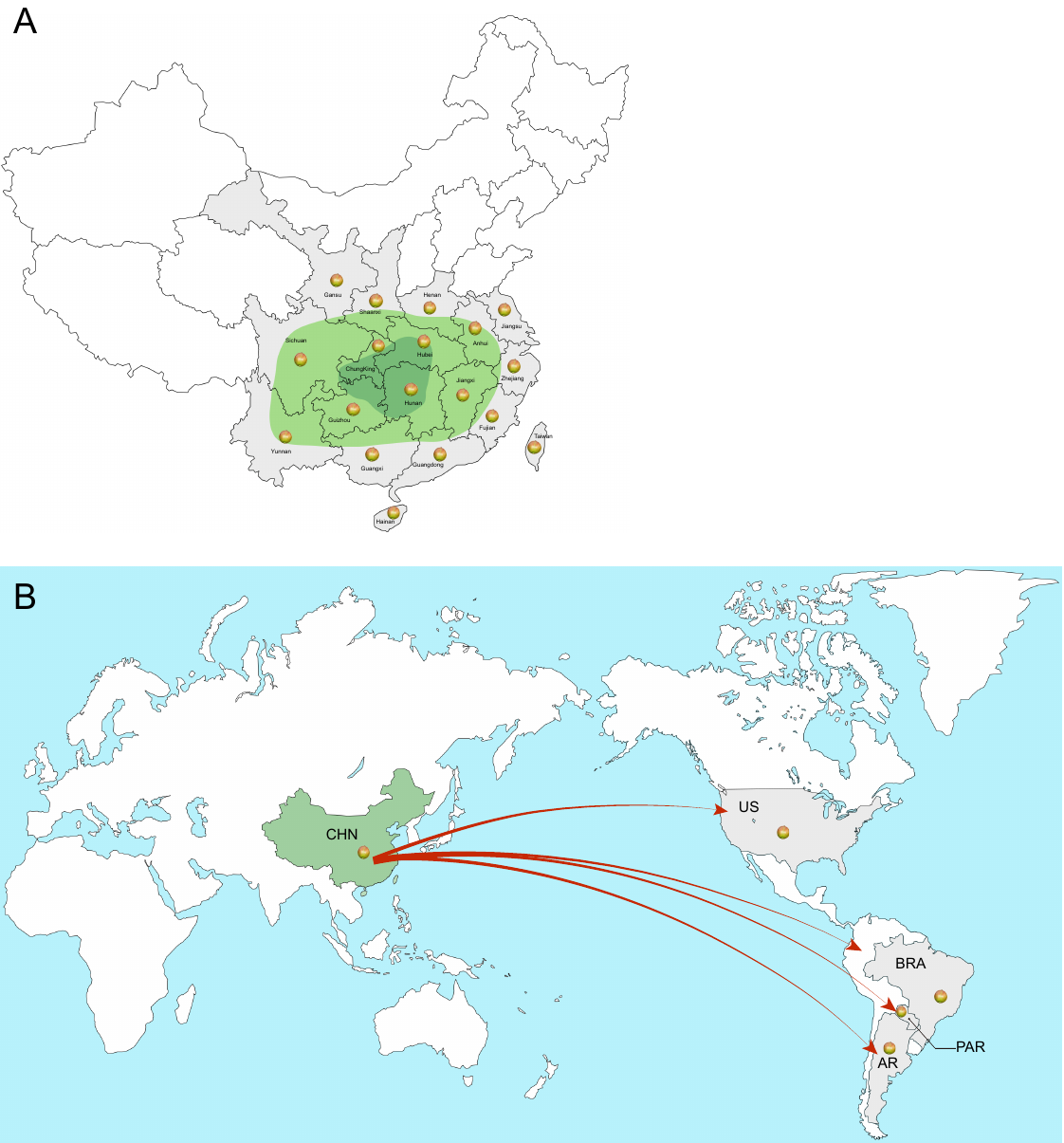

**Figure S1. Distribution of tung tree (*V. fordii*) in China (A) and America (B).** A: Dark green, core production areas; Green, main production areas; Gray, cultivated areas. B: Arrow indicates the route of tung tree introduction to the US, Brazil, Paraguay and Argentina.

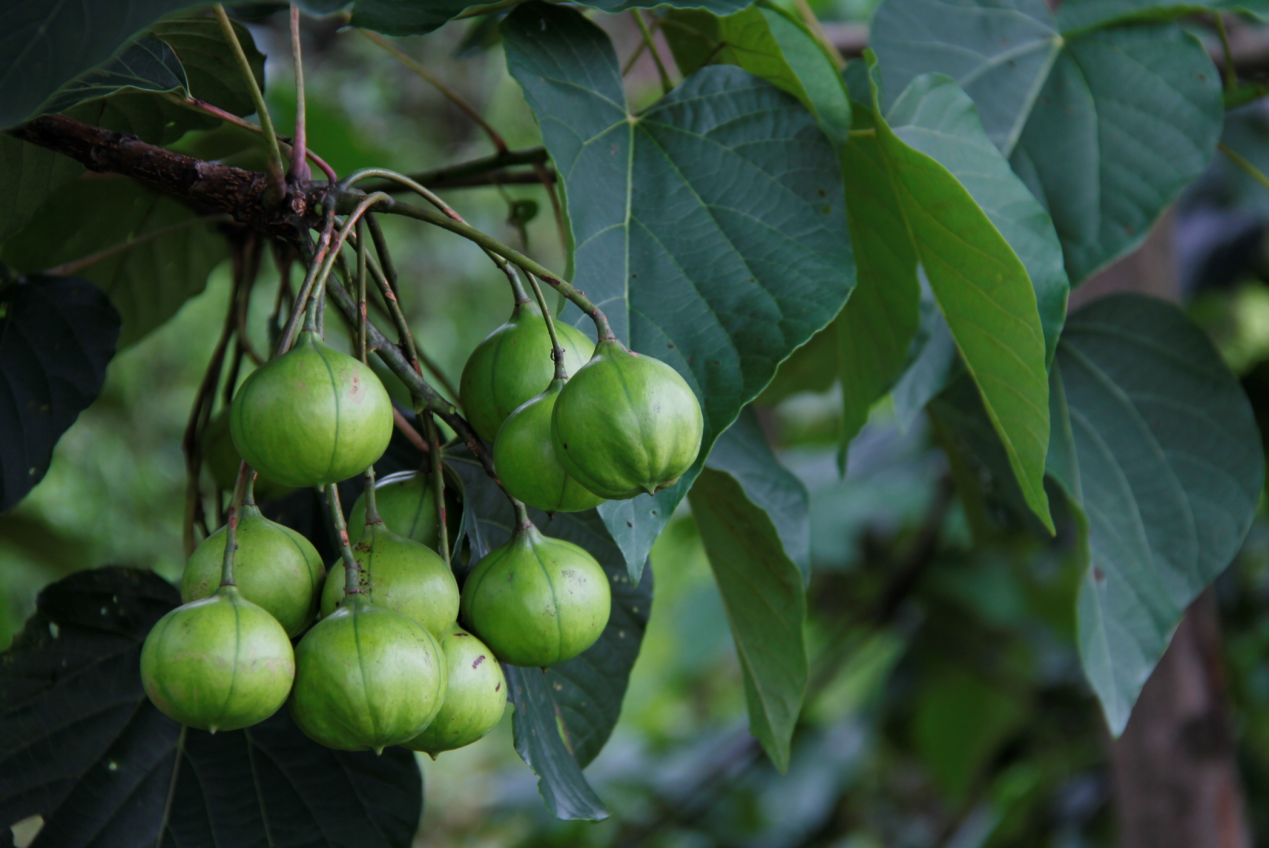

**Figure S2. Fruit setting of elite tung tree cv. Putaotong.**

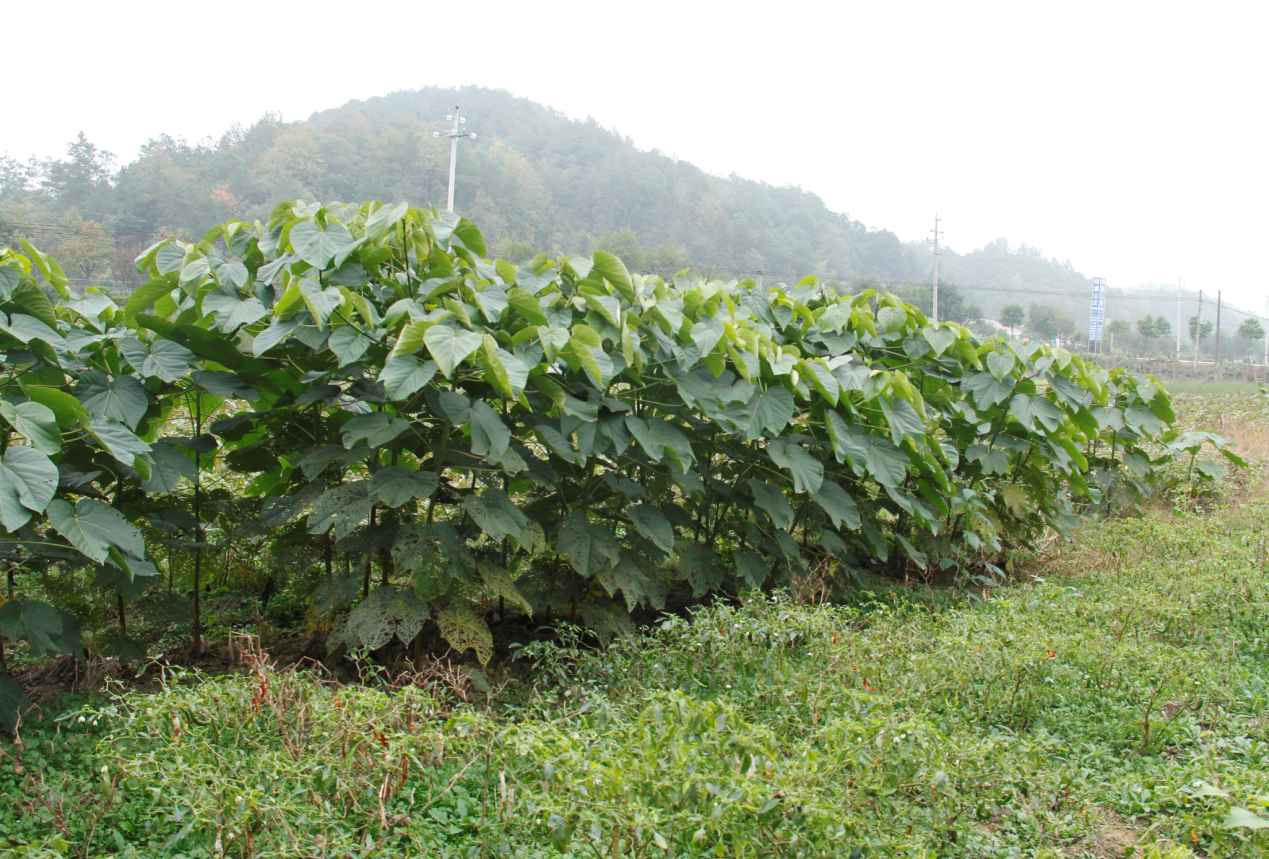

**Figure S3. The self-bred progenies of ‘Putaotong’.**

**The photo was taken in 2013.**

**Table S1 SSR markers used for heterozygosity estimation of tung tree self-bred progenies**

| **Marker** | **Primer sequences (5′– 3′)** | **Repeat** | **Size (bp)** | ***T_a_* (°C)** |
| --- | --- | --- | --- | --- |
| VFEST3 | F:TGGGAAACAATAATGGGAGG | (TG)_13_ | 392 | 59 |
|  | R:CGGGAACTAATAAAATCAAGCC |  |  |  |
| VFEST20 | F:TGGCATTGGCACTCACTACAG | (AT)_14_ | 256 | 59 |
|  | R:TAAGTTCACAAAAGCGGTCACA |  |  |  |
| VFEST51 | F:AGCGGCAACACCAGCAACT | (GCA)_7_ | 280 | 64 |
|  | R:TGGGTAGAGGGAGGAGGCAT |  |  |  |
| VFEST55 | F:GTCTCTCTCTTTCTATCTGTAACC | (CAC)_8_ | 248 | 59 |
|  | R:GCTTCAGGCTCTAAATCTTC |  |  |  |
| VFEST58 | F:ATCCCTATTGATGAGACC | (TAA)_10_ | 177 | 57 |
|  | R:TTAACACTAACTATACTTGACACT |  |  |  |
| VFEST60 | F:CTCCACCCAGTCTTCTACTTCAC | (GTGA)_5_ | 284 | 59 |
|  | R:ATCCAATAGCGTAAGATGACAAAG |  |  |  |
| VFEST68 | F:ATCAGGGCTTGGTTTTGGGT | (TC)_15_(TA)_10_ | 176 | 60 |
|  | R:ATAGGTAGGGGAGGCAGAGGAG |  |  |  |
| TT05 | F:AACTTGTCTGTGCATGTGCC | (CA) _7_ | 102 | 55 |
|  | R:TGCGTAACGTTCGAGTTGTC |  |  |  |
| TT06 | F:TTCTATGGCTTGAAGGGGTG | (CA) _17_ | 190 | 55 |
|  | R:TGCACTTGAATGTTTGTGCC |  |  |  |
| TT07 | F:CGCCTCGGTAATGCTCTAAC | (AC) _9_ | 163 | 55 |
|  | R:TCCCATTTCCTGAAAGCAAG |  |  |  |
| TT11 | F:CCAGTGGAAATGCAACAGAA | (AC) _9_ | 132 | 55 |
|  | R:AAAACAAAACCCAATGCAGC |  |  |  |
| TT14 | F:CGTATTGTCATCGTCTCCCA | (TG) _13_ | 104 | 55 |
|  | R:TCAGTCCTTCTCCTTATTTGAACA |  |  |  |
| TT16 | F:TGCTTGCCCAGTTTAGGTCT | (CA) _10_ | 127 | 55 |
|  | R:GCACACTTACCAAAACACACAA |  |  |  |
| TT17 | F:ATGAAGGCACTGTGAAGGGA | (GT) _6_ | 270 | 55 |
|  | R:CCATCCCAAATCCTCTAGCA |  |  |  |
| TT28 | F:GCTCTAGTTGGCCCTTCAAA | (TG) _13_ | 185 | 55 |
|  | R:AAGGGATGTTGCAGCTATGG |  |  |  |
| TT29 | F:ATGAGAGCATTGCACCACAC | (TC) _10_ | 117 | 55 |
|  | R:ATTTGCCATTCAAGTCTCGC |  |  |  |
| TT31 | F:TGGCAGCAAAGAAACTGAGA | (AG) _16_ | 169 | 55 |
|  | R:GAGTCCTGAGTAAGCCGTCG |  |  |  |
| TT32 | F:AATCAGTGGCAAGAAGTGGG | (AG) _22_ | 269 | 55 |
|  | R:GTCCTGAGTAAGGGCATCCA |  |  |  |
| TT34 | F:TTCCACACGAATTTTCTCCTG | (AT) _8_ (GT) _11_ | 137 | 55 |
|  | R:TGCAGATATTCTGCTGCCAC |  |  |  |
| TT39 | F:TCAATGCACTCAAGTCTGGC | (GA) _15_ | 107 | 55 |
|  | R:TCAGGTTCTTGTTTTTGCCC |  |  |  |
| TT42 | F:GTTCGATATCCAAGCCCTGT | (TG) _17_ | 158 | 55 |
|  | R:CCGCCCCTGCATACATATAG |  |  |  |
| TT50 | F:CGGGTCAAACCCACAAGATA | (AG) _14_ | 180 | 55 |
|  | R:AACTGACATTGTAAGGCAGCTC |  |  |  |
| TT53 | F:TGGTGACAGCTTTGCGTAAC | (TG) _11_ (GA) _11_ | 149 | 55 |
|  | R:TGATTAGCATTGCAGCAAAAA |  |  |  |
| TT55 | F:TCCTGAGTAAACAATCAACATCTCA | (AT) _6_ (AG) _8_ | 119 | 55 |
|  | R:TGGGACTAGCCTTGCCTCTA |  |  |  |
| TT57 | F:CAATAACAATGCGACAATGC | (AG) _9_ | 159 | 55 |
|  | R:AGACCACCCTGTTTTTGCTG |  |  |  |
| TT69 | F:GCACTATCCCCTTACGCAAC | (GA) _21_ | 143 | 55 |
|  | R:GTGCTTGTTCTGCTCCCTTC |  |  |  |
| TT71 | F:CACTCCTAGGTGAAATGCCC | (CT) 18 | 238 | 55 |
|  | R:TGCTCCAAAAATAGGAGTGGA |  |  |  |
