## Supplementary material for "The Tung Tree (*Vernicia Fordii*) Genome Provides A Resource for Understanding Genome Evolution and Oil Improvement": File S2

**File S2: Estimation of genome size and heterozygosity**

A total of 36.51 Gb data from a library of short insert sizes (500 bp) was used to estimate the genome size by a modified Lander-Waterman algorithm i.e. a formula G = Bnum/Bdepth = Knum/Kdepth [1]. In this formula, G, Bnum, Bdepth, Knum, and Kdepth refer to genome size, the total read number, the depth of bases, the total number of K-mer, and the overall depth of K-mer, respectively. Knum is calculated by N × (L – K + 1) where N, L, and K represent the number of K-mer, the length of reads, and the size of K-mer, respectively. Heterozygosity was estimated from the k-mer distribution. A single peak indicated that the sample genome harbored low heterozygosity level. The heterozygous rate was estimated with GenomeScope [2].
