## Supplementary material for "The Tung Tree (*Vernicia Fordii*) Genome Provides A Resource for Understanding Genome Evolution and Oil Improvement": File S3

**File S3: Whole-genome shotgun sequencing**

Nuclear DNA was isolated from fresh leaf tissues by using a DNeasy Plant Mini Kit (Qiagen, CA, USA). A series of DNA libraries were constructed with various insert sizes including 180 bp, 500 bp, 800 bp, 3 kb, 10 kb, and 15 kb, with paired-end libraries being generated for small insert sizes (< 1 kb) and mate-paired libraries for large insert sizes (> 1 kb). The DNA libraries were sequenced with an Illumina HiSeq 2000 sequencing platform (Illumina, CA, USA). In addition, SMRTbell template libraries were constructed using Pacific Biosciences’ template Prep Kit 1.0 (part 100-259-100) according to the protocol for 20 kb libraries. SMRTbell templates were bound to polymerases using the DNA/Polymerase Binding Kit P5 (part 100-259-100). Polymerase-template complexes were bound to magbeads by using Pacific Biosciences Magbead Bing kit (part 100-133-600), and sequencing was carried out on the PacBio RSII by using C3 sequencing reagents. Movie lengths with 240 min were recorded for each SMART cell. Subreads filtering was performed with Pacific Biosciences’ SMART analysis software.
