## Supplementary material for "The Tung Tree (*Vernicia Fordii*) Genome Provides A Resource for Understanding Genome Evolution and Oil Improvement": File S4

**File S4: Data quality control and Genome assembly and assessment**

The illumina raw data were filtered by FastQC [1] and NGS_QC_Toolkit [2] with the following steps: (1) Remove reads contaminated by adapters; (2) Trimming of continuous low-quality bases on both 5’ and 3’; (3) Remove reads that had Ns > 10% of the read length; (4) Remove duplicated reads caused by PCR amplification; (5) Trimming of paired-end reads if any reads with more than 50% low-quality bases. After removing low-quality reads, all the remaining data had high quality and was used for genome assembly. The whole genome assembly of tung tree was performed with a hierarchical assembly strategy due to its homozygous genome with many repetitive sequences. Firstly, Allpaths-LG was applied to assemble the clean Hiseq data, and GapCloser [3] was used to fill gaps and improve the quality of the scaffolds. Second, the paired-end or mate-pair reads were used to link the scaffolds/contigs into super-scaffold sequences using SSPACE [4]. Finally, PacBio reads were used to close gaps in scaffolds by PBJelly, an automated pipeline for gap filling and genome improvement that aligns long sequence reads to draft assembles in order to close or improve captured gaps [5]. The core eukaryotic genes (CEGs) were mapped against the genome assembly and the completeness of the CEGs were determined by Core Eukaryotic Genes Mapping Approach (CEGMA) with default parameters [6]. BUSCO analysis was also applied to validate the genome completeness [7]. In order to check the completeness of assembly, the RNA-seq reads from different tissues were mapped to the assembled genome using TopHat with default parameters [8]. Furthermore, unigenes of different transcriptomes were mapped to the assembled genome with BLAT using default parameters.

**References**

[1] S A. A quality control tool for high throughput seqence data. Referene Source 2010.
