## Supplementary material for "The Tung Tree (*Vernicia Fordii*) Genome Provides A Resource for Understanding Genome Evolution and Oil Improvement": File S5

**File S5: Gene prediction and functional annotation**

Gene prediction in the tung tree genome was conducted using *de novo* prediction, homology information and RNA-seq data. Augustus, GlimmHmm, and Geneid were used on the repeat masked genome for *de novo* prediction. For homology-based prediction, protein sequences from five sequenced plants (*A. thaliana, J. curcas, O. sativa, R. communis,* and *V. vinifera*) and Uniprot were initially mapped onto the *V. fordii* genome using tBlastn. The homologous genome sequences were aligned against the matching protein using GeneWise [1] for precise spliced alignments. Furthermore, the RNA-seq reads were aligned to the reference genome by TopHat to identify candidate exon regions and the donor/acceptor sites. Then the alignments were assembled into transcripts and open reading frames (ORFs) were confirmed to obtain reliable transcripts. Unigenes were aligned to reference genome for annotating protein-coding genes and alternatively spliced isoforms. All the predictions were combined by EVidenceModeler (EVM) [2] to generate a consensus gene set that was then aligned to transposon database, and the genes homologous to transposons were removed from the final gene set.

Gene functions were assigned according to the best match derived from the alignments to proteins annotated in SwissProt and TrEMBL [3] databases using Blastp, and the pathway in which the gene might be involved was annotated by KAAS [4]. Motifs and domains were annotated using Inter ProScan (Version 5.2-45.0) [5] by searching against publicly available databases in InterPro [6], including ProDom, PRINTS, Pfam, SMART, PANTHER, PROSITE etc. The Gene Ontology IDs for each gene were assigned using the corresponding InterPro entry.
