## Supplementary material for "The Tung Tree (*Vernicia Fordii*) Genome Provides A Resource for Understanding Genome Evolution and Oil Improvement": File S6

**File S6: Evolutionary analysis**

Besides *V. fordii*, the genome data of seven species including *A. thaliana, P. trichocarpa, V. vinifera, R. communis, M. esculenta, H. brasiliensis*, and *J. curcas* were used for evolutionary analysis including gene identification, positively selected genes (PSGs), recognition and whole genome duplication (WGD).

Single-copy gene families from the eight species were obtained and their protein sequences were aligned by Mafft [1] and back translated into CDS alignments. Four-fold degenerate sites of all the single-copy genes in each species were extracted and concatenated to be one supergene for phylogeny construction by the maximum likelihood (ML) method.

The divergence times for the eight species were estimated based on all single-copy genes and 4-fold degenerate sites with the program MCMCTree of the PAML package [2]. The neutral evolutionary rate was calculated via Bayes estimation with Markov Chain Monte Carlo algorithm.

Gene families which underwent expansions or contractions were identified using the CAFE (Computational Analysis of gene Family Evolution) program [3]. Orthologous genes were identified among eight species. High value of Ka/Ks usually indicates evolution under positive selection. The values of Ks and Ka substitution rates and the Ka/Ks ratio were estimated in each homologous cluster by using the Codeml program in the PAML package. The selection pressure of tung tree in the phylogenetic tree was calculated by Codeml. The significance of the identified positively selected genes (PSGs) was verified using a Chi-square test.

Whole-genome duplication events (WGD) were identified by two methods. First, the protein sequences of seven species (*P. trichocarpa, V. vinifera, V. fordii, J. curcas, R. communis, M. esculenta*, and *H. brasiliensis*) were searched against each other using Blastp with an e-value cutoff of less than 1e-5 and homologous genes were obtained using OrthoMCL. The 4DTv (four-fold synonymous third-codon transversion) rate of homologous genes was calculated, and homologous genes whose 4DTv sites numbered less than 20 were removed [4]. The distribution of the 4DTv values from seven species was plotted. Second, we also calculated the synonymous Ks to estimate duplication events using KaKs_calculator with an in-house perl script [5].

To detect the colinear regions, the protein sequences of tung tree, cassava and grapevine genomes were compared to each other using BLASTP with an e-value cutoff of 1e-10 e-value. The MCScanX [6] software was used to detect the syntenic blocks by chaining the BLASTP hits with a requirement at least five gene pairs per synteny block. The circular representation of these species’ genomes was plotted using the Circos package [7]. The comparison between tung tree and cassava or grapevine genomes was also performed using the MCScanX [6] based on the all-against-all BLASTP search of their protein sequences, and then visualized by the TBtools [8]. To more clearly present the synteny relationships, each chromosome of tung tree was selected to compare the genome of cassava and grapevine, respectively, and finally a total of 11 schematics were generated.
