## Supplementary material for "The Tung Tree (*Vernicia Fordii*) Genome Provides A Resource for Understanding Genome Evolution and Oil Improvement": File S7

**File S7: Repeat sequence analysis**

A *de novo* and homology-based approach was used to identify repetitive sequence and transposable elements (TEs) in the tung tree genome. A *de novo* repeat library by Repeat Modeler was constructed to generate the consensus sequences and classification information for each repeat family [1]. Repeat Masker was then applied for DNA-level identification using the *de novo* library and two databases, Repbase [2] and Mips-Redat [3]. At the protein level, Repeat Protein Mask was used to perform WU-BLATX against the TE protein database. The overlapping TEs belonging to the same type of repeats were collated and combined according to their coordinates in the genome. Tandem repeats were annotated with TRF (Tandem Repeat Finder) [4].

**Note 7.1: Identification of simple sequence repeats (SSRs)**

We searched the simple sequence repeats (SSRs) from the tung tree genome sequences by using MIcroSAtellite Identification Tool (MISA). We identified 663,931 SSRs with the density of 593.49 SSRs per Mb across the whole tung tree genome, of which compound format SSRs were 172,997, accounting for 26.06% (table 38). The minimum repeat unit size for mononucleotide was set at ten, for dinucleotide at six, and for tri-, tetra-, penta-, and hexa-nucleotide at five. In total, we identified six types of SSRs including mono-, di-, tri-, tetra-, penta-, and hexa-nucleotide. The annotated SSRs were mostly mononucleotide (263,069; 39.62%), and dinucleotide (88,805; 13.38%), and less trinucleotide (36,459; 5.49%), tetranucleotide (4,884; 0.74%), and pentanucleotide (2,073; 0.31%), and hexanucleotide (639; 0.10%) (table 39). These SSRs will provide valuable genetic markers to assist tung tree breeding programs.

**Note 7.2: Intact LTR-RT identification and insertion time estimation**

The long-terminal repeat retrotransposons (LTR-RTs) in the whole tung tree genome were identified with LTR-finder software. To classify the types of LTR-RTs, the repeat regions were masked using RepeatMasker and the LTR-RTs were located to RepeatMasker results. LTR insertion time was estimated by computing the nucleotide substitution rate of the two LTRs in an intact LTR-RT since they were assumed to be identical at the retroelement insertion time [5]. LTRs in each pair were aligned to calculate the substitution rate by the baseml algorithm of the PAML package with the TN93 model. The insertion time was counted by the formula of T=*K*/2r. T: element insertion time; r: synonymous mutation/site/year; *K*: the divergence between the LTRs and consensus sequence in the TE library. The substitution rate was 1.3×10^-8^ substitutions per site per year, 2-fold higher than the synonymous substitution rate of the coding region (6.5×10^-9^).

All intact LTR retrotransposons were classified into families by using BLASTClust and all-to-all BLAST of 5’ LTR sequences, followed by manual inspection [6]. The family classification standard was considered acceptable if more than 50% of the 5’ LTR mapped and sequence identify exceeded 80%. As a result, we classified 2,991 intact retrotransposon elements into a total of 1,130 families which included 89 multi-member families (>=5 intact members), 154 median-copy families (2-4 intact members) and 887 single-copy families (Table S42).

All intact LTR retrotransposons were classified into *Ty1/copia*, *Ty3/gypsy* and unclassified groups according to both RT gene similarity and the order of ORFs using PFAM [7]. The RT sequences were retrieved from each retrotransposon element and further checked by homology searches using ClustalW [8] against the published RTs that were downloaded from the Gypsy Database (GyDB) [9].

**Note 7.3: Phylogenetic analysis of LTR retrotransposon families**

We extracted nucleotide sequences of RTs from intact LTR retrotransposon elements and conducted alignments of amino acid sequences of RTs using ClustalW2 [8]. Consequently, unrooted neighbor-joining (NJ) phylogenetic trees were generated by using MEGA 6 [10]. In total, 347 were grouped into Ty1/copia families, including Ale, Angela, Bianca, Horpia2, Lkya, Owis, Rare1, and Rare2, of which Ale was most abundant (115) accounting for 33.14% (Figure 3 A and B; Table S43). A total of 622 were Ty3/gypsy sequences, including Bagy2, Cereba, Dagan, Erika, GA, Geneva, Laura, and Retrosat, of which Dagan was most abundant (373) accounting for 59.97% (Figure 3 A and B; Table S43).

**Note 7.4: Expression of LTR retrotransposons**

To understand the activation of the most abundant LTR retrotransposons in the tung tree genome, expression levels of LTR retrotransposons were estimated by computing the number of reads in LTR retrotransposons within RNA-Seq datasets from six tissues including 10 samples (root, stem, young leaf, female flower, male flower and seed at five developing stages) generated in this study. Transcriptome data from five seed samples were normalized to be used for expression calculation. The transcriptomes were masked by RepeatMasker (version 4.0.5) using the same TE library as used for annotation of the tung tree genome. TopHat2 [11] (version 2.0.5) and StringTie (version 1.3.0) [12] were used to count the number of reads in RNA-Seq data (Table S44).

Based on our RNA-seq data, we found 1,738 out of the total 2,991 LTR retrotransposons exhibited expression across six tissues and different LTR retrotransposon families exhibited distinct expression patterns. Generally, *Ty3/gypsy* LTR retrotransposons exhibited higher expression levels than Ty1/Copia retrotransposons, ranging from 0.71-fold in seed to 4.09-fold in leaf with approximately two-fold higher on average (Table S44). Among the 1,738 LTR retrotransposons, 701 showed the highest expression level in seeds, of which 60.77% belongs to high-copy families (Figure 3D). However, VL0631, a single-copy LTR retrotransposon, was the most highly expressed in seeds (Supplementary table S45). In the top 29 LTR retrotransposons highly expressed in seeds (FMKM value >=1), high-copy families, median-copy families and sing-copy families are 3, 6, and 20, accounting for 10.34%, 20.69%, and 68.97%, respectively (Table S45). In addition, 184, 204, 244, 148, and 257 LTR retrotransposons exhibited the highest expression levels in root, stem, leaf, female flower and male flower, respectively (Figure S22). Among these genes, similar to seeds, high-copy LTR families also accounted for the highest proportion in the other five tissues and single-copy families accounted for the highest proportion in the most highly expressed LTR retrotransposons (Table S46-S50; Figure S22).

**Supplementary Figures** **and Tables**

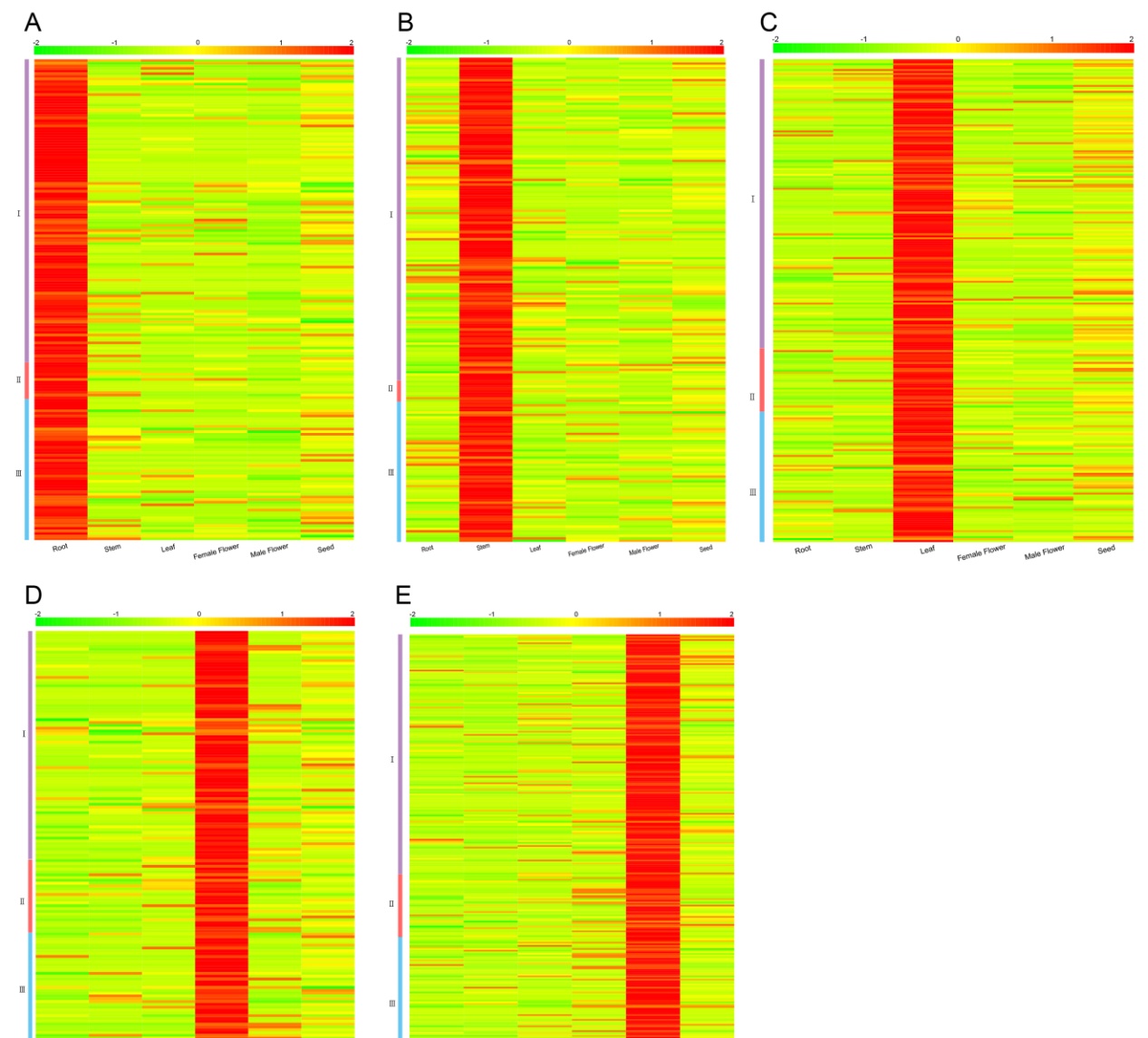

**Figure S22. Heat map of expression patterns of tung tree LTR retrotransposons.** A-E: indicates LTR retrotransposons showing the highest expression level in root, stem, leaf, female flower and male flower, respectively

**Table S38. Distribution of simple sequence repeats (SSRs) in tung tree genome**

|  | Number/Size |
| --- | --- |
| Total examined sequences | 4,577/ 1,118,693,778bp |
| Total identified SSR | 663,931 |
| Compound format | 172,997 |
| Sequences contain SSR | 3,800 |
| Sequences containing SSR(>1) | 3,426 |

**Table S39. Summary of SSR types in tung tree genome**

| Type | Unit size (repeat number) | Number | Percentage |
| --- | --- | --- | --- |
| p1 | 1 (>=10) | 263,069 | 39.62% |
| p2 | 2 (>=6) | 88,805 | 13.38% |
| p3 | 3 (>=5) | 36,459 | 5.49% |
| p4 | 4 (>=5) | 4,884 | 0.74% |
| p5 | 5 (>=5) | 2,073 | 0.31% |
| p6 | 6 (>=5) | 639 | 0.10% |

p1-p6 indicate six types of SSRs, i.e., mononucleotide, dinucleotide, trinucleotide tetranucleotide, pentanucleotide and hexanucleotide, respectively.

**Table S42 Classification of LTR families**

| **LTR families** | **Copy numuber** | **Proportion (%)** |
| --- | --- | --- |
| VL0001 | 130 | 4.346372451 |
| VL0002 | 117 | 3.911735206 |
| VL0003 | 114 | 3.811434303 |
| VL0004 | 75 | 2.507522568 |
| VL0005 | 70 | 2.340354397 |
| VL0006 | 70 | 2.340354397 |
| VL0007 | 64 | 2.139752591 |
| VL0008 | 61 | 2.039451688 |
| VL0009 | 61 | 2.039451688 |
| VL0010 | 46 | 1.537947175 |
| VL0011 | 42 | 1.404212638 |
| VL0012 | 42 | 1.404212638 |
| VL0013 | 37 | 1.237044467 |
| VL0014 | 37 | 1.237044467 |
| VL0015 | 37 | 1.237044467 |
| VL0016 | 32 | 1.069876296 |
| VL0017 | 32 | 1.069876296 |
| VL0018 | 30 | 1.003009027 |
| VL0019 | 23 | 0.768973587 |
| VL0020 | 21 | 0.702106319 |
| VL0021 | 20 | 0.668672685 |
| VL0022 | 20 | 0.668672685 |
| VL0023 | 18 | 0.601805416 |
| VL0024 | 18 | 0.601805416 |
| VL0025 | 17 | 0.568371782 |
| VL0026 | 17 | 0.568371782 |
| VL0027 | 16 | 0.534938148 |
| VL0028 | 15 | 0.501504514 |
| VL0029 | 14 | 0.468070879 |
| VL0030 | 13 | 0.434637245 |
| VL0031 | 13 | 0.434637245 |
| VL0032 | 13 | 0.434637245 |
| VL0033 | 13 | 0.434637245 |
| VL0034 | 12 | 0.401203611 |
| VL0035 | 12 | 0.401203611 |
| VL0036 | 12 | 0.401203611 |
| VL0037 | 12 | 0.401203611 |
| VL0038 | 10 | 0.334336342 |
| VL0039 | 9 | 0.300902708 |
| VL0040 | 9 | 0.300902708 |
| VL0041 | 9 | 0.300902708 |
| VL0042 | 9 | 0.300902708 |
| VL0043 | 8 | 0.267469074 |
| VL0044 | 8 | 0.267469074 |
| VL0045 | 8 | 0.267469074 |
| VL0046 | 8 | 0.267469074 |
| VL0047 | 8 | 0.267469074 |
| VL0048 | 7 | 0.23403544 |
| VL0049 | 7 | 0.23403544 |
| VL0050 | 7 | 0.23403544 |
| VL0051 | 7 | 0.23403544 |
| VL0052 | 7 | 0.23403544 |
| VL0053 | 7 | 0.23403544 |
| VL0054 | 6 | 0.200601805 |
| VL0055 | 6 | 0.200601805 |
| VL0056 | 6 | 0.200601805 |
| VL0057 | 6 | 0.200601805 |
| VL0058 | 6 | 0.200601805 |
| VL0059 | 6 | 0.200601805 |
| VL0060 | 6 | 0.200601805 |
| VL0061 | 6 | 0.200601805 |
| VL0062 | 6 | 0.200601805 |
| VL0063 | 6 | 0.200601805 |
| VL0064 | 6 | 0.200601805 |
| VL0065 | 6 | 0.200601805 |
| VL0066 | 6 | 0.200601805 |
| VL0067 | 6 | 0.200601805 |
| VL0068 | 6 | 0.200601805 |
| VL0069 | 6 | 0.200601805 |
| VL0070 | 6 | 0.200601805 |
| VL0071 | 6 | 0.200601805 |
| VL0072 | 6 | 0.200601805 |
| VL0073 | 5 | 0.167168171 |
| VL0074 | 5 | 0.167168171 |
| VL0075 | 5 | 0.167168171 |
| VL0076 | 5 | 0.167168171 |
| VL0077 | 5 | 0.167168171 |
| VL0078 | 5 | 0.167168171 |
| VL0079 | 5 | 0.167168171 |
| VL0080 | 5 | 0.167168171 |
| VL0081 | 5 | 0.167168171 |
| VL0082 | 5 | 0.167168171 |
| VL0083 | 5 | 0.167168171 |
| VL0084 | 5 | 0.167168171 |
| VL0085 | 5 | 0.167168171 |
| VL0086 | 5 | 0.167168171 |
| VL0087 | 5 | 0.167168171 |
| VL0088 | 5 | 0.167168171 |
| VL0089 | 5 | 0.167168171 |
| VL0090 | 4 | 0.133734537 |
| VL0091 | 4 | 0.133734537 |
| VL0092 | 4 | 0.133734537 |
| VL0093 | 4 | 0.133734537 |
| VL0094 | 4 | 0.133734537 |
| VL0095 | 4 | 0.133734537 |
| VL0096 | 4 | 0.133734537 |
| VL0097 | 4 | 0.133734537 |
| VL0098 | 4 | 0.133734537 |
| VL0099 | 4 | 0.133734537 |
| VL0100 | 4 | 0.133734537 |
| VL0101 | 4 | 0.133734537 |
| VL0102 | 4 | 0.133734537 |
| VL0103 | 4 | 0.133734537 |
| VL0104 | 4 | 0.133734537 |
| VL0105 | 4 | 0.133734537 |
| VL0106 | 4 | 0.133734537 |
| VL0107 | 4 | 0.133734537 |
| VL0108 | 3 | 0.100300903 |
| VL0109 | 3 | 0.100300903 |
| VL0110 | 3 | 0.100300903 |
| VL0111 | 3 | 0.100300903 |
| VL0112 | 3 | 0.100300903 |
| VL0113 | 3 | 0.100300903 |
| VL0114 | 3 | 0.100300903 |
| VL0115 | 3 | 0.100300903 |
| VL0116 | 3 | 0.100300903 |
| VL0117 | 3 | 0.100300903 |
| VL0118 | 3 | 0.100300903 |
| VL0119 | 3 | 0.100300903 |
| VL0120 | 3 | 0.100300903 |
| VL0121 | 3 | 0.100300903 |
| VL0122 | 3 | 0.100300903 |
| VL0123 | 3 | 0.100300903 |
| VL0124 | 3 | 0.100300903 |
| VL0125 | 3 | 0.100300903 |
| VL0126 | 3 | 0.100300903 |
| VL0127 | 3 | 0.100300903 |
| VL0128 | 3 | 0.100300903 |
| VL0129 | 3 | 0.100300903 |
| VL0130 | 3 | 0.100300903 |
| VL0131 | 3 | 0.100300903 |
| VL0132 | 3 | 0.100300903 |
| VL0133 | 3 | 0.100300903 |
| VL0134 | 3 | 0.100300903 |
| VL0135 | 3 | 0.100300903 |
| VL0136 | 3 | 0.100300903 |
| VL0137 | 3 | 0.100300903 |
| VL0138 | 3 | 0.100300903 |
| VL0139 | 3 | 0.100300903 |
| VL0140 | 3 | 0.100300903 |
| VL0141 | 3 | 0.100300903 |
| VL0142 | 3 | 0.100300903 |
| VL0143 | 3 | 0.100300903 |
| VL0144 | 3 | 0.100300903 |
| VL0145 | 2 | 0.066867268 |
| VL0146 | 2 | 0.066867268 |
| VL0147 | 2 | 0.066867268 |
| VL0148 | 2 | 0.066867268 |
| VL0149 | 2 | 0.066867268 |
| VL0150 | 2 | 0.066867268 |
| VL0151 | 2 | 0.066867268 |
| VL0152 | 2 | 0.066867268 |
| VL0153 | 2 | 0.066867268 |
| VL0154 | 2 | 0.066867268 |
| VL0155 | 2 | 0.066867268 |
| VL0156 | 2 | 0.066867268 |
| VL0157 | 2 | 0.066867268 |
| VL0158 | 2 | 0.066867268 |
| VL0159 | 2 | 0.066867268 |
| VL0160 | 2 | 0.066867268 |
| VL0161 | 2 | 0.066867268 |
| VL0162 | 2 | 0.066867268 |
| VL0163 | 2 | 0.066867268 |
| VL0164 | 2 | 0.066867268 |
| VL0165 | 2 | 0.066867268 |
| VL0166 | 2 | 0.066867268 |
| VL0167 | 2 | 0.066867268 |
| VL0168 | 2 | 0.066867268 |
| VL0169 | 2 | 0.066867268 |
| VL0170 | 2 | 0.066867268 |
| VL0171 | 2 | 0.066867268 |
| VL0172 | 2 | 0.066867268 |
| VL0173 | 2 | 0.066867268 |
| VL0174 | 2 | 0.066867268 |
| VL0175 | 2 | 0.066867268 |
| VL0176 | 2 | 0.066867268 |
| VL0177 | 2 | 0.066867268 |
| VL0178 | 2 | 0.066867268 |
| VL0179 | 2 | 0.066867268 |
| VL0180 | 2 | 0.066867268 |
| VL0181 | 2 | 0.066867268 |
| VL0182 | 2 | 0.066867268 |
| VL0183 | 2 | 0.066867268 |
| VL0184 | 2 | 0.066867268 |
| VL0185 | 2 | 0.066867268 |
| VL0186 | 2 | 0.066867268 |
| VL0187 | 2 | 0.066867268 |
| VL0188 | 2 | 0.066867268 |
| VL0189 | 2 | 0.066867268 |
| VL0190 | 2 | 0.066867268 |
| VL0191 | 2 | 0.066867268 |
| VL0192 | 2 | 0.066867268 |
| VL0193 | 2 | 0.066867268 |
| VL0194 | 2 | 0.066867268 |
| VL0195 | 2 | 0.066867268 |
| VL0196 | 2 | 0.066867268 |
| VL0197 | 2 | 0.066867268 |
| VL0198 | 2 | 0.066867268 |
| VL0199 | 2 | 0.066867268 |
| VL0200 | 2 | 0.066867268 |
| VL0201 | 2 | 0.066867268 |
| VL0202 | 2 | 0.066867268 |
| VL0203 | 2 | 0.066867268 |
| VL0204 | 2 | 0.066867268 |
| VL0205 | 2 | 0.066867268 |
| VL0206 | 2 | 0.066867268 |
| VL0207 | 2 | 0.066867268 |
| VL0208 | 2 | 0.066867268 |
| VL0209 | 2 | 0.066867268 |
| VL0210 | 2 | 0.066867268 |
| VL0211 | 2 | 0.066867268 |
| VL0212 | 2 | 0.066867268 |
| VL0213 | 2 | 0.066867268 |
| VL0214 | 2 | 0.066867268 |
| VL0215 | 2 | 0.066867268 |
| VL0216 | 2 | 0.066867268 |
| VL0217 | 2 | 0.066867268 |
| VL0218 | 2 | 0.066867268 |
| VL0219 | 2 | 0.066867268 |
| VL0220 | 2 | 0.066867268 |
| VL0221 | 2 | 0.066867268 |
| VL0222 | 2 | 0.066867268 |
| VL0223 | 2 | 0.066867268 |
| VL0224 | 2 | 0.066867268 |
| VL0225 | 2 | 0.066867268 |
| VL0226 | 2 | 0.066867268 |
| VL0227 | 2 | 0.066867268 |
| VL0228 | 2 | 0.066867268 |
| VL0229 | 2 | 0.066867268 |
| VL0230 | 2 | 0.066867268 |
| VL0231 | 2 | 0.066867268 |
| VL0232 | 2 | 0.066867268 |
| VL0233 | 2 | 0.066867268 |
| VL0234 | 2 | 0.066867268 |
| VL0235 | 2 | 0.066867268 |
| VL0236 | 2 | 0.066867268 |
| VL0237 | 2 | 0.066867268 |
| VL0238 | 2 | 0.066867268 |
| VL0239 | 2 | 0.066867268 |
| VL0240 | 2 | 0.066867268 |
| VL0241 | 2 | 0.066867268 |
| VL0242 | 2 | 0.066867268 |
| VL0243 | 2 | 0.066867268 |
| VL0244 | 1 | 0.033433634 |
| VL0245 | 1 | 0.033433634 |
| VL0246 | 1 | 0.033433634 |
| VL0247 | 1 | 0.033433634 |
| VL0248 | 1 | 0.033433634 |
| VL0249 | 1 | 0.033433634 |
| VL0250 | 1 | 0.033433634 |
| VL0251 | 1 | 0.033433634 |
| VL0252 | 1 | 0.033433634 |
| VL0253 | 1 | 0.033433634 |
| VL0254 | 1 | 0.033433634 |
| VL0255 | 1 | 0.033433634 |
| VL0256 | 1 | 0.033433634 |
| VL0257 | 1 | 0.033433634 |
| VL0258 | 1 | 0.033433634 |
| VL0259 | 1 | 0.033433634 |
| VL0260 | 1 | 0.033433634 |
| VL0261 | 1 | 0.033433634 |
| VL0262 | 1 | 0.033433634 |
| VL0263 | 1 | 0.033433634 |
| VL0264 | 1 | 0.033433634 |
| VL0265 | 1 | 0.033433634 |
| VL0266 | 1 | 0.033433634 |
| VL0267 | 1 | 0.033433634 |
| VL0268 | 1 | 0.033433634 |
| VL0269 | 1 | 0.033433634 |
| VL0270 | 1 | 0.033433634 |
| VL0271 | 1 | 0.033433634 |
| VL0272 | 1 | 0.033433634 |
| VL0273 | 1 | 0.033433634 |
| VL0274 | 1 | 0.033433634 |
| VL0275 | 1 | 0.033433634 |
| VL0276 | 1 | 0.033433634 |
| VL0277 | 1 | 0.033433634 |
| VL0278 | 1 | 0.033433634 |
| VL0279 | 1 | 0.033433634 |
| VL0280 | 1 | 0.033433634 |
| VL0281 | 1 | 0.033433634 |
| VL0282 | 1 | 0.033433634 |
| VL0283 | 1 | 0.033433634 |
| VL0284 | 1 | 0.033433634 |
| VL0285 | 1 | 0.033433634 |
| VL0286 | 1 | 0.033433634 |
| VL0287 | 1 | 0.033433634 |
| VL0288 | 1 | 0.033433634 |
| VL0289 | 1 | 0.033433634 |
| VL0290 | 1 | 0.033433634 |
| VL0291 | 1 | 0.033433634 |
| VL0292 | 1 | 0.033433634 |
| VL0293 | 1 | 0.033433634 |
| VL0294 | 1 | 0.033433634 |
| VL0295 | 1 | 0.033433634 |
| VL0296 | 1 | 0.033433634 |
| VL0297 | 1 | 0.033433634 |
| VL0298 | 1 | 0.033433634 |
| VL0299 | 1 | 0.033433634 |
| VL0300 | 1 | 0.033433634 |
| VL0301 | 1 | 0.033433634 |
| VL0302 | 1 | 0.033433634 |
| VL0303 | 1 | 0.033433634 |
| VL0304 | 1 | 0.033433634 |
| VL0305 | 1 | 0.033433634 |
| VL0306 | 1 | 0.033433634 |
| VL0307 | 1 | 0.033433634 |
| VL0308 | 1 | 0.033433634 |
| VL0309 | 1 | 0.033433634 |
| VL0310 | 1 | 0.033433634 |
| VL0311 | 1 | 0.033433634 |
| VL0312 | 1 | 0.033433634 |
| VL0313 | 1 | 0.033433634 |
| VL0314 | 1 | 0.033433634 |
| VL0315 | 1 | 0.033433634 |
| VL0316 | 1 | 0.033433634 |
| VL0317 | 1 | 0.033433634 |
| VL0318 | 1 | 0.033433634 |
| VL0319 | 1 | 0.033433634 |
| VL0320 | 1 | 0.033433634 |
| VL0321 | 1 | 0.033433634 |
| VL0322 | 1 | 0.033433634 |
| VL0323 | 1 | 0.033433634 |
| VL0324 | 1 | 0.033433634 |
| VL0325 | 1 | 0.033433634 |
| VL0326 | 1 | 0.033433634 |
| VL0327 | 1 | 0.033433634 |
| VL0328 | 1 | 0.033433634 |
| VL0329 | 1 | 0.033433634 |
| VL0330 | 1 | 0.033433634 |
| VL0331 | 1 | 0.033433634 |
| VL0332 | 1 | 0.033433634 |
| VL0333 | 1 | 0.033433634 |
| VL0334 | 1 | 0.033433634 |
| VL0335 | 1 | 0.033433634 |
| VL0336 | 1 | 0.033433634 |
| VL0337 | 1 | 0.033433634 |
| VL0338 | 1 | 0.033433634 |
| VL0339 | 1 | 0.033433634 |
| VL0340 | 1 | 0.033433634 |
| VL0341 | 1 | 0.033433634 |
| VL0342 | 1 | 0.033433634 |
| VL0343 | 1 | 0.033433634 |
| VL0344 | 1 | 0.033433634 |
| VL0345 | 1 | 0.033433634 |
| VL0346 | 1 | 0.033433634 |
| VL0347 | 1 | 0.033433634 |
| VL0348 | 1 | 0.033433634 |
| VL0349 | 1 | 0.033433634 |
| VL0350 | 1 | 0.033433634 |
| VL0351 | 1 | 0.033433634 |
| VL0352 | 1 | 0.033433634 |
| VL0353 | 1 | 0.033433634 |
| VL0354 | 1 | 0.033433634 |
| VL0355 | 1 | 0.033433634 |
| VL0356 | 1 | 0.033433634 |
| VL0357 | 1 | 0.033433634 |
| VL0358 | 1 | 0.033433634 |
| VL0359 | 1 | 0.033433634 |
| VL0360 | 1 | 0.033433634 |
| VL0361 | 1 | 0.033433634 |
| VL0362 | 1 | 0.033433634 |
| VL0363 | 1 | 0.033433634 |
| VL0364 | 1 | 0.033433634 |
| VL0365 | 1 | 0.033433634 |
| VL0366 | 1 | 0.033433634 |
| VL0367 | 1 | 0.033433634 |
| VL0368 | 1 | 0.033433634 |
| VL0369 | 1 | 0.033433634 |
| VL0370 | 1 | 0.033433634 |
| VL0371 | 1 | 0.033433634 |
| VL0372 | 1 | 0.033433634 |
| VL0373 | 1 | 0.033433634 |
| VL0374 | 1 | 0.033433634 |
| VL0375 | 1 | 0.033433634 |
| VL0376 | 1 | 0.033433634 |
| VL0377 | 1 | 0.033433634 |
| VL0378 | 1 | 0.033433634 |
| VL0379 | 1 | 0.033433634 |
| VL0380 | 1 | 0.033433634 |
| VL0381 | 1 | 0.033433634 |
| VL0382 | 1 | 0.033433634 |
| VL0383 | 1 | 0.033433634 |
| VL0384 | 1 | 0.033433634 |
| VL0385 | 1 | 0.033433634 |
| VL0386 | 1 | 0.033433634 |
| VL0387 | 1 | 0.033433634 |
| VL0388 | 1 | 0.033433634 |
| VL0389 | 1 | 0.033433634 |
| VL0390 | 1 | 0.033433634 |
| VL0391 | 1 | 0.033433634 |
| VL0392 | 1 | 0.033433634 |
| VL0393 | 1 | 0.033433634 |
| VL0394 | 1 | 0.033433634 |
| VL0395 | 1 | 0.033433634 |
| VL0396 | 1 | 0.033433634 |
| VL0397 | 1 | 0.033433634 |
| VL0398 | 1 | 0.033433634 |
| VL0399 | 1 | 0.033433634 |
| VL0400 | 1 | 0.033433634 |
| VL0401 | 1 | 0.033433634 |
| VL0402 | 1 | 0.033433634 |
| VL0403 | 1 | 0.033433634 |
| VL0404 | 1 | 0.033433634 |
| VL0405 | 1 | 0.033433634 |
| VL0406 | 1 | 0.033433634 |
| VL0407 | 1 | 0.033433634 |
| VL0408 | 1 | 0.033433634 |
| VL0409 | 1 | 0.033433634 |
| VL0410 | 1 | 0.033433634 |
| VL0411 | 1 | 0.033433634 |
| VL0412 | 1 | 0.033433634 |
| VL0413 | 1 | 0.033433634 |
| VL0414 | 1 | 0.033433634 |
| VL0415 | 1 | 0.033433634 |
| VL0416 | 1 | 0.033433634 |
| VL0417 | 1 | 0.033433634 |
| VL0418 | 1 | 0.033433634 |
| VL0419 | 1 | 0.033433634 |
| VL0420 | 1 | 0.033433634 |
| VL0421 | 1 | 0.033433634 |
| VL0422 | 1 | 0.033433634 |
| VL0423 | 1 | 0.033433634 |
| VL0424 | 1 | 0.033433634 |
| VL0425 | 1 | 0.033433634 |
| VL0426 | 1 | 0.033433634 |
| VL0427 | 1 | 0.033433634 |
| VL0428 | 1 | 0.033433634 |
| VL0429 | 1 | 0.033433634 |
| VL0430 | 1 | 0.033433634 |
| VL0431 | 1 | 0.033433634 |
| VL0432 | 1 | 0.033433634 |
| VL0433 | 1 | 0.033433634 |
| VL0434 | 1 | 0.033433634 |
| VL0435 | 1 | 0.033433634 |
| VL0436 | 1 | 0.033433634 |
| VL0437 | 1 | 0.033433634 |
| VL0438 | 1 | 0.033433634 |
| VL0439 | 1 | 0.033433634 |
| VL0440 | 1 | 0.033433634 |
| VL0441 | 1 | 0.033433634 |
| VL0442 | 1 | 0.033433634 |
| VL0443 | 1 | 0.033433634 |
| VL0444 | 1 | 0.033433634 |
| VL0445 | 1 | 0.033433634 |
| VL0446 | 1 | 0.033433634 |
| VL0447 | 1 | 0.033433634 |
| VL0448 | 1 | 0.033433634 |
| VL0449 | 1 | 0.033433634 |
| VL0450 | 1 | 0.033433634 |
| VL0451 | 1 | 0.033433634 |
| VL0452 | 1 | 0.033433634 |
| VL0453 | 1 | 0.033433634 |
| VL0454 | 1 | 0.033433634 |
| VL0455 | 1 | 0.033433634 |
| VL0456 | 1 | 0.033433634 |
| VL0457 | 1 | 0.033433634 |
| VL0458 | 1 | 0.033433634 |
| VL0459 | 1 | 0.033433634 |
| VL0460 | 1 | 0.033433634 |
| VL0461 | 1 | 0.033433634 |
| VL0462 | 1 | 0.033433634 |
| VL0463 | 1 | 0.033433634 |
| VL0464 | 1 | 0.033433634 |
| VL0465 | 1 | 0.033433634 |
| VL0466 | 1 | 0.033433634 |
| VL0467 | 1 | 0.033433634 |
| VL0468 | 1 | 0.033433634 |
| VL0469 | 1 | 0.033433634 |
| VL0470 | 1 | 0.033433634 |
| VL0471 | 1 | 0.033433634 |
| VL0472 | 1 | 0.033433634 |
| VL0473 | 1 | 0.033433634 |
| VL0474 | 1 | 0.033433634 |
| VL0475 | 1 | 0.033433634 |
| VL0476 | 1 | 0.033433634 |
| VL0477 | 1 | 0.033433634 |
| VL0478 | 1 | 0.033433634 |
| VL0479 | 1 | 0.033433634 |
| VL0480 | 1 | 0.033433634 |
| VL0481 | 1 | 0.033433634 |
| VL0482 | 1 | 0.033433634 |
| VL0483 | 1 | 0.033433634 |
| VL0484 | 1 | 0.033433634 |
| VL0485 | 1 | 0.033433634 |
| VL0486 | 1 | 0.033433634 |
| VL0487 | 1 | 0.033433634 |
| VL0488 | 1 | 0.033433634 |
| VL0489 | 1 | 0.033433634 |
| VL0490 | 1 | 0.033433634 |
| VL0491 | 1 | 0.033433634 |
| VL0492 | 1 | 0.033433634 |
| VL0493 | 1 | 0.033433634 |
| VL0494 | 1 | 0.033433634 |
| VL0495 | 1 | 0.033433634 |
| VL0496 | 1 | 0.033433634 |
| VL0497 | 1 | 0.033433634 |
| VL0498 | 1 | 0.033433634 |
| VL0499 | 1 | 0.033433634 |
| VL0500 | 1 | 0.033433634 |
| VL0501 | 1 | 0.033433634 |
| VL0502 | 1 | 0.033433634 |
| VL0503 | 1 | 0.033433634 |
| VL0504 | 1 | 0.033433634 |
| VL0505 | 1 | 0.033433634 |
| VL0506 | 1 | 0.033433634 |
| VL0507 | 1 | 0.033433634 |
| VL0508 | 1 | 0.033433634 |
| VL0509 | 1 | 0.033433634 |
| VL0510 | 1 | 0.033433634 |
| VL0511 | 1 | 0.033433634 |
| VL0512 | 1 | 0.033433634 |
| VL0513 | 1 | 0.033433634 |
| VL0514 | 1 | 0.033433634 |
| VL0515 | 1 | 0.033433634 |
| VL0516 | 1 | 0.033433634 |
| VL0517 | 1 | 0.033433634 |
| VL0518 | 1 | 0.033433634 |
| VL0519 | 1 | 0.033433634 |
| VL0520 | 1 | 0.033433634 |
| VL0521 | 1 | 0.033433634 |
| VL0522 | 1 | 0.033433634 |
| VL0523 | 1 | 0.033433634 |
| VL0524 | 1 | 0.033433634 |
| VL0525 | 1 | 0.033433634 |
| VL0526 | 1 | 0.033433634 |
| VL0527 | 1 | 0.033433634 |
| VL0528 | 1 | 0.033433634 |
| VL0529 | 1 | 0.033433634 |
| VL0530 | 1 | 0.033433634 |
| VL0531 | 1 | 0.033433634 |
| VL0532 | 1 | 0.033433634 |
| VL0533 | 1 | 0.033433634 |
| VL0534 | 1 | 0.033433634 |
| VL0535 | 1 | 0.033433634 |
| VL0536 | 1 | 0.033433634 |
| VL0537 | 1 | 0.033433634 |
| VL0538 | 1 | 0.033433634 |
| VL0539 | 1 | 0.033433634 |
| VL0540 | 1 | 0.033433634 |
| VL0541 | 1 | 0.033433634 |
| VL0542 | 1 | 0.033433634 |
| VL0543 | 1 | 0.033433634 |
| VL0544 | 1 | 0.033433634 |
| VL0545 | 1 | 0.033433634 |
| VL0546 | 1 | 0.033433634 |
| VL0547 | 1 | 0.033433634 |
| VL0548 | 1 | 0.033433634 |
| VL0549 | 1 | 0.033433634 |
| VL0550 | 1 | 0.033433634 |
| VL0551 | 1 | 0.033433634 |
| VL0552 | 1 | 0.033433634 |
| VL0553 | 1 | 0.033433634 |
| VL0554 | 1 | 0.033433634 |
| VL0555 | 1 | 0.033433634 |
| VL0556 | 1 | 0.033433634 |
| VL0557 | 1 | 0.033433634 |
| VL0558 | 1 | 0.033433634 |
| VL0559 | 1 | 0.033433634 |
| VL0560 | 1 | 0.033433634 |
| VL0561 | 1 | 0.033433634 |
| VL0562 | 1 | 0.033433634 |
| VL0563 | 1 | 0.033433634 |
| VL0564 | 1 | 0.033433634 |
| VL0565 | 1 | 0.033433634 |
| VL0566 | 1 | 0.033433634 |
| VL0567 | 1 | 0.033433634 |
| VL0568 | 1 | 0.033433634 |
| VL0569 | 1 | 0.033433634 |
| VL0570 | 1 | 0.033433634 |
| VL0571 | 1 | 0.033433634 |
| VL0572 | 1 | 0.033433634 |
| VL0573 | 1 | 0.033433634 |
| VL0574 | 1 | 0.033433634 |
| VL0575 | 1 | 0.033433634 |
| VL0576 | 1 | 0.033433634 |
| VL0577 | 1 | 0.033433634 |
| VL0578 | 1 | 0.033433634 |
| VL0579 | 1 | 0.033433634 |
| VL0580 | 1 | 0.033433634 |
| VL0581 | 1 | 0.033433634 |
| VL0582 | 1 | 0.033433634 |
| VL0583 | 1 | 0.033433634 |
| VL0584 | 1 | 0.033433634 |
| VL0585 | 1 | 0.033433634 |
| VL0586 | 1 | 0.033433634 |
| VL0587 | 1 | 0.033433634 |
| VL0588 | 1 | 0.033433634 |
| VL0589 | 1 | 0.033433634 |
| VL0590 | 1 | 0.033433634 |
| VL0591 | 1 | 0.033433634 |
| VL0592 | 1 | 0.033433634 |
| VL0593 | 1 | 0.033433634 |
| VL0594 | 1 | 0.033433634 |
| VL0595 | 1 | 0.033433634 |
| VL0596 | 1 | 0.033433634 |
| VL0597 | 1 | 0.033433634 |
| VL0598 | 1 | 0.033433634 |
| VL0599 | 1 | 0.033433634 |
| VL0600 | 1 | 0.033433634 |
| VL0601 | 1 | 0.033433634 |
| VL0602 | 1 | 0.033433634 |
| VL0603 | 1 | 0.033433634 |
| VL0604 | 1 | 0.033433634 |
| VL0605 | 1 | 0.033433634 |
| VL0606 | 1 | 0.033433634 |
| VL0607 | 1 | 0.033433634 |
| VL0608 | 1 | 0.033433634 |
| VL0609 | 1 | 0.033433634 |
| VL0610 | 1 | 0.033433634 |
| VL0611 | 1 | 0.033433634 |
| VL0612 | 1 | 0.033433634 |
| VL0613 | 1 | 0.033433634 |
| VL0614 | 1 | 0.033433634 |
| VL0615 | 1 | 0.033433634 |
| VL0616 | 1 | 0.033433634 |
| VL0617 | 1 | 0.033433634 |
| VL0618 | 1 | 0.033433634 |
| VL0619 | 1 | 0.033433634 |
| VL0620 | 1 | 0.033433634 |
| VL0621 | 1 | 0.033433634 |
| VL0622 | 1 | 0.033433634 |
| VL0623 | 1 | 0.033433634 |
| VL0624 | 1 | 0.033433634 |
| VL0625 | 1 | 0.033433634 |
| VL0626 | 1 | 0.033433634 |
| VL0627 | 1 | 0.033433634 |
| VL0628 | 1 | 0.033433634 |
| VL0629 | 1 | 0.033433634 |
| VL0630 | 1 | 0.033433634 |
| VL0631 | 1 | 0.033433634 |
| VL0632 | 1 | 0.033433634 |
| VL0633 | 1 | 0.033433634 |
| VL0634 | 1 | 0.033433634 |
| VL0635 | 1 | 0.033433634 |
| VL0636 | 1 | 0.033433634 |
| VL0637 | 1 | 0.033433634 |
| VL0638 | 1 | 0.033433634 |
| VL0639 | 1 | 0.033433634 |
| VL0640 | 1 | 0.033433634 |
| VL0641 | 1 | 0.033433634 |
| VL0642 | 1 | 0.033433634 |
| VL0643 | 1 | 0.033433634 |
| VL0644 | 1 | 0.033433634 |
| VL0645 | 1 | 0.033433634 |
| VL0646 | 1 | 0.033433634 |
| VL0647 | 1 | 0.033433634 |
| VL0648 | 1 | 0.033433634 |
| VL0649 | 1 | 0.033433634 |
| VL0650 | 1 | 0.033433634 |
| VL0651 | 1 | 0.033433634 |
| VL0652 | 1 | 0.033433634 |
| VL0653 | 1 | 0.033433634 |
| VL0654 | 1 | 0.033433634 |
| VL0655 | 1 | 0.033433634 |
| VL0656 | 1 | 0.033433634 |
| VL0657 | 1 | 0.033433634 |
| VL0658 | 1 | 0.033433634 |
| VL0659 | 1 | 0.033433634 |
| VL0660 | 1 | 0.033433634 |
| VL0661 | 1 | 0.033433634 |
| VL0662 | 1 | 0.033433634 |
| VL0663 | 1 | 0.033433634 |
| VL0664 | 1 | 0.033433634 |
| VL0665 | 1 | 0.033433634 |
| VL0666 | 1 | 0.033433634 |
| VL0667 | 1 | 0.033433634 |
| VL0668 | 1 | 0.033433634 |
| VL0669 | 1 | 0.033433634 |
| VL0670 | 1 | 0.033433634 |
| VL0671 | 1 | 0.033433634 |
| VL0672 | 1 | 0.033433634 |
| VL0673 | 1 | 0.033433634 |
| VL0674 | 1 | 0.033433634 |
| VL0675 | 1 | 0.033433634 |
| VL0676 | 1 | 0.033433634 |
| VL0677 | 1 | 0.033433634 |
| VL0678 | 1 | 0.033433634 |
| VL0679 | 1 | 0.033433634 |
| VL0680 | 1 | 0.033433634 |
| VL0681 | 1 | 0.033433634 |
| VL0682 | 1 | 0.033433634 |
| VL0683 | 1 | 0.033433634 |
| VL0684 | 1 | 0.033433634 |
| VL0685 | 1 | 0.033433634 |
| VL0686 | 1 | 0.033433634 |
| VL0687 | 1 | 0.033433634 |
| VL0688 | 1 | 0.033433634 |
| VL0689 | 1 | 0.033433634 |
| VL0690 | 1 | 0.033433634 |
| VL0691 | 1 | 0.033433634 |
| VL0692 | 1 | 0.033433634 |
| VL0693 | 1 | 0.033433634 |
| VL0694 | 1 | 0.033433634 |
| VL0695 | 1 | 0.033433634 |
| VL0696 | 1 | 0.033433634 |
| VL0697 | 1 | 0.033433634 |
| VL0698 | 1 | 0.033433634 |
| VL0699 | 1 | 0.033433634 |
| VL0700 | 1 | 0.033433634 |
| VL0701 | 1 | 0.033433634 |
| VL0702 | 1 | 0.033433634 |
| VL0703 | 1 | 0.033433634 |
| VL0704 | 1 | 0.033433634 |
| VL0705 | 1 | 0.033433634 |
| VL0706 | 1 | 0.033433634 |
| VL0707 | 1 | 0.033433634 |
| VL0708 | 1 | 0.033433634 |
| VL0709 | 1 | 0.033433634 |
| VL0710 | 1 | 0.033433634 |
| VL0711 | 1 | 0.033433634 |
| VL0712 | 1 | 0.033433634 |
| VL0713 | 1 | 0.033433634 |
| VL0714 | 1 | 0.033433634 |
| VL0715 | 1 | 0.033433634 |
| VL0716 | 1 | 0.033433634 |
| VL0717 | 1 | 0.033433634 |
| VL0718 | 1 | 0.033433634 |
| VL0719 | 1 | 0.033433634 |
| VL0720 | 1 | 0.033433634 |
| VL0721 | 1 | 0.033433634 |
| VL0722 | 1 | 0.033433634 |
| VL0723 | 1 | 0.033433634 |
| VL0724 | 1 | 0.033433634 |
| VL0725 | 1 | 0.033433634 |
| VL0726 | 1 | 0.033433634 |
| VL0727 | 1 | 0.033433634 |
| VL0728 | 1 | 0.033433634 |
| VL0729 | 1 | 0.033433634 |
| VL0730 | 1 | 0.033433634 |
| VL0731 | 1 | 0.033433634 |
| VL0732 | 1 | 0.033433634 |
| VL0733 | 1 | 0.033433634 |
| VL0734 | 1 | 0.033433634 |
| VL0735 | 1 | 0.033433634 |
| VL0736 | 1 | 0.033433634 |
| VL0737 | 1 | 0.033433634 |
| VL0738 | 1 | 0.033433634 |
| VL0739 | 1 | 0.033433634 |
| VL0740 | 1 | 0.033433634 |
| VL0741 | 1 | 0.033433634 |
| VL0742 | 1 | 0.033433634 |
| VL0743 | 1 | 0.033433634 |
| VL0744 | 1 | 0.033433634 |
| VL0745 | 1 | 0.033433634 |
| VL0746 | 1 | 0.033433634 |
| VL0747 | 1 | 0.033433634 |
| VL0748 | 1 | 0.033433634 |
| VL0749 | 1 | 0.033433634 |
| VL0750 | 1 | 0.033433634 |
| VL0751 | 1 | 0.033433634 |
| VL0752 | 1 | 0.033433634 |
| VL0753 | 1 | 0.033433634 |
| VL0754 | 1 | 0.033433634 |
| VL0755 | 1 | 0.033433634 |
| VL0756 | 1 | 0.033433634 |
| VL0757 | 1 | 0.033433634 |
| VL0758 | 1 | 0.033433634 |
| VL0759 | 1 | 0.033433634 |
| VL0760 | 1 | 0.033433634 |
| VL0761 | 1 | 0.033433634 |
| VL0762 | 1 | 0.033433634 |
| VL0763 | 1 | 0.033433634 |
| VL0764 | 1 | 0.033433634 |
| VL0765 | 1 | 0.033433634 |
| VL0766 | 1 | 0.033433634 |
| VL0767 | 1 | 0.033433634 |
| VL0768 | 1 | 0.033433634 |
| VL0769 | 1 | 0.033433634 |
| VL0770 | 1 | 0.033433634 |
| VL0771 | 1 | 0.033433634 |
| VL0772 | 1 | 0.033433634 |
| VL0773 | 1 | 0.033433634 |
| VL0774 | 1 | 0.033433634 |
| VL0775 | 1 | 0.033433634 |
| VL0776 | 1 | 0.033433634 |
| VL0777 | 1 | 0.033433634 |
| VL0778 | 1 | 0.033433634 |
| VL0779 | 1 | 0.033433634 |
| VL0780 | 1 | 0.033433634 |
| VL0781 | 1 | 0.033433634 |
| VL0782 | 1 | 0.033433634 |
| VL0783 | 1 | 0.033433634 |
| VL0784 | 1 | 0.033433634 |
| VL0785 | 1 | 0.033433634 |
| VL0786 | 1 | 0.033433634 |
| VL0787 | 1 | 0.033433634 |
| VL0788 | 1 | 0.033433634 |
| VL0789 | 1 | 0.033433634 |
| VL0790 | 1 | 0.033433634 |
| VL0791 | 1 | 0.033433634 |
| VL0792 | 1 | 0.033433634 |
| VL0793 | 1 | 0.033433634 |
| VL0794 | 1 | 0.033433634 |
| VL0795 | 1 | 0.033433634 |
| VL0796 | 1 | 0.033433634 |
| VL0797 | 1 | 0.033433634 |
| VL0798 | 1 | 0.033433634 |
| VL0799 | 1 | 0.033433634 |
| VL0800 | 1 | 0.033433634 |
| VL0801 | 1 | 0.033433634 |
| VL0802 | 1 | 0.033433634 |
| VL0803 | 1 | 0.033433634 |
| VL0804 | 1 | 0.033433634 |
| VL0805 | 1 | 0.033433634 |
| VL0806 | 1 | 0.033433634 |
| VL0807 | 1 | 0.033433634 |
| VL0808 | 1 | 0.033433634 |
| VL0809 | 1 | 0.033433634 |
| VL0810 | 1 | 0.033433634 |
| VL0811 | 1 | 0.033433634 |
| VL0812 | 1 | 0.033433634 |
| VL0813 | 1 | 0.033433634 |
| VL0814 | 1 | 0.033433634 |
| VL0815 | 1 | 0.033433634 |
| VL0816 | 1 | 0.033433634 |
| VL0817 | 1 | 0.033433634 |
| VL0818 | 1 | 0.033433634 |
| VL0819 | 1 | 0.033433634 |
| VL0820 | 1 | 0.033433634 |
| VL0821 | 1 | 0.033433634 |
| VL0822 | 1 | 0.033433634 |
| VL0823 | 1 | 0.033433634 |
| VL0824 | 1 | 0.033433634 |
| VL0825 | 1 | 0.033433634 |
| VL0826 | 1 | 0.033433634 |
| VL0827 | 1 | 0.033433634 |
| VL0828 | 1 | 0.033433634 |
| VL0829 | 1 | 0.033433634 |
| VL0830 | 1 | 0.033433634 |
| VL0831 | 1 | 0.033433634 |
| VL0832 | 1 | 0.033433634 |
| VL0833 | 1 | 0.033433634 |
| VL0834 | 1 | 0.033433634 |
| VL0835 | 1 | 0.033433634 |
| VL0836 | 1 | 0.033433634 |
| VL0837 | 1 | 0.033433634 |
| VL0838 | 1 | 0.033433634 |
| VL0839 | 1 | 0.033433634 |
| VL0840 | 1 | 0.033433634 |
| VL0841 | 1 | 0.033433634 |
| VL0842 | 1 | 0.033433634 |
| VL0843 | 1 | 0.033433634 |
| VL0844 | 1 | 0.033433634 |
| VL0845 | 1 | 0.033433634 |
| VL0846 | 1 | 0.033433634 |
| VL0847 | 1 | 0.033433634 |
| VL0848 | 1 | 0.033433634 |
| VL0849 | 1 | 0.033433634 |
| VL0850 | 1 | 0.033433634 |
| VL0851 | 1 | 0.033433634 |
| VL0852 | 1 | 0.033433634 |
| VL0853 | 1 | 0.033433634 |
| VL0854 | 1 | 0.033433634 |
| VL0855 | 1 | 0.033433634 |
| VL0856 | 1 | 0.033433634 |
| VL0857 | 1 | 0.033433634 |
| VL0858 | 1 | 0.033433634 |
| VL0859 | 1 | 0.033433634 |
| VL0860 | 1 | 0.033433634 |
| VL0861 | 1 | 0.033433634 |
| VL0862 | 1 | 0.033433634 |
| VL0863 | 1 | 0.033433634 |
| VL0864 | 1 | 0.033433634 |
| VL0865 | 1 | 0.033433634 |
| VL0866 | 1 | 0.033433634 |
| VL0867 | 1 | 0.033433634 |
| VL0868 | 1 | 0.033433634 |
| VL0869 | 1 | 0.033433634 |
| VL0870 | 1 | 0.033433634 |
| VL0871 | 1 | 0.033433634 |
| VL0872 | 1 | 0.033433634 |
| VL0873 | 1 | 0.033433634 |
| VL0874 | 1 | 0.033433634 |
| VL0875 | 1 | 0.033433634 |
| VL0876 | 1 | 0.033433634 |
| VL0877 | 1 | 0.033433634 |
| VL0878 | 1 | 0.033433634 |
| VL0879 | 1 | 0.033433634 |
| VL0880 | 1 | 0.033433634 |
| VL0881 | 1 | 0.033433634 |
| VL0882 | 1 | 0.033433634 |
| VL0883 | 1 | 0.033433634 |
| VL0884 | 1 | 0.033433634 |
| VL0885 | 1 | 0.033433634 |
| VL0886 | 1 | 0.033433634 |
| VL0887 | 1 | 0.033433634 |
| VL0888 | 1 | 0.033433634 |
| VL0889 | 1 | 0.033433634 |
| VL0890 | 1 | 0.033433634 |
| VL0891 | 1 | 0.033433634 |
| VL0892 | 1 | 0.033433634 |
| VL0893 | 1 | 0.033433634 |
| VL0894 | 1 | 0.033433634 |
| VL0895 | 1 | 0.033433634 |
| VL0896 | 1 | 0.033433634 |
| VL0897 | 1 | 0.033433634 |
| VL0898 | 1 | 0.033433634 |
| VL0899 | 1 | 0.033433634 |
| VL0900 | 1 | 0.033433634 |
| VL0901 | 1 | 0.033433634 |
| VL0902 | 1 | 0.033433634 |
| VL0903 | 1 | 0.033433634 |
| VL0904 | 1 | 0.033433634 |
| VL0905 | 1 | 0.033433634 |
| VL0906 | 1 | 0.033433634 |
| VL0907 | 1 | 0.033433634 |
| VL0908 | 1 | 0.033433634 |
| VL0909 | 1 | 0.033433634 |
| VL0910 | 1 | 0.033433634 |
| VL0911 | 1 | 0.033433634 |
| VL0912 | 1 | 0.033433634 |
| VL0913 | 1 | 0.033433634 |
| VL0914 | 1 | 0.033433634 |
| VL0915 | 1 | 0.033433634 |
| VL0916 | 1 | 0.033433634 |
| VL0917 | 1 | 0.033433634 |
| VL0918 | 1 | 0.033433634 |
| VL0919 | 1 | 0.033433634 |
| VL0920 | 1 | 0.033433634 |
| VL0921 | 1 | 0.033433634 |
| VL0922 | 1 | 0.033433634 |
| VL0923 | 1 | 0.033433634 |
| VL0924 | 1 | 0.033433634 |
| VL0925 | 1 | 0.033433634 |
| VL0926 | 1 | 0.033433634 |
| VL0927 | 1 | 0.033433634 |
| VL0928 | 1 | 0.033433634 |
| VL0929 | 1 | 0.033433634 |
| VL0930 | 1 | 0.033433634 |
| VL0931 | 1 | 0.033433634 |
| VL0932 | 1 | 0.033433634 |
| VL0933 | 1 | 0.033433634 |
| VL0934 | 1 | 0.033433634 |
| VL0935 | 1 | 0.033433634 |
| VL0936 | 1 | 0.033433634 |
| VL0937 | 1 | 0.033433634 |
| VL0938 | 1 | 0.033433634 |
| VL0939 | 1 | 0.033433634 |
| VL0940 | 1 | 0.033433634 |
| VL0941 | 1 | 0.033433634 |
| VL0942 | 1 | 0.033433634 |
| VL0943 | 1 | 0.033433634 |
| VL0944 | 1 | 0.033433634 |
| VL0945 | 1 | 0.033433634 |
| VL0946 | 1 | 0.033433634 |
| VL0947 | 1 | 0.033433634 |
| VL0948 | 1 | 0.033433634 |
| VL0949 | 1 | 0.033433634 |
| VL0950 | 1 | 0.033433634 |
| VL0951 | 1 | 0.033433634 |
| VL0952 | 1 | 0.033433634 |
| VL0953 | 1 | 0.033433634 |
| VL0954 | 1 | 0.033433634 |
| VL0955 | 1 | 0.033433634 |
| VL0956 | 1 | 0.033433634 |
| VL0957 | 1 | 0.033433634 |
| VL0958 | 1 | 0.033433634 |
| VL0959 | 1 | 0.033433634 |
| VL0960 | 1 | 0.033433634 |
| VL0961 | 1 | 0.033433634 |
| VL0962 | 1 | 0.033433634 |
| VL0963 | 1 | 0.033433634 |
| VL0964 | 1 | 0.033433634 |
| VL0965 | 1 | 0.033433634 |
| VL0966 | 1 | 0.033433634 |
| VL0967 | 1 | 0.033433634 |
| VL0968 | 1 | 0.033433634 |
| VL0969 | 1 | 0.033433634 |
| VL0970 | 1 | 0.033433634 |
| VL0971 | 1 | 0.033433634 |
| VL0972 | 1 | 0.033433634 |
| VL0973 | 1 | 0.033433634 |
| VL0974 | 1 | 0.033433634 |
| VL0975 | 1 | 0.033433634 |
| VL0976 | 1 | 0.033433634 |
| VL0977 | 1 | 0.033433634 |
| VL0978 | 1 | 0.033433634 |
| VL0979 | 1 | 0.033433634 |
| VL0980 | 1 | 0.033433634 |
| VL0981 | 1 | 0.033433634 |
| VL0982 | 1 | 0.033433634 |
| VL0983 | 1 | 0.033433634 |
| VL0984 | 1 | 0.033433634 |
| VL0985 | 1 | 0.033433634 |
| VL0986 | 1 | 0.033433634 |
| VL0987 | 1 | 0.033433634 |
| VL0988 | 1 | 0.033433634 |
| VL0989 | 1 | 0.033433634 |
| VL0990 | 1 | 0.033433634 |
| VL0991 | 1 | 0.033433634 |
| VL0992 | 1 | 0.033433634 |
| VL0993 | 1 | 0.033433634 |
| VL0994 | 1 | 0.033433634 |
| VL0995 | 1 | 0.033433634 |
| VL0996 | 1 | 0.033433634 |
| VL0997 | 1 | 0.033433634 |
| VL0998 | 1 | 0.033433634 |
| VL0999 | 1 | 0.033433634 |
| VL1000 | 1 | 0.033433634 |
| VL1001 | 1 | 0.033433634 |
| VL1002 | 1 | 0.033433634 |
| VL1003 | 1 | 0.033433634 |
| VL1004 | 1 | 0.033433634 |
| VL1005 | 1 | 0.033433634 |
| VL1006 | 1 | 0.033433634 |
| VL1007 | 1 | 0.033433634 |
| VL1008 | 1 | 0.033433634 |
| VL1009 | 1 | 0.033433634 |
| VL1010 | 1 | 0.033433634 |
| VL1011 | 1 | 0.033433634 |
| VL1012 | 1 | 0.033433634 |
| VL1013 | 1 | 0.033433634 |
| VL1014 | 1 | 0.033433634 |
| VL1015 | 1 | 0.033433634 |
| VL1016 | 1 | 0.033433634 |
| VL1017 | 1 | 0.033433634 |
| VL1018 | 1 | 0.033433634 |
| VL1019 | 1 | 0.033433634 |
| VL1020 | 1 | 0.033433634 |
| VL1021 | 1 | 0.033433634 |
| VL1022 | 1 | 0.033433634 |
| VL1023 | 1 | 0.033433634 |
| VL1024 | 1 | 0.033433634 |
| VL1025 | 1 | 0.033433634 |
| VL1026 | 1 | 0.033433634 |
| VL1027 | 1 | 0.033433634 |
| VL1028 | 1 | 0.033433634 |
| VL1029 | 1 | 0.033433634 |
| VL1030 | 1 | 0.033433634 |
| VL1031 | 1 | 0.033433634 |
| VL1032 | 1 | 0.033433634 |
| VL1033 | 1 | 0.033433634 |
| VL1034 | 1 | 0.033433634 |
| VL1035 | 1 | 0.033433634 |
| VL1036 | 1 | 0.033433634 |
| VL1037 | 1 | 0.033433634 |
| VL1038 | 1 | 0.033433634 |
| VL1039 | 1 | 0.033433634 |
| VL1040 | 1 | 0.033433634 |
| VL1041 | 1 | 0.033433634 |
| VL1042 | 1 | 0.033433634 |
| VL1043 | 1 | 0.033433634 |
| VL1044 | 1 | 0.033433634 |
| VL1045 | 1 | 0.033433634 |
| VL1046 | 1 | 0.033433634 |
| VL1047 | 1 | 0.033433634 |
| VL1048 | 1 | 0.033433634 |
| VL1049 | 1 | 0.033433634 |
| VL1050 | 1 | 0.033433634 |
| VL1051 | 1 | 0.033433634 |
| VL1052 | 1 | 0.033433634 |
| VL1053 | 1 | 0.033433634 |
| VL1054 | 1 | 0.033433634 |
| VL1055 | 1 | 0.033433634 |
| VL1056 | 1 | 0.033433634 |
| VL1057 | 1 | 0.033433634 |
| VL1058 | 1 | 0.033433634 |
| VL1059 | 1 | 0.033433634 |
| VL1060 | 1 | 0.033433634 |
| VL1061 | 1 | 0.033433634 |
| VL1062 | 1 | 0.033433634 |
| VL1063 | 1 | 0.033433634 |
| VL1064 | 1 | 0.033433634 |
| VL1065 | 1 | 0.033433634 |
| VL1066 | 1 | 0.033433634 |
| VL1067 | 1 | 0.033433634 |
| VL1068 | 1 | 0.033433634 |
| VL1069 | 1 | 0.033433634 |
| VL1070 | 1 | 0.033433634 |
| VL1071 | 1 | 0.033433634 |
| VL1072 | 1 | 0.033433634 |
| VL1073 | 1 | 0.033433634 |
| VL1074 | 1 | 0.033433634 |
| VL1075 | 1 | 0.033433634 |
| VL1076 | 1 | 0.033433634 |
| VL1077 | 1 | 0.033433634 |
| VL1078 | 1 | 0.033433634 |
| VL1079 | 1 | 0.033433634 |
| VL1080 | 1 | 0.033433634 |
| VL1081 | 1 | 0.033433634 |
| VL1082 | 1 | 0.033433634 |
| VL1083 | 1 | 0.033433634 |
| VL1084 | 1 | 0.033433634 |
| VL1085 | 1 | 0.033433634 |
| VL1086 | 1 | 0.033433634 |
| VL1087 | 1 | 0.033433634 |
| VL1088 | 1 | 0.033433634 |
| VL1089 | 1 | 0.033433634 |
| VL1090 | 1 | 0.033433634 |
| VL1091 | 1 | 0.033433634 |
| VL1092 | 1 | 0.033433634 |
| VL1093 | 1 | 0.033433634 |
| VL1094 | 1 | 0.033433634 |
| VL1095 | 1 | 0.033433634 |
| VL1096 | 1 | 0.033433634 |
| VL1097 | 1 | 0.033433634 |
| VL1098 | 1 | 0.033433634 |
| VL1099 | 1 | 0.033433634 |
| VL1100 | 1 | 0.033433634 |
| VL1101 | 1 | 0.033433634 |
| VL1102 | 1 | 0.033433634 |
| VL1103 | 1 | 0.033433634 |
| VL1104 | 1 | 0.033433634 |
| VL1105 | 1 | 0.033433634 |
| VL1106 | 1 | 0.033433634 |
| VL1107 | 1 | 0.033433634 |
| VL1108 | 1 | 0.033433634 |
| VL1109 | 1 | 0.033433634 |
| VL1110 | 1 | 0.033433634 |
| VL1111 | 1 | 0.033433634 |
| VL1112 | 1 | 0.033433634 |
| VL1113 | 1 | 0.033433634 |
| VL1114 | 1 | 0.033433634 |
| VL1115 | 1 | 0.033433634 |
| VL1116 | 1 | 0.033433634 |
| VL1117 | 1 | 0.033433634 |
| VL1118 | 1 | 0.033433634 |
| VL1119 | 1 | 0.033433634 |
| VL1120 | 1 | 0.033433634 |
| VL1121 | 1 | 0.033433634 |
| VL1122 | 1 | 0.033433634 |
| VL1123 | 1 | 0.033433634 |
| VL1124 | 1 | 0.033433634 |
| VL1125 | 1 | 0.033433634 |
| VL1126 | 1 | 0.033433634 |
| VL1127 | 1 | 0.033433634 |
| VL1128 | 1 | 0.033433634 |
| VL1129 | 1 | 0.033433634 |
| VL1130 | 1 | 0.033433634 |

**Table S43. Classification of Ty1/Copia and Ty3/Gypsy families**

| Super family |  | Number | Percentage (%) |
| --- | --- | --- | --- |
| Ty1/Copia | Ale | 115 | 33.14 |
|  | Angela | 49 | 14.12 |
|  | Bianca | 8 | 2.31 |
|  | Horpia2 | 14 | 4.03 |
|  | Lkya | 10 | 2.88 |
|  | Owis | 91 | 26.22 |
|  | Rare1 | 10 | 2.88 |
|  | Rare2 | 50 | 14.41 |
| Ty3/Gypsy | Bagy2 | 50 | 8.04 |
|  | Cereba | 40 | 6.43 |
|  | Dagan | 373 | 59.97 |
|  | Erika | 6 | 0.96 |
|  | GA | 4 | 0.64 |
|  | Geneva | 73 | 11.74 |
|  | Laura | 6 | 0.96 |
|  | Retrosat | 70 | 11.25 |

**Table S44. The percentage of reads derived from LTR retrotransposons measured by RNA-seq from six tissues in tung tree**

| Sample | Total reads | Mapped reads (%) | LTR | | Gypsy | | Copia | | Other LTR RT | |
| --- | --- | --- | --- | --- | --- | --- | --- | --- | --- | --- |
|  |  |  | Reads | Percentage | Reads | Percentage | Reads | Percentage | Reads | Percentage |
| Root | 64,936,708 | 91.67 | 41718 | 0.0642% | 7430 | 0.0194% | 12626 | 0.0114% | 21662 | 0.0334% |
| Stem | 50,995,280 | 79.44 | 30285 | 0.0594% | 3421 | 0.0203% | 10335 | 0.0067% | 16529 | 0.0324% |
| Leaf | 76,372,384 | 89.94 | 41669 | 0.0546% | 4733 | 0.0254% | 19375 | 0.0062% | 17561 | 0.0230% |
| Female flower | 74,438,216 | 90.57 | 69533 | 0.0934% | 5210 | 0.0237% | 17610 | 0.0070% | 46713 | 0.0628% |
| Male flower | 63,252,746 | 88.61 | 59982 | 0.0948% | 5040 | 0.0300% | 18994 | 0.0080% | 35948 | 0.0568% |
| Seed | 789,108,408 | 95.95 | 449631 | 0.0173% | 129526 | 0.0164% | 136577 | 0.0233% | 183528 | 0.0570% |

**Table S45. Expression quantity (FPKM value) of the top 29 tung tree LTR families highly expressed in seed**

| Gene | Root | Stem | Leaf | Female flower | Male flower | Seed |
| --- | --- | --- | --- | --- | --- | --- |
| VL0631 | 8.092319 | 8.006303 | 6.414458 | 1.706871 | 5.312487 | 11.08873 |
| VL0883 | 2.209019 | 2.624494 | 5.973387 | 5.386334 | 5.130156 | 7.22597 |
| VL0861 | 1.883799 | 1.8369 | 2.148359 | 2.253322 | 2.129104 | 5.863583 |
| VL0496 | 0.728858 | 0.730687 | 1.933841 | 1.143417 | 0.780329 | 3.269679 |
| VL0088 | 0.58881 | 0.888904 | 0.986239 | 1.034833 | 1.173811 | 3.120569 |
| VL0552 | 0.918293 | 0.925285 | 1.686557 | 0.871957 | 1.764536 | 2.9408 |
| VL0826 | 0.254741 | 0.522089 | 0.521843 | 2.50044 | 2.9506 | 2.636805 |
| VL0150 | 1.580628 | 2.054055 | 2.050102 | 0.648338 | 0.531715 | 2.619298 |
| VL0835 | 0.918091 | 1.421359 | 6.239324 | 0 | 0.016514 | 2.447857 |
| VL0520 | 0.137844 | 22.16072 | 4.908813 | 38.10004 | 5.935157 | 2.382111 |
| VL1071 | 2.438324 | 2.098244 | 1.679527 | 1.96772 | 1.670929 | 2.354918 |
| VL0782 | 0.008761 | 0.048932 | 0 | 0.075398 | 0 | 2.219964 |
| VL0460 | 0.799925 | 0.650949 | 0.811581 | 0.624721 | 0.008957 | 2.135695 |
| VL0188 | 1.13376 | 1.125293 | 1.998828 | 1.406752 | 1.544601 | 1.630482 |
| VL0012 | 2.136787 | 1.559183 | 0.605148 | 0.170329 | 0.155143 | 1.587106 |
| VL0524 | 0.44412 | 0.362364 | 0.331538 | 0.4539 | 0.166061 | 1.452035 |
| VL0236 | 0.360006 | 0.363647 | 0.75151 | 0.541554 | 0.640357 | 1.401164 |
| VL0848 | 0.236198 | 0.384418 | 0.752909 | 0.446161 | 0.966846 | 1.375178 |
| VL0228 | 0.411259 | 0.148656 | 0.602734 | 1.032288 | 0.977943 | 1.374741 |
| VL0003 | 0.219942 | 0.176796 | 0.185389 | 0.058865 | 0.064364 | 1.362043 |
| VL1127 | 17.07058 | 10.77815 | 0.151209 | 0.235101 | 0.048466 | 1.306228 |
| VL1026 | 2.377122 | 2.809219 | 4.572779 | 79.75539 | 115.9007 | 1.237784 |
| VL0276 | 0.00115 | 0.176157 | 2.926163 | 0.337224 | 0.073071 | 1.218812 |
| VL1113 | 0.954345 | 0.741692 | 1.215123 | 0.629325 | 0.884435 | 1.190684 |
| VL0107 | 0.779902 | 1.33176 | 1.306248 | 2.333978 | 2.805971 | 1.189859 |
| VL0562 | 0.580446 | 0.407536 | 0.311459 | 0.176183 | 0.17664 | 1.169551 |
| VL0377 | 2.206839 | 2.500936 | 1.86184 | 3.710045 | 4.156882 | 1.064139 |
| VL0703 | 0.848481 | 3.126913 | 1.454458 | 4.096578 | 1.107515 | 1.014236 |
| VL0105 | 0.682184 | 1.519819 | 1.845069 | 1.104327 | 1.054452 | 1.013158 |

**Table S46. Expression quantity (FPKM value) of the top 21 tung tree LTR families highly expressed in root**

| gene | Root | Stem | Leaf | Female flower | Male flower | Seed |
| --- | --- | --- | --- | --- | --- | --- |
| VL1115 | 21.33359 | 7.457783 | 6.243737 | 6.065106 | 4.342405 | 0.12242 |
| VL1127 | 17.07058 | 10.77815 | 0.151209 | 0.235101 | 0.048466 | 1.306228 |
| VL0631 | 8.092319 | 8.006303 | 6.414458 | 1.706871 | 5.312487 | 11.08873 |
| VL0454 | 6.310483 | 3.795426 | 0.513461 | 0.31047 | 0.170469 | 0.954286 |
| VL0661 | 2.74045 | 1.578605 | 2.516884 | 5.676684 | 3.820723 | 0.203869 |
| VL1071 | 2.438324 | 2.098244 | 1.679527 | 1.96772 | 1.670929 | 2.354918 |
| VL1026 | 2.377122 | 2.809219 | 4.572779 | 79.75539 | 115.9007 | 1.237784 |
| VL0883 | 2.209019 | 2.624494 | 5.973387 | 5.386334 | 5.130156 | 7.22597 |
| VL0377 | 2.206839 | 2.500936 | 1.86184 | 3.710045 | 4.156882 | 1.064139 |
| VL0012 | 2.136787 | 1.559183 | 0.605148 | 0.170329 | 0.155143 | 1.587106 |
| VL0220 | 2.072145 | 1.675573 | 0.348069 | 0.348467 | 0.218255 | 0.842588 |
| VL0861 | 1.883799 | 1.8369 | 2.148359 | 2.253322 | 2.129104 | 5.863583 |
| VL0150 | 1.580628 | 2.054055 | 2.050102 | 0.648338 | 0.531715 | 2.619298 |
| VL0551 | 1.513242 | 0.351403 | 0.023586 | 0.041536 | 0.02113 | 0.013707 |
| VL0938 | 1.503375 | 0.14105 | 0.087448 | 0.050588 | 0.0766 | 0.233974 |
| VL0656 | 1.47918 | 0.350603 | 2.154012 | 0.006411 | 0.007419 | 0.025392 |
| VL0039 | 1.195776 | 0.880303 | 0.050356 | 1.928643 | 1.986594 | 0.030409 |
| VL0188 | 1.13376 | 1.125293 | 1.998828 | 1.406752 | 1.544601 | 1.630482 |
| VL0015 | 1.082185 | 0.423298 | 0.125069 | 0.045023 | 0.076431 | 0.220365 |
| VL0922 | 1.053553 | 0.017271 | 0.118915 | 0.79605 | 0 | 0.650855 |
| VL0912 | 1.028603 | 0.59906 | 0 | 0 | 0 | 0.702665 |

**Table S47. Expression quantity (FPKM value) of the top 23 tung tree LTR families highly expressed in stem**

| gene | Root | Stem | Leaf | Female flower | Male flower | Seed |
| --- | --- | --- | --- | --- | --- | --- |
| VL0520 | 0.137844 | 22.16072 | 4.908813 | 38.10004 | 5.935157 | 2.382111 |
| VL0259 | 0.912291 | 16.09971 | 5.661364 | 0.3524 | 0.150702 | 0.562424 |
| VL1127 | 17.07058 | 10.77815 | 0.151209 | 0.235101 | 0.048466 | 1.306228 |
| VL0631 | 8.092319 | 8.006303 | 6.414458 | 1.706871 | 5.312487 | 11.08873 |
| VL1115 | 21.33359 | 7.457783 | 6.243737 | 6.065106 | 4.342405 | 0.12242 |
| VL0454 | 6.310483 | 3.795426 | 0.513461 | 0.31047 | 0.170469 | 0.954286 |
| VL0703 | 0.848481 | 3.126913 | 1.454458 | 4.096578 | 1.107515 | 1.014236 |
| VL1026 | 2.377122 | 2.809219 | 4.572779 | 79.75539 | 115.9007 | 1.237784 |
| VL0883 | 2.209019 | 2.624494 | 5.973387 | 5.386334 | 5.130156 | 7.22597 |
| VL0377 | 2.206839 | 2.500936 | 1.86184 | 3.710045 | 4.156882 | 1.064139 |
| VL1071 | 2.438324 | 2.098244 | 1.679527 | 1.96772 | 1.670929 | 2.354918 |
| VL0150 | 1.580628 | 2.054055 | 2.050102 | 0.648338 | 0.531715 | 2.619298 |
| VL1028 | 0.313923 | 2.031457 | 0.985075 | 0.074385 | 0.733173 | 0.153328 |
| VL0861 | 1.883799 | 1.8369 | 2.148359 | 2.253322 | 2.129104 | 5.863583 |
| VL0220 | 2.072145 | 1.675573 | 0.348069 | 0.348467 | 0.218255 | 0.842588 |
| VL0661 | 2.74045 | 1.578605 | 2.516884 | 5.676684 | 3.820723 | 0.203869 |
| VL0012 | 2.136787 | 1.559183 | 0.605148 | 0.170329 | 0.155143 | 1.587106 |
| VL0105 | 0.682184 | 1.519819 | 1.845069 | 1.104327 | 1.054452 | 1.013158 |
| VL0321 | 0.858037 | 1.428517 | 0.4515 | 1.050623 | 1.260532 | 0.119997 |
| VL0835 | 0.918091 | 1.421359 | 6.239324 | 0 | 0.016514 | 2.447857 |
| VL0107 | 0.779902 | 1.33176 | 1.306248 | 2.333978 | 2.805971 | 1.189859 |
| VL0188 | 1.13376 | 1.125293 | 1.998828 | 1.406752 | 1.544601 | 1.630482 |
| VL0814 | 0.04675 | 1.095052 | 0 | 0 | 0.003295 | 0.01613 |

**Table S48. Expression quantity (FPKM value) of the top 28 tung tree LTR families highly expressed in leaf**

| gene | Root | Stem | Leaf | Female flower | Male flower | Seed |
| --- | --- | --- | --- | --- | --- | --- |
| VL0631 | 8.092319 | 8.006303 | 6.414458 | 1.706871 | 5.312487 | 11.08873 |
| VL1115 | 21.33359 | 7.457783 | 6.243737 | 6.065106 | 4.342405 | 0.12242 |
| VL0835 | 0.918091 | 1.421359 | 6.239324 | 0 | 0.016514 | 2.447857 |
| VL0883 | 2.209019 | 2.624494 | 5.973387 | 5.386334 | 5.130156 | 7.22597 |
| VL0259 | 0.912291 | 16.09971 | 5.661364 | 0.3524 | 0.150702 | 0.562424 |
| VL0520 | 0.137844 | 22.16072 | 4.908813 | 38.10004 | 5.935157 | 2.382111 |
| VL1026 | 2.377122 | 2.809219 | 4.572779 | 79.75539 | 115.9007 | 1.237784 |
| VL0276 | 0.00115 | 0.176157 | 2.926163 | 0.337224 | 0.073071 | 1.218812 |
| VL0661 | 2.74045 | 1.578605 | 2.516884 | 5.676684 | 3.820723 | 0.203869 |
| VL0656 | 1.47918 | 0.350603 | 2.154012 | 0.006411 | 0.007419 | 0.025392 |
| VL0861 | 1.883799 | 1.8369 | 2.148359 | 2.253322 | 2.129104 | 5.863583 |
| VL0150 | 1.580628 | 2.054055 | 2.050102 | 0.648338 | 0.531715 | 2.619298 |
| VL0188 | 1.13376 | 1.125293 | 1.998828 | 1.406752 | 1.544601 | 1.630482 |
| VL0496 | 0.728858 | 0.730687 | 1.933841 | 1.143417 | 0.780329 | 3.269679 |
| VL0377 | 2.206839 | 2.500936 | 1.86184 | 3.710045 | 4.156882 | 1.064139 |
| VL0105 | 0.682184 | 1.519819 | 1.845069 | 1.104327 | 1.054452 | 1.013158 |
| VL0552 | 0.918293 | 0.925285 | 1.686557 | 0.871957 | 1.764536 | 2.9408 |
| VL0626 | 0.089444 | 0 | 1.682331 | 0.344712 | 0.907399 | 0 |
| VL1071 | 2.438324 | 2.098244 | 1.679527 | 1.96772 | 1.670929 | 2.354918 |
| VL0014 | 0.335804 | 0.435906 | 1.554937 | 1.071195 | 0.940023 | 0.961424 |
| VL0703 | 0.848481 | 3.126913 | 1.454458 | 4.096578 | 1.107515 | 1.014236 |
| VL0279 | 0.047318 | 0.282033 | 1.34891 | 0.13856 | 0 | 0.397733 |
| VL0035 | 0.60895 | 0.858776 | 1.332527 | 0.637033 | 1.119235 | 0.992614 |
| VL0107 | 0.779902 | 1.33176 | 1.306248 | 2.333978 | 2.805971 | 1.189859 |
| VL0106 | 0.351077 | 0.6345 | 1.301602 | 0.669997 | 0.501688 | 0.883033 |
| VL0857 | 0.127112 | 0.174801 | 1.243526 | 0.526918 | 0.10229 | 0.956696 |
| VL1113 | 0.954345 | 0.741692 | 1.215123 | 0.629325 | 0.884435 | 1.190684 |
| VL0920 | 0 | 0.113396 | 1.082603 | 0.026823 | 0 | 0 |

**Table S49. Expression quantity (FPKM value) of the top 24 tung tree LTR families highly expressed in female flower**

| gene | Root | Stem | Leaf | Female flower | Male flower | Seed |
| --- | --- | --- | --- | --- | --- | --- |
| VL1026 | 2.377122 | 2.809219 | 4.572779 | 79.75539 | 115.9007 | 1.237784 |
| VL0520 | 0.137844 | 22.16072 | 4.908813 | 38.10004 | 5.935157 | 2.382111 |
| VL0098 | 0.258193 | 0.086624 | 0.066451 | 18.21615 | 33.78163 | 0.004591 |
| VL0584 | 0.746367 | 0.069567 | 0.114307 | 8.729215 | 14.60092 | 0.017624 |
| VL1115 | 21.33359 | 7.457783 | 6.243737 | 6.065106 | 4.342405 | 0.12242 |
| VL0661 | 2.74045 | 1.578605 | 2.516884 | 5.676684 | 3.820723 | 0.203869 |
| VL0883 | 2.209019 | 2.624494 | 5.973387 | 5.386334 | 5.130156 | 7.22597 |
| VL0221 | 0.088597 | 0.041333 | 0.403978 | 5.246941 | 2.591211 | 0.365582 |
| VL0703 | 0.848481 | 3.126913 | 1.454458 | 4.096578 | 1.107515 | 1.014236 |
| VL0377 | 2.206839 | 2.500936 | 1.86184 | 3.710045 | 4.156882 | 1.064139 |
| VL0826 | 0.254741 | 0.522089 | 0.521843 | 2.50044 | 2.9506 | 2.636805 |
| VL0107 | 0.779902 | 1.33176 | 1.306248 | 2.333978 | 2.805971 | 1.189859 |
| VL0861 | 1.883799 | 1.8369 | 2.148359 | 2.253322 | 2.129104 | 5.863583 |
| VL1071 | 2.438324 | 2.098244 | 1.679527 | 1.96772 | 1.670929 | 2.354918 |
| VL0039 | 1.195776 | 0.880303 | 0.050356 | 1.928643 | 1.986594 | 0.030409 |
| VL1088 | 0.056541 | 0.084124 | 0.302071 | 1.786303 | 2.38244 | 0.325862 |
| VL0631 | 8.092319 | 8.006303 | 6.414458 | 1.706871 | 5.312487 | 11.08873 |
| VL0188 | 1.13376 | 1.125293 | 1.998828 | 1.406752 | 1.544601 | 1.630482 |
| VL0496 | 0.728858 | 0.730687 | 1.933841 | 1.143417 | 0.780329 | 3.269679 |
| VL0105 | 0.682184 | 1.519819 | 1.845069 | 1.104327 | 1.054452 | 1.013158 |
| VL0014 | 0.335804 | 0.435906 | 1.554937 | 1.071195 | 0.940023 | 0.961424 |
| VL0321 | 0.858037 | 1.428517 | 0.4515 | 1.050623 | 1.260532 | 0.119997 |
| VL0088 | 0.58881 | 0.888904 | 0.986239 | 1.034833 | 1.173811 | 3.120569 |
| VL0228 | 0.411259 | 0.148656 | 0.602734 | 1.032288 | 0.977943 | 1.374741 |

**Table S50. Expression quantity (FPKM value) of the top 27 tung tree LTR families highly expressed in male flower**

| gene | Root | Stem | Leaf | Female flower | Male flower | Seed |
| --- | --- | --- | --- | --- | --- | --- |
| VL1026 | 2.377122 | 2.809219 | 4.572779 | 79.75539 | 115.9007 | 1.237784 |
| VL0098 | 0.258193 | 0.086624 | 0.066451 | 18.21615 | 33.78163 | 0.004591 |
| VL0584 | 0.746367 | 0.069567 | 0.114307 | 8.729215 | 14.60092 | 0.017624 |
| VL0520 | 0.137844 | 22.16072 | 4.908813 | 38.10004 | 5.935157 | 2.382111 |
| VL0631 | 8.092319 | 8.006303 | 6.414458 | 1.706871 | 5.312487 | 11.08873 |
| VL0883 | 2.209019 | 2.624494 | 5.973387 | 5.386334 | 5.130156 | 7.22597 |
| VL1115 | 21.33359 | 7.457783 | 6.243737 | 6.065106 | 4.342405 | 0.12242 |
| VL0377 | 2.206839 | 2.500936 | 1.86184 | 3.710045 | 4.156882 | 1.064139 |
| VL0661 | 2.74045 | 1.578605 | 2.516884 | 5.676684 | 3.820723 | 0.203869 |
| VL0826 | 0.254741 | 0.522089 | 0.521843 | 2.50044 | 2.9506 | 2.636805 |
| VL0107 | 0.779902 | 1.33176 | 1.306248 | 2.333978 | 2.805971 | 1.189859 |
| VL0221 | 0.088597 | 0.041333 | 0.403978 | 5.246941 | 2.591211 | 0.365582 |
| VL1088 | 0.056541 | 0.084124 | 0.302071 | 1.786303 | 2.38244 | 0.325862 |
| VL0861 | 1.883799 | 1.8369 | 2.148359 | 2.253322 | 2.129104 | 5.863583 |
| VL0039 | 1.195776 | 0.880303 | 0.050356 | 1.928643 | 1.986594 | 0.030409 |
| VL0552 | 0.918293 | 0.925285 | 1.686557 | 0.871957 | 1.764536 | 2.9408 |
| VL0666 | 0.132577 | 0.164701 | 0.018932 | 0.029776 | 1.700584 | 0.00065 |
| VL1071 | 2.438324 | 2.098244 | 1.679527 | 1.96772 | 1.670929 | 2.354918 |
| VL0188 | 1.13376 | 1.125293 | 1.998828 | 1.406752 | 1.544601 | 1.630482 |
| VL0321 | 0.858037 | 1.428517 | 0.4515 | 1.050623 | 1.260532 | 0.119997 |
| VL0769 | 0.004385 | 0.003187 | 0.003317 | 0.004415 | 1.241686 | 0.000311 |
| VL0088 | 0.58881 | 0.888904 | 0.986239 | 1.034833 | 1.173811 | 3.120569 |
| VL0035 | 0.60895 | 0.858776 | 1.332527 | 0.637033 | 1.119235 | 0.992614 |
| VL0703 | 0.848481 | 3.126913 | 1.454458 | 4.096578 | 1.107515 | 1.014236 |
| VL0924 | 0.006819 | 0.012854 | 0.001768 | 0.6544 | 1.06763 | 0.000242 |
| VL0105 | 0.682184 | 1.519819 | 1.845069 | 1.104327 | 1.054452 | 1.013158 |
| VL0234 | 0.382646 | 0.671551 | 0.455598 | 0.739313 | 1.010168 | 0.750374 |
