## Supplementary material for "The Tung Tree (*Vernicia Fordii*) Genome Provides A Resource for Understanding Genome Evolution and Oil Improvement": File S8

**File S8: Transcriptome sequencing, assembly and eFP browser**

A total of 17 samples of ‘Luxiputaotong’ were collected. The seeds were collected at 10, 15, 20, 25, and 30 WAF (weeks after flowering). Flowers were sampled at 30, 20, 10, and 1 DBF (days before flowering). Young leaves, stems, and roots were sampled in the spring. All samples were collected at 6:00 am-8:00 am. Three biological replicates were performed for seed and flower samples. Total RNA was extracted from each sample using Micro-to-Midi Total RNA Purification System according to the manufacture’s protocols (Life Technologies Carlsbad, CA, USA). Poly-A containing mRNA was purified from 2 µg of total RNA using oligo (dT) magnetic beads and fragmented into 200-500 bp pieces with divalent cations at 94°C for 5 min using a similar procedure for tung tree seed transcriptome cDNA library construction. The mRNA fragments were reverse-transcribed into first-strand cDNA using SuperScript II reverse transcriptase and random primers (Life Technologies). After double-stranded cDNA synthesis, fragments were end-repaired and A-tailed. The final cDNA library was created by purifying and enriching the double-stranded cDNA by PCR. The cDNA sequencing was performed using a paired-end flow cell with an Illumina HiSeq X Ten Sequencing System. After filtering, about 6 Gb clean paired-end data for each library were produced and aligned to the tung tree genome sequences using TopHat [1] and bowtie2 [2]. The FKPM values (fragments per kilobase of transcript per million mapped fragments) were generated for calculating gene expression level.

The tung tree eFP browser was set up as described by Winter et al [3]. We provided three intuitive modes for the gene expression level in tung tree eFP browser engine i.e. “Absolute”, “Relative” and “Compare”. In “Absolute”, the expression of each gene in each tissue was directly compared to the highest signal recorded for the given gene, with high levels of expression colored red and low levels colored yellow. The “Relative” mode displayed the ratio of a tissue’s expression level to appropriate control signal-typically the median or mock treatment. In the case of the Development Map and some of the other series we calculated the media value across all displayed samples for each probe set and loaded these into our database as a separate sample. In the other case, the appropriate untreated control data set value was used to calculate the relative value for the samples within a specified <group>. The output had tissues coloured with expression levels above the control signal value between yellow and blue. The “Compare” mode accepted two gene identifiers as inputs and compared the primary relative expression levels to the secondary in each tissue, using the same colour scheme described for relative. This is useful for identifying tissues in which one gene is more abundantly expressed relative to another.
