## Supplementary material for "The Tung Tree (*Vernicia Fordii*) Genome Provides A Resource for Understanding Genome Evolution and Oil Improvement": File S9

**File S9: Identification and expression of NBS-encoding gene families**

NBS-encoding genes in tung tree genome were identified using HMMER V 3.1 [1] search analysis to screen the predicted proteome against the raw hidden Markov model (HMM) corresponding to the Pfam NBS (NB-ARC) family domain. The TIR and LRR domains in the predicted NBS-encoding amino acid sequences were screened using HMMER search analysis against the HMM model Pfam TIR and LRR domains, respectively. CC motifs were analyzed using Paircoil2 [2] with a P-score cutoff of 0.025. To map the location of NBS genes in tung tree genome, the chromosomal distribution of NBS genes were generated by Mapinspect software according to their position given in the tung tree sequence [3]. The expression abundances of NBS genes in roots before and after Fusarium wilt infection were estimated according to Pertea et al [4]. Briefly, first, the HISAT2 package was applied to extract splice site and exon information from the tung tree genome annotation file and build a HISAT2 index. Then reads of each transcriptome were aligned to the tung tree genome. Second, samtools1.5 was used to sort and convert SAM files to binary BAM files. Finally, StringTie 1.3.3 was used to assemble reads to transcripts for each sample and transcripts from all samples were merged. Gene expression abundance in each sample was generated. The infection experiment design was described by Chen et al [5]. The original reads data for each transcriptome were downloaded at <https://www.ncbi.nlm.nih.gov/gds/> (SRR3374614, SRR3374616, SRR3374618, SRR3374620,	SRR3374622, SRR3374624, SRR3374626, SRR3374628, SRR3374630, SRR3374632, SRR3374634 and SRR3374636).
