## Supplementary material for "The Tung Tree (*Vernicia Fordii*) Genome Provides A Resource for Understanding Genome Evolution and Oil Improvement": File S10

**File S10: Lipid analysis and electron microscopy observation**

Seeds at different developmental stages including 10 weeks after flowering (WAF), 15 WAF, 20 WAF, 25 WAF, and 30 WAF were collected. Seed oil content and fatty acid composition were analyzed for each sample (three biological repeats) with the methods as described by Zhang et al [1]. Oil bodies were also observed with electron microscopy [2]. Tung seed endosperm tissues were cut into approximately 2×2 mm pieces and fixed for more than 2 h with 2.5% glutaraldehyde, 4% paraformaldehyde, and 0.1 M potassium phosphate, pH 7.0, at 4℃ for 24 h. The samples were washed with 0.1 M potassium phosphate buffer for 15 min three times and then treated with 1% OsO_4_, and 0.1 M potassium phosphate, pH7.0, at room temperature for 2-3 h. The fixed samples were rinsed with 0.1 M potassium phosphate buffer and dehydrated through an acetone series and embedded in Spurrs medium. Ultrathin sections (70 nm) were obtained with a Leica Reichert UltracutSorLeica EM UC6 ultramicrotome. Sections were stained with uranyl acetate and lead citrate and examined with a JEM1230 transmission electronmicroscope (JEOL, Japan) at 40-120 kV. Results showed that tung oil biosynthesis in the seeds started in mid-June (10 WAF), increased rapidly until September 31 (25 WAF) with the oil content of 59.22% (Figure 6C), and ended by mid-October. During the whole developmental stages, oleic acid (C18:1Δ9) accounted for minor percentage, whereas linoleic acid (C18:2Δ9,12) accounted for the major content (43%) at the initiation stage (10 and 15 WAF) and then gradually decreased in more mature seeds. Interestingly, accumulation of linoleic acid and α-ESA (α-C18:3Δ9,11,13) showed dramatically opposite patterns in the developing tung seeds (Figure 6C), which was partly because linoleic acid is the common substrate for synthesizing α-ESA and α-ALA (α-linolenic acid, C18:3Δ9,12,15). The α-ESA synthesis started after mid-July (15 WAF) and then increased rapidly up to 72.35% of seed oil following seed ripening (Figure 6C). The α-ALA was observed in mid-June (10 WAF) and accounted for minor percentage during the whole developmental stages, although it shared the same substrate with α-ESA. The oil is packed in subcellular structures called oil bodies or lipid droplets (Figure 6B). Tung seed oil droplets formed following the pattern of α-ESA accumulation in the seeds (Figure 6B and C). No visible oil droplet was observed in 10 WAF seeds and small oil droplets were observed in 15 WAF seeds. The number and sizes of oil droplets were dramatically increased in more mature seeds (20, 25, and 30 WAF).
