## Supplementary material for "The Tung Tree (*Vernicia Fordii*) Genome Provides A Resource for Understanding Genome Evolution and Oil Improvement": File S11

**File S11: Oil biosynthesis-related gene family identification and phyleogentic analysis**

To identify important genes in different species, six representative plant genomes (*V. fordii, J. curcas, R. communis, A. thaliana, S. indicum and G. max*) and predicted proteome sequences were obtained from available public databases. Among them, the *A. thaliana, G. max* and *R. communis* proteome sequences were downloaded from the Phytozome12 (https://phytozome.jgi.doe.gov/pz/portal.html) database, the *J. curcas* and *S. indicum* were derived from Jatropha genome database (<http://www.kazusa.or.jp/jatropha/>) and sinbase (<http://ocri-genomics.org/Sinbase/>), respectively. The tung tree proteome sequence was obtained in this study. Proteins with a PEPC domain (PF00311) were identified by the hidden Markov model-based HMMER program [1]. After local searches were performed in the proteome datasets with the PEPC domain, the resulting sequences were manually adjusted in multiple sequence alignments to correct obvious errors. Similarly, the OLE and KAS genes were also identified based on the oleosin domain (PF) and ketoacyl-synt domain (PF) by using the HMMER program [1]. The protein sequences of SAD, FAD, MAT, HAD, FAT, PP, PDCT, DAG-CPT, DGAT and PDAT of tung tree were used as queries for a local BLASTP search against these six plant proteome datasets with an e-value cut-off of less than 1e-5. Multiple sequence alignment was performed in MUSCLE using the default parameters [2]. ML trees were constructed using FastTree with the approximate likelihood ratio test (aLRT) method [3, 4] (Figures S26-S30).

**Supplementary Figures**

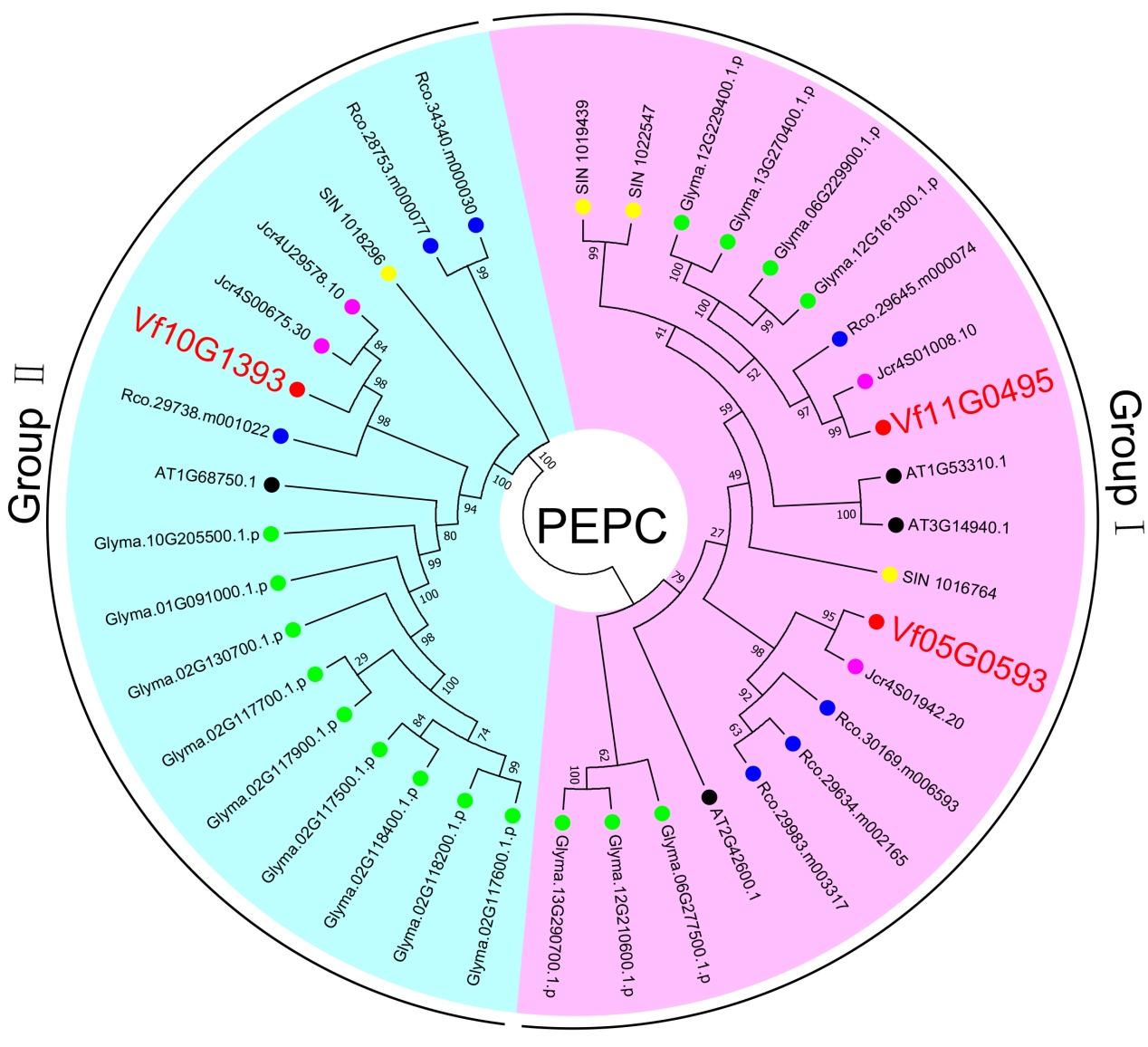

**Figure S26.** Phylogenetic analysis of PEPC genes. A maximum-likelihood phylogenetic tree constructed from protein sequences from *V. fordii* (red dots), *J. curcas* (pink dots), *S. indicum* (yellow dots), *R. communis* (blue dots), *G. max* (green dots) and *A. thaliana* (black dots). Different color represents different gene group generated from the tree.

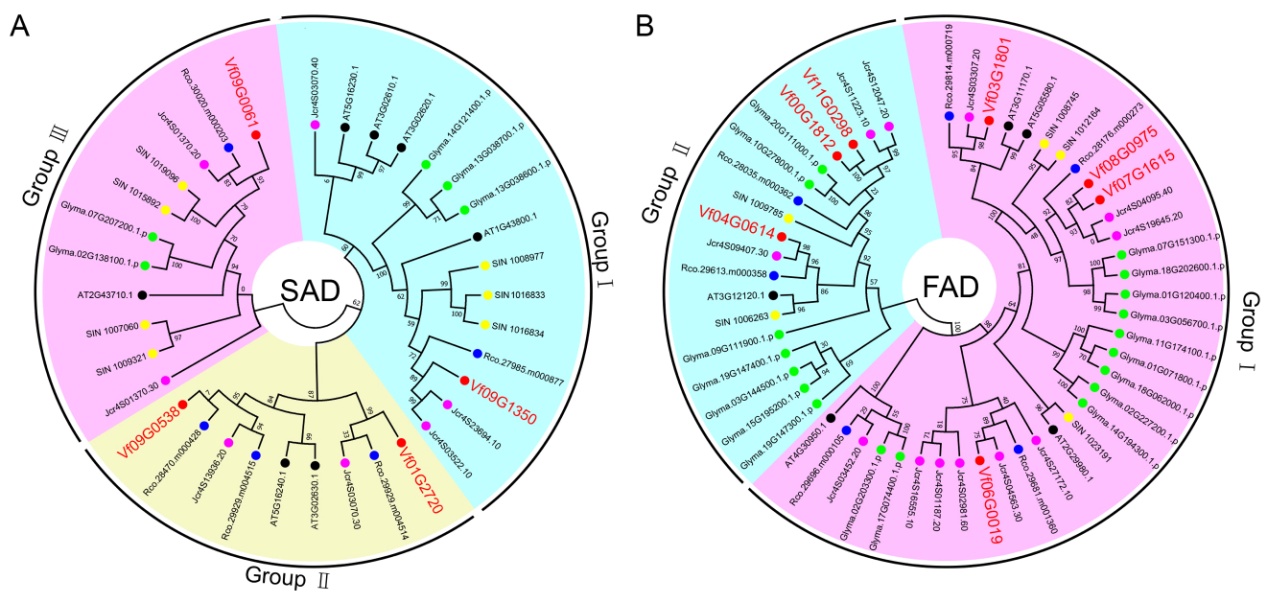

**Figure S27.** Phylogenetic analysis of fatty acid desaturases.

Maximum-likelihood phylogenetic trees of SAD (A) and FAD (B) constructed from protein sequences from *V. fordii* (red dots), *J. curcas* (pink dots), *S. indicum* (yellow dots), *R. communis* (blue dots), *G. max* (green dots) and *A. thaliana* (black dots). Different color represents different gene group generated from the tree.

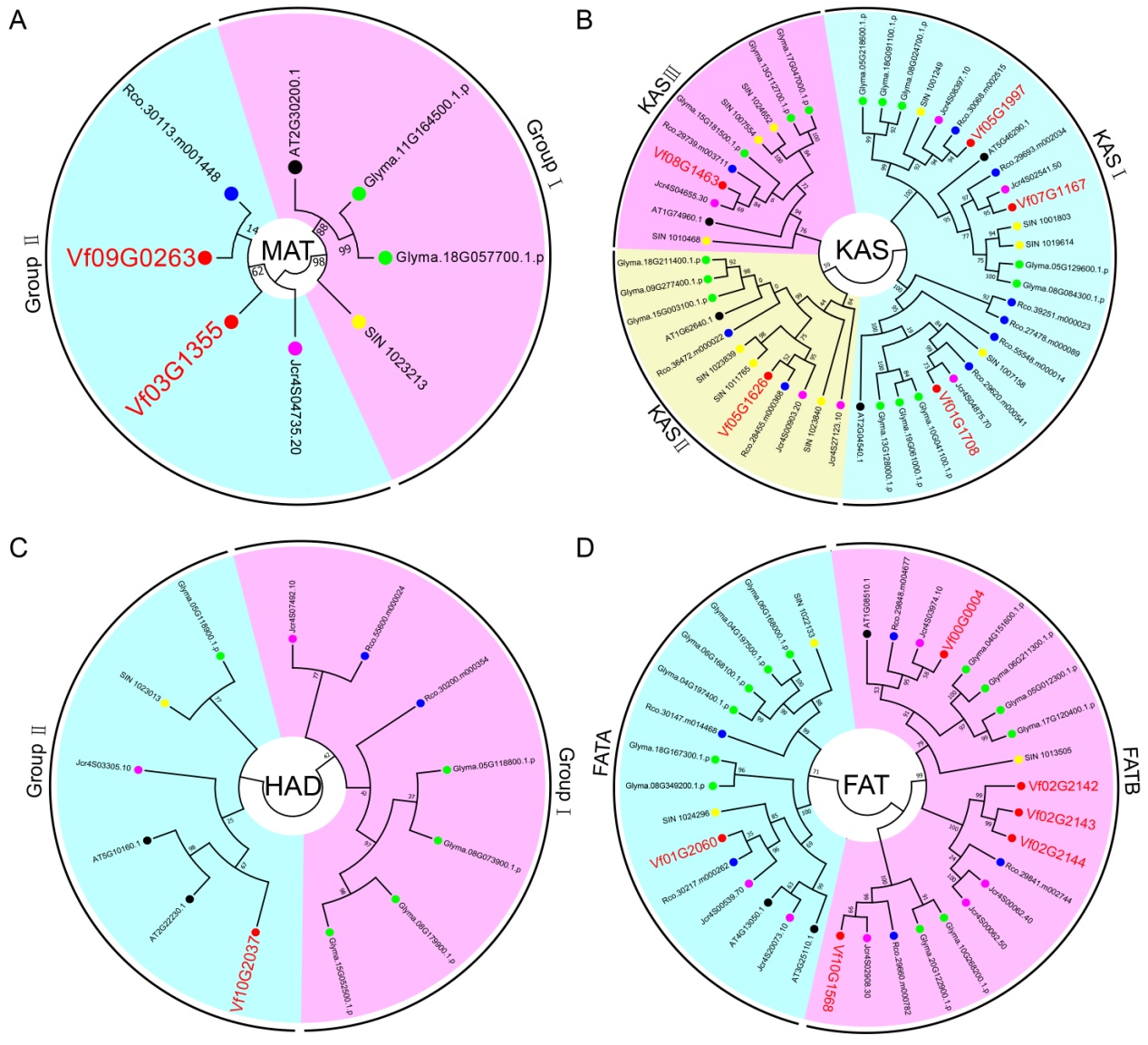

**Figure S28.** Phylogenetic analysis of fatty acid synthetases.

Maximum-likelihood phylogenetic trees of MAT (A), KAS (B), HAD (C) and FAT (D) constructed from protein sequences from *V. fordii* (red dots), *J. curcas* (pink dots), *S. indicum* (yellow dots), *R. communis* (blue dots), *G. max* (green dots) and *A. thaliana* (black dots). Different color represents different gene group generated from the tree.

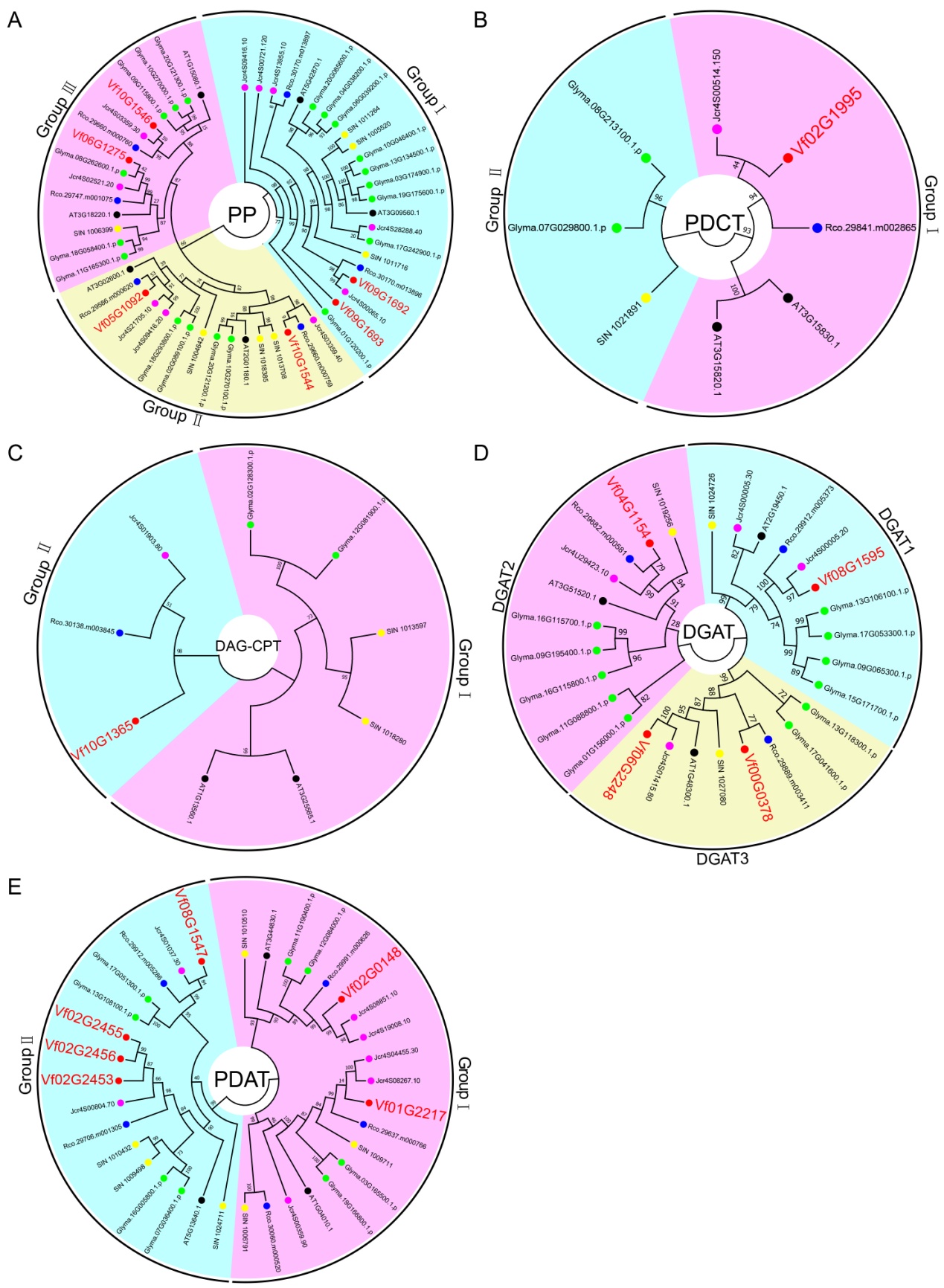

**Figure S29.** Phylogenetic analysis of TAG synthesis-related genes. Maximum-likelihood phylogenetic trees of PP (A), PDCT (B), DAG-CPT (C), DGAT (D) and PDAT (E) constructed from protein sequences from *V. fordii* (red dots), *J. curcas* (pink dots), *S. indicum* (yellow dots), *R. communis* (blue dots), *G. max* (green dots) and *A. thaliana* (black dots). Different color represents different gene group generated from the tree.

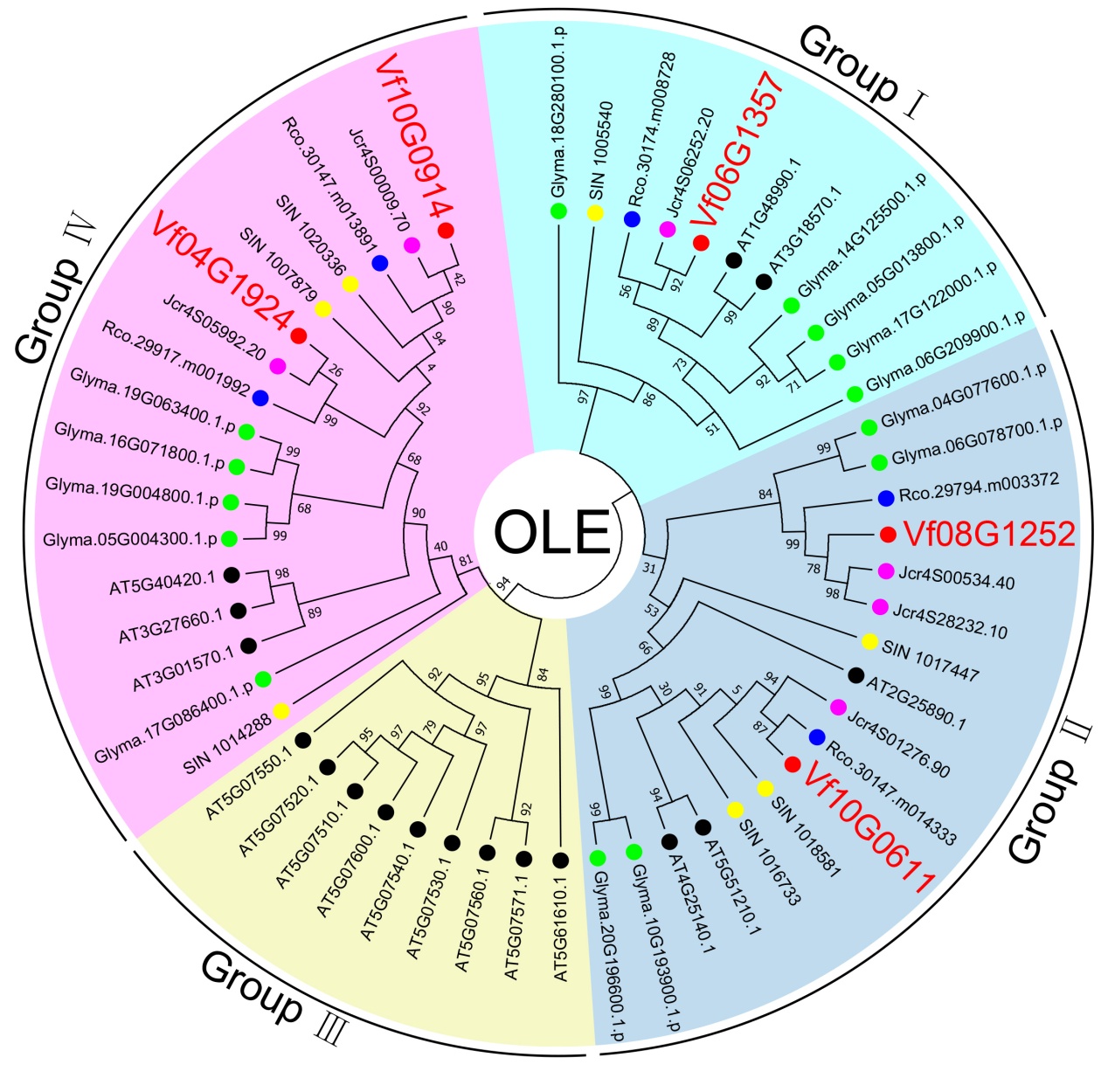

**Figure S30.** Phylogenetic analysis of oleosins. A Maximum-likelihood phylogenetic tree constructed from protein sequences from *V. fordii* (red dots), *J. curcas* (pink dots), *S. indicum* (yellow dots), *R. communis* (blue dots), *G. max* (green dots) and *A. thaliana* (black dots). Different color represents different gene group generated from the tree.
