## Supplementary material for "The Tung Tree (*Vernicia Fordii*) Genome Provides A Resource for Understanding Genome Evolution and Oil Improvement": File S12

**File S12: Gene co-expression analysis**

The primary transcripts with an average abundance (calculated from three biological replicates) of greater than 1 FPKM in at least one of the five seed samples were utilized to construct a weighted gene co-expression network using the R package WGCNA. Modules were constructed using the following parameters: maxBlockSize = 8000, power = 18, networkType = “unsigned”, mergeCutHeight = 0.25, minModuleSize = 30. Then, an effective threshold (0.8) of the Pearson’s correlation coefficient (PCC) value and p-value of 0.1 was trained to generate the lowest density networks. Finally, the oil synthesis genes and transcription factor (TF) (nodes) genes with their adjacent connections (edges) were extracted for our analysis. The constructed co-expression networks were visualized using the Cytoscape v.3.5.1 program (<http://www.cytoscape.org/>) [1].
