## Supplementary material for "The Tung Tree (*Vernicia Fordii*) Genome Provides A Resource for Understanding Genome Evolution and Oil Improvement": File S13

**File S13: Yeast two-hybrid assay of transcription factors**

FUS3, LEC1-1, LEC1-2, and ABI3 coding regions were cloned into the Y2H bait vector (BD) pDEST32 and the prey vector (AD) pDEST22 (Invitrogen, America), respectively, after amplifying with the primers shown in Table S59. Bait and pray constructs were co-transformed into *Saccharomyces cerevisiae* (SC) strain MaV203 (Invitrogen, America) and transformants were selected on glucose medium supplemented with Try^–^Leu^–^ drop-out solution (BD Biosciences) according to instructions of Invitrogen manual. The positive control plasmids BD-Krev1 and AD-RalGDS-wt were purchased from Invitrogen. To test the interactions between FUS3 and LEC1-1, FUS3 and LEC1-2, and FUS3 and ABI3 proteins, transformed yeast colonies were plated on SC-galactose/rafinose inducing medium containing Try^–^Leu^–^Ura^–^ drop out supplement and 80 μg/ml X-Gal, and incubated for 3-4 days at 30°C [1].

**Supplementary Table**

**Table S59. Primers used for yeast two-hybrid assay of tung tree transcription factors**

| Primer name | Primer sequence(5’——3’) | Annealing temperature (°C) |
| --- | --- | --- |
| ABI3-F | ATGAAGAGTTTGCACTTGC | 50.5 |
| ABI3-R | TTATACTGTTTGAGTTTGATTAGA | 49.7 |
| FUS3-F | ATGATGATGGAACAAGAGGC | 54.5 |
| FUS3-R | TTAATTGAACTCGTCGAGAGAC | 54.2 |
| LEC1-1-F | CTGTTCGTGATCCACTCC | 50.1 |
| LEC1-1-R | TCACTTGAACTGAGTAAATGG | 50.2 |
| LEC1-2-F | ATGGAACGTGCAGGAGGTTTCC | 65 |
| LEC1-2-R | TCATTTATGCTGGGCAAATGC | 61.2 |
