## Supplementary material for "The Tung Tree (*Vernicia Fordii*) Genome Provides A Resource for Understanding Genome Evolution and Oil Improvement": Table S2

**Table S2 Sequencing data for 500 kb-library used in genome survey**

| **Insert size (bp)** | **Raw reads (million)** | **Raw base (Gb)** | **Read length (bp)** | **Q20** | **GC (%)** |
| --- | --- | --- | --- | --- | --- |
| 500 | 365.14 | 36.51 | 100PE | 94.59 | 35.31 |
