## Supplementary material for "The Tung Tree (*Vernicia Fordii*) Genome Provides A Resource for Understanding Genome Evolution and Oil Improvement": Table S3

**Supplementary Table S3. Sequencing data for 17-mer analysis**

| Kmer size (bp) | Kmer number | Kmer depth (X) | Genome size (bp) | Used bases (bp) | Used reads | X |
| --- | --- | --- | --- | --- | --- | --- |
| 17 | 30,007,782,380 | 22.82 | 1,314,741,534 | 36,513,719,600 | 365,137,196 | 27.77 |
