## Supplementary material for "The Tung Tree (*Vernicia Fordii*) Genome Provides A Resource for Understanding Genome Evolution and Oil Improvement": Table S4

**Supplementary Table S4. Statistics of sequencing data for tung tree genome**

| Lib | Insert size (bp) | Raw data | | | | Clean data | | | | dupRae |
| --- | --- | --- | --- | --- | --- | --- | --- | --- | --- | --- |
|  |  | Reads length (bp) | Reads number (million) | Base number (Gb) | Depth (X) | Reads length (bp) | Reads number (million) | Base number (Gb) | Depth (X) |  |
| 180 | 159 | 100 | 465.19 | 46.52 | 35.38 | 95 | 443.35 | 42.12 | 32.04 | 3.91% |
| 500_1 | 460 | 100 | 365.14 | 36.51 | 27.77 | 95 | 323.35 | 30.72 | 23.36 | 2.67% |
| 500_2 | 474 | 100 | 61.59 | 6.16 | 4.68 | 95 | 52.23 | 4.96 | 3.77 | 0.73% |
| 800 | 734 | 100 | 276.98 | 27.70 | 21.07 | 95 | 231.37 | 21.98 | 16.72 | 16.84% |
| 3K_1 | 2,680 | 100 | 344.45 | 34.44 | 26.20 | 95 | 47.06 | 4.47 | 3.40 | 77.76% |
| 3K_2 | 3,360 | 100 | 389.91 | 38.99 | 29.99 | 95 | 285.36 | 27.11 | 20.85 | 25.28% |
| 10K_1 | 8,354 | 100 | 220.02 | 22.00 | 16.73 | 95 | 27.34 | 2.60 | 1.98 | 42.29% |
| 10K_2 | 8,828 | 100 | 360.96 | 36.10 | 27.77 | 95 | 159.57 | 15.16 | 11.66 | 62.09% |
| 15K_1 | 10,785 | 100 | 145.27 | 14.53 | 11.05 | 95 | 21.58 | 2.05 | 1.56 | 40.99% |
| 15K_2 | 14,052 | 100 | 458.61 | 45.86 | 35.28 | 95 | 95.19 | 9.04 | 6.96 | 70.15% |
| Pacbio_1 | 20,000 | 6,197.13 | 2.82 | 17.49 | 13.45 | 6,222 | 2.74 | 17.03 | 13.1 | - |
| Pacbio_2 | 20,000 | 8,136.20 | 0.09 | 0.76 | 0.58 | 8,349 | 0.05 | 0.44 | 0.33 | - |
| Total | - | - | 3,091.03 | 327.07 | 251.59 | - | 1,689.19 | 177.68 | 135.73 | - |
