## Supplementary material for "The Tung Tree (*Vernicia Fordii*) Genome Provides A Resource for Understanding Genome Evolution and Oil Improvement": Table S5

**Supplementary Table S5. Statistics of the tung tree genome assembly**

| Stat type | Scaffold length | Scaffold number | Contig length | Contig number |
| --- | --- | --- | --- | --- |
| N50 | 803,761 | 406 | 60,554 | 5,277 |
| N60 | 639,424 | 561 | 49,364 | 7,219 |
| N70 | 486,231 | 761 | 38,598 | 9,649 |
| N80 | 359,334 | 1,028 | 28,347 | 12,844 |
| N90 | 225,380 | 1,415 | 17,260 | 17,576 |
| Longest | 5,087,465 | 1 | 544,109 | 1 |
| Total | 1,118,693,778 | 4,577 | 1,060,078,069 | 34,773 |
| Length>100bp | 1,118,693,778 | 4,577 | 1,060,078,069 | 34,773 |
| Length>2kb | 1,116,988,597 | 3,333 | 1,052,727,832 | 29,721 |
