## Supplementary material for "The Tung Tree (*Vernicia Fordii*) Genome Provides A Resource for Understanding Genome Evolution and Oil Improvement": Table S7

**Supplementary Table S7. Mapped reads of Hi-C library to tung tree genome**

| Mapping type | Number | Ratio（%） |
| --- | --- | --- |
| Total read pairs | 206,982,355 | 100 |
| Mapped reads | 393,022,114 | 94.94 |
| Unique mapped read pairs | 95,209,829 | 46.00 |
