## Supplementary material for "The Tung Tree (*Vernicia Fordii*) Genome Provides A Resource for Understanding Genome Evolution and Oil Improvement": Table S8

**Supplementary Table S8. Statistics of Hi-C sequencing data**

| Type | Number | Ratio (%) |
| --- | --- | --- |
| Unique paired alignments | 95,209,829 | 100 |
| Valid interaction pairs | 85,519,051 | 89.82 |
| Dangling end pairs | 3,094,066 | 3.25 |
| Re-ligation pairs | 902,166 | 0.95 |
| Self-cycle pairs | 482,455 | 0.51 |
| Dumped pairs | 5,212,091 | 5.47 |
