## Supplementary material for "The Tung Tree (*Vernicia Fordii*) Genome Provides A Resource for Understanding Genome Evolution and Oil Improvement": Table S9

**Supplementary Table S9. Efficient coverage of Hi-C data to tung tree genome assembly**

| Cov Num | Num | Num ratio (%) | Len (bp) | Len ratio (%) |
| --- | --- | --- | --- | --- |
| 0 | 1,148 | 25.08 | 2,481,599 | 0.22 |
| >1 | 3,429 | 74.92 | 1,116,212,179 | 99.78 |
| >5 | 3,263 | 71.29 | 1,115,492,730 | 99.71 |
| >10 | 3,198 | 69.87 | 1,115,280,626 | 99.69 |
| >20 | 3,156 | 68.95 | 1,114,995,846 | 99.67 |

Note: Cov Num: number of efficiently coverage read pairs; Num: number of efficiently coverage sequences; Num Ratio: ratio of number of efficiently coverage sequences to number of whole genome sequences; Len: length of efficiently coverage sequences; Len Ratio: ration of length of efficiently coverage sequences to length of whole genome.
