## Supplementary material for "The Tung Tree (*Vernicia Fordii*) Genome Provides A Resource for Understanding Genome Evolution and Oil Improvement": Table S10

**Supplementary Table S10. Data of tung tree genome after Hi-C assembly**

| Group | Sequence number | Sequence length (bp) |
| --- | --- | --- |
| Chromosome1 | 400 | 120,568,480 |
| Chromosome2 | 385 | 114,111,371 |
| Chromosome3 | 398 | 114,133,434 |
| Chromosome4 | 390 | 112,337,999 |
| Chromosome5 | 344 | 95,844,693 |
| Chromosome6 | 319 | 95,162,274 |
| Chromosome7 | 272 | 90,491,488 |
| Chromosome8 | 292 | 86,622,686 |
| Chromosome9 | 286 | 88,869,008 |
| Chromosome10 | 176 | 63,428,274 |
| Chromosome11 | 305 | 81,518,211 |
| Total Sequences Clustered | 3,567 | 1,063,087,918 |
| Total Sequences Ordered and Oriented | 2,989 | 962,130,436 |
