## Supplementary material for "The Tung Tree (*Vernicia Fordii*) Genome Provides A Resource for Understanding Genome Evolution and Oil Improvement": Table S11

**Supplementary Table S11. Scaffold information after Hi-C assembly**

| Scaffold number | Scaffold length (bp) | Scaffold N50 (bp) | Scaffold N90 (bp) | Scaffold max (bp) | Gap total length (bp) |
| --- | --- | --- | --- | --- | --- |
| 3,314 | 1,117,565,834 | 87,145,426 | 418,876 | 112,054,921 | 57,502,275 |

Note: Scaffold number: scaffold number with length ≥1Kb; Scaffold length (bp): scaffold length with length ≥1Kb; Scaffold N50 (bp): sequence length of scaffold N50 with length ≥1Kb; Scaffold N90 (bp): sequence length of scaffold N90 with length ≥1Kb; Scaffold max (bp): sequence length of longest scaffold.
