## Supplementary material for "The Tung Tree (*Vernicia Fordii*) Genome Provides A Resource for Understanding Genome Evolution and Oil Improvement": Table S12

**Supplementary Table S12. Assessment for completeness of tung tree genome by CEGMA**

|  | Prots | % Completeness | Total | Average | % Ortho |
| --- | --- | --- | --- | --- | --- |
| Complete | 218 | 87.9 | 350 | 1.61 | 39.91 |
| Group 1 | 59 | 89.39 | 81 | 1.37 | 25.42 |
| Group 2 | 49 | 87.5 | 74 | 1.51 | 36.73 |
| Group 3 | 51 | 83.61 | 89 | 1.75 | 49.02 |
| Group 4 | 59 | 90.77 | 106 | 1.8 | 49.15 |
| Partial | 238 | 95.97 | 440 | 1.85 | 53.78 |
| Group 1 | 63 | 95.45 | 98 | 1.56 | 38.1 |
| Group 2 | 54 | 96.43 | 99 | 1.83 | 51.85 |
| Group 3 | 58 | 95.08 | 117 | 2.02 | 65.52 |
| Group 4 | 63 | 96.92 | 126 | 2 | 60.32 |
