## Supplementary material for "The Tung Tree (*Vernicia Fordii*) Genome Provides A Resource for Understanding Genome Evolution and Oil Improvement": Table S13

**Supplementary Table S13. Assessment for completeness of tung tree genome by BUSCO**

| **Type** | **Number** | **Percent (%)** |
| --- | --- | --- |
| **Complete BUSCOs (C)** | 1379 | 95.7 |
| **Complete and single-copy BUSCOs (S)** | 1338 | 92.9 |
| **Complete and duplicated BUSCOs (D)** | 41 | 2.8 |
| **Fragmented BUSCOs (F)** | 16 | 1.1 |
| **Missing BUSCOs (M)** | 45 | 3.2 |
| **Total BUSCO groups searched** | 1,440 | - |
