## Supplementary material for "The Tung Tree (*Vernicia Fordii*) Genome Provides A Resource for Understanding Genome Evolution and Oil Improvement": Table S14

**Supplementary Table S14. Coverage of tung tree genome from male flower unigenes**

| Dataset | Number | Total length (bp) | Covered by assembly (%) | Covered >90% in one sequence | | Covered >50% in one sequence | |
| --- | --- | --- | --- | --- | --- | --- | --- |
|  |  |  |  | Number | Percent (%) | Number | Percent (%) |
| All | 65,481 | 44,918,574 | 94.97 | 59,173 | 90.36 | 62,058 | 94.77 |
| >200bp | 65,214 | 44,865,174 | 94.99 | 58,949 | 90.39 | 61,820 | 94.79 |
| >500bp | 30,053 | 33,955,900 | 98.03 | 28,216 | 93.88 | 29,392 | 97.80 |
| >1000bp | 13,570 | 22,216,702 | 99.12 | 12,792 | 94.26 | 13,405 | 98.78 |
