## Supplementary material for "The Tung Tree (*Vernicia Fordii*) Genome Provides A Resource for Understanding Genome Evolution and Oil Improvement": Table S16

**Supplementary Table S15. Coverage of tung tree genome from female flower unigenes**

| Dataset | Number | Total length (bp) | Covered by assembly (%) | Covered >90% in one sequence | | Covered >50% in one sequence | |
| --- | --- | --- | --- | --- | --- | --- | --- |
|  |  |  |  | Number | Percent (%) | Number | Percent (%) |
| All | 59,270 | 44,714,006 | 99.59 | 57,395 | 96.83 | 58,933 | 99.43 |
| >200 bp | 59,088 | 44,677,606 | 99.59 | 57,217 | 96.83 | 58,754 | 99.43 |
| >500 bp | 28,509 | 35,308,668 | 99.81 | 27,374 | 96.01 | 28,407 | 99.64 |
| >1000 bp | 14,829 | 25,481,547 | 99.86 | 14,047 | 94.72 | 14,779 | 99.66 |
