## Supplementary material for "The Tung Tree (*Vernicia Fordii*) Genome Provides A Resource for Understanding Genome Evolution and Oil Improvement": Table S17

**Supplementary Table S17. Coverage bof tung tree genome from seed2 unigenes**

| Dataset | Number | Total length (bp) | Covered by assembly (%) | Covered >90% in one sequence | | Covered >50% in one sequence | |
| --- | --- | --- | --- | --- | --- | --- | --- |
|  |  |  |  | Number | Percent (%) | Number | Percent (%) |
| All | 54,679 | 41,808,916 | 99.62 | 52,222 | 95.50 | 54,358 | 99.41 |
| >200bp | 46,663 | 40,425,178 | 99.69 | 44,491 | 95.34 | 46,411 | 99.45 |
| >500bp | 24,843 | 33,634,583 | 99.87 | 23,350 | 93.99 | 24,748 | 99.61 |
| >1000bp | 14,290 | 26,030,052 | 99.92 | 13,228 | 92.56 | 14,229 | 99.57 |
