## Supplementary material for "The Tung Tree (*Vernicia Fordii*) Genome Provides A Resource for Understanding Genome Evolution and Oil Improvement": Table S18

**Supplementary Table S18. Coverage of tung tree genome from seed 3 unigenes**

| Dataset | Number | Total length (bp) | Covered by assembly (%) | Covered >90% in one sequence | | Covered >50% in one sequence | |
| --- | --- | --- | --- | --- | --- | --- | --- |
|  |  |  |  | Number | Percent (%) | Number | Percent (%) |
| All | 44,495 | 28,580,185 | 99.52 | 42,930 | 96.48 | 44,189 | 99.31 |
| >200bp | 38,184 | 27,483,212 | 99.61 | 36,845 | 96.49 | 37,951 | 99.38 |
| >500bp | 17,803 | 21,194,898 | 99.83 | 16,936 | 95.13 | 17,714 | 99.50 |
| >1000bp | 8,863 | 14,820,054 | 99.81 | 8,306 | 93.71 | 8,810 | 99.40 |
