## Supplementary material for "The Tung Tree (*Vernicia Fordii*) Genome Provides A Resource for Understanding Genome Evolution and Oil Improvement": Table S19

**Supplementary Table S19. Coverage of tung tree genome assembly from merged seed unigenes**

| Dataset | Number | Total length (bp) | Covered by assembly (%) | Covered >90% in one sequence | | Covered >50% in one sequence | |
| --- | --- | --- | --- | --- | --- | --- | --- |
|  |  |  |  | Number | Percent (%) | Number | Percent (%) |
| All | 58,439 | 51,962,216 | 99.61 | 54,728 | 93.64 | 58,060 | 99.35 |
| >200bp | 58,246 | 51,923,616 | 99.61 | 54,539 | 93.63 | 57,867 | 99.34 |
| >500bp | 31,146 | 43,542,191 | 99.85 | 28,552 | 91.67 | 30,986 | 99.48 |
| >1000bp | 18,321 | 34,303,644 | 99.89 | 16,462 | 89.85 | 18,211 | 99.39 |
