## Supplementary material for "The Tung Tree (*Vernicia Fordii*) Genome Provides A Resource for Understanding Genome Evolution and Oil Improvement": Table S20

**Supplementary Table S20. Transcriptomic reads mapped to tung tree genome**

| Sample | All reads number | Mapped reads number | Mapping rate  (%) |
| --- | --- | --- | --- |
| Male flower | 51,746,368 | 45,686,156 | 88.3 |
| Female flower | 54,398,380 | 51,994,799 | 95.6 |
| seed1 | 52,734,436 | 49,093,204 | 93.1 |
| seed2 | 52,039,198 | 48,517,867 | 93.2 |
| seed3 | 51,886,702 | 48,538,495 | 93.5 |
