## Supplementary material for "The Tung Tree (*Vernicia Fordii*) Genome Provides A Resource for Understanding Genome Evolution and Oil Improvement": Table S21

**Supplementary Table S21. Comparison of gene modules between tung tree and other species**

| Species | Total number of gene | Average trainscript length (bp) | Average CDS length (bp) | Average exons number per gene | Average exon length (b) | Average intron length (bp) |
| --- | --- | --- | --- | --- | --- | --- |
| *V.fordii* | 28,422 | 3,785.26 | 1,033.92 | 4.85 | 213.11 | 714.36 |
| *A.thaliana* | 27,173 | 1,876.19 | 1,221.69 | 5.15 | 237 | 157.53 |
| *R.communis* | 29,957 | 2,246.22 | 1,019.48 | 4.26 | 239.49 | 376.66 |
| *O.sativa* | 39,044 | 2,330.36 | 1,064.12 | 4.12 | 258.54 | 406.38 |
| *V.vinifera* | 26,346 | 5,936.60 | 1,137.11 | 5.95 | 191.1 | 969.55 |
| *J.curcas* | 22,139 | 3,478.87 | 1,321.13 | 5.42 | 243.83 | 488.37 |
