## Supplementary material for "The Tung Tree (*Vernicia Fordii*) Genome Provides A Resource for Understanding Genome Evolution and Oil Improvement": Table S22

**Supplementary Table S22. The GC content across the tung tree genome**

| Base | Number (bp) | % of genome |
| --- | --- | --- |
| A | 351420780 | 31.41 |
| T | 351480608 | 31.42 |
| G | 178680351 | 15.97 |
| C | 178496330 | 15.96 |
| N | 58615709 | 5.24 |
| GC | 357176681 | 31.93 |
| Total bases | 1118693778 |  |
