## Supplementary material for "The Tung Tree (*Vernicia Fordii*) Genome Provides A Resource for Understanding Genome Evolution and Oil Improvement": Table S23

**Supplementary Table S23. The GC content in coding sequences in the tung tree genome**

| Base | Number (bp) | % of genome |
| --- | --- | --- |
| A | 8495424 | 29.05 |
| T | 8493364 | 29.04 |
| G | 6146790 | 21.02 |
| C | 6112416 | 20.90 |
| N | 62 | 0.00 |
| GC | 12259206 | 41.91 |
| Total bases | 29248056 |  |
