## Supplementary material for "The Tung Tree (*Vernicia Fordii*) Genome Provides A Resource for Understanding Genome Evolution and Oil Improvement": Table S24

**Supplementary Table S24. The GC content in intron regions in the tung tree genome**

| Base | Number (bp) | % of genome |
| --- | --- | --- |
| A | 26394030 | 33.90 |
| T | 26406844 | 33.92 |
| G | 12126220 | 15.58 |
| C | 12131617 | 15.58 |
| N | 797965 | 1.02 |
| GC | 24257837 | 31.16 |
| Total bases | 77856676 |  |
