## Supplementary material for "The Tung Tree (*Vernicia Fordii*) Genome Provides A Resource for Understanding Genome Evolution and Oil Improvement": Table S25

**Supplementary Table S25. Assessment for completeness of predicted tung tree genes by BUSCO**

| **Type** | **Number** | **Percent (%)** |
| --- | --- | --- |
| **Complete BUSCOs (C)** | 1290 | 89.6 |
| **Complete and single-copy BUSCOs (S)** | 1251 | 86.9 |
| **Complete and duplicated BUSCOs (D)** | 39 | 2.7 |
| **Fragmented BUSCOs (F)** | 38 | 2.6 |
| **Missing BUSCOs (M)** | 112 | 7.8 |
| **Total BUSCO groups searched** | 1,440 | - |
