## Supplementary material for "The Tung Tree (*Vernicia Fordii*) Genome Provides A Resource for Understanding Genome Evolution and Oil Improvement": Table S28

**Supplementary Table S28. Non-coding RNAs in tung tree genome**

| Type | | Copy number | Average length (bp) | Total length (bp) | Percentage of genome (%) |
| --- | --- | --- | --- | --- | --- |
| rRNA | rRNA | 116 | 983 | 105,204 | 0.009404 |
|  | 18S | 8 | 2,384 | 19,069 | 0.001705 |
|  | 28S | 13 | 5,865 | 76,247 | 0.006816 |
|  | 5.8S | 9 | 155 | 1,392 | 0.000124 |
|  | 5S | 86 | 115 | 9,888 | 0.000884 |
| snRNA | snRNA | 1,414 | 107 | 151,107 | 0.013507 |
|  | CD-box | 1,289 | 104 | 134,149 | 0.011992 |
|  | HACA-box | 39 | 125 | 4,858 | 0.000434 |
|  | splicing | 86 | 141 | 12,100 | 0.001082 |
| miRNA | | 465 | 148 | 68,972 | 0.006165 |
