## Supplementary material for "The Tung Tree (*Vernicia Fordii*) Genome Provides A Resource for Understanding Genome Evolution and Oil Improvement": Table S29

**Supplementary Table S29. Statistics of gene families of *V. forrdi* and other 7 species**

| Species | Genes number | Genes number in  families | Unclustered genes number | Family number | Unique  families number | Average genes  number per family |
| --- | --- | --- | --- | --- | --- | --- |
| *A.thaliana* | 27,173 | 22,858 | 4,315 | 12,892 | 744 | 1.77 |
| *H.brasiliensis* | 34,661 | 30,787 | 3,874 | 15,410 | 299 | 2 |
| *J.carcas* | 22,139 | 20,785 | 1,354 | 14,969 | 166 | 1.39 |
| *M.esculenta* | 33,033 | 26,921 | 6,112 | 15,641 | 184 | 1.72 |
| *P.trichocarpa* | 41,335 | 32,970 | 8,365 | 15,376 | 803 | 2.14 |
| *R.communis* | 29,957 | 20,499 | 9,458 | 15,294 | 565 | 1.34 |
| *V.fordii* | 28,422 | 22,991 | 5,431 | 15,038 | 635 | 1.53 |
| *V.vinifera* | 26,346 | 19,025 | 7,321 | 12,865 | 623 | 1.48 |

Unclustered genes: species-specific genes that were not assigned to any families.

Unique family: species-specific paralogous gene families.
