## Supplementary material for "The Tung Tree (*Vernicia Fordii*) Genome Provides A Resource for Understanding Genome Evolution and Oil Improvement": Table S40

**Supplementary Table S40. Summary of repeat sequences in tung tree genome**

| Type | Repeat size(bp) | % of genome |
| --- | --- | --- |
| Trf | 49,820,132 | 4.70 |
| RepeatProteinMask | 222,266,652 | 20.97 |
| RepeatMasker (Repbase) | 194,834,599 | 18.38 |
| RepeatMasker (Mips-REdat) | 209,512,020 | 19.76 |
| Denovo (RepeatModeler) | 717,652,523 | 67.70 |
| Total | 777,464,634 | 73.34 |

Note: Total indicates non-redundant repeat size generated by the above five methods.
