## Supplementary material for "The Tung Tree (*Vernicia Fordii*) Genome Provides A Resource for Understanding Genome Evolution and Oil Improvement": Table S41

**Supplementary Table S41. Annotation of repeat sequences in tung tree genome**

|  |  |  | count | total_repeat | repeat_percent (%) | genome_percent (%) |
| --- | --- | --- | --- | --- | --- | --- |
| total repeat fraction |  |  | 1105885 | 777464634 | 100 | 73.34 |
| class i: retroelement |  |  | 403333 | 550023721 | 70.74581877 | 51.88520894 |
|  | LTR retrotransposon |  | 382737 | 538155445 | 69.21928297 | 50.76564271 |
|  |  | Ty1/Copia | 84180 | 117652898 | 15.13289388 | 11.09851259 |
|  |  | Ty3/Gypsy | 284597 | 415630052 | 53.45967313 | 39.20749463 |
|  |  | Other | 13960 | 4872495 | 0.626715967 | 0.459635487 |
|  | Non-LTR retrotransposon |  | 20585 | 11867162 | 1.526392518 | 1.119461137 |
|  |  | LINE | 19094 | 11683118 | 1.502720187 | 1.102099774 |
|  |  | SINE | 1491 | 184044 | 0.023672331 | 0.017361363 |
|  | Retroposon |  | 11 | 1114 | 0.000143286 | 0.000105087 |
| class ii: DNA transposon |  |  | 140493 | 59048669 | 7.595029589 | 5.570218904 |
|  |  | CMC | 22828 | 6803128 | 0.875040189 | 0.641757263 |
|  |  | hAT | 5560 | 1179911 | 0.151763945 | 0.111304161 |
|  |  | PIF/Harbinger | 653 | 364430 | 0.046874158 | 0.034377657 |
|  |  | Other | 111452 | 50701200 | 6.521351298 | 4.782779824 |
| Simple_repeat | non | non | 222091 | 80716851 | 10.3820608 | 7.614236476 |
| Unknown | non | non | 351473 | 112227111 | 14.43501172 | 10.58668359 |
| Satellite | non | non | 690 | 112112 | 0.014420206 | 0.010575825 |
