## Supplementary material for "The Tung Tree (*Vernicia Fordii*) Genome Provides A Resource for Understanding Genome Evolution and Oil Improvement": Table S51

**Supplementary Table S51．Four types of functional conservation and diversification of tung tree homologs**

| Type | Gene ID | Gene annotation |
| --- | --- | --- |
|  | Vf03G0652 | Feruloyl CoA ortho-hydroxylase 2 GN=F6&apos;H2 OS=Arabidopsis thaliana (Mouse-ear cress) PE=1 SV=1 |
| Conservation of function | Vf00G0634 | Feruloyl CoA ortho-hydroxylase 1 GN=F6&apos;H1 OS=Arabidopsis thaliana (Mouse-ear cress) PE=1 SV=1 |
|  | Vf03G0623 | Feruloyl CoA ortho-hydroxylase 1 GN=F6&apos;H1 OS=Arabidopsis thaliana (Mouse-ear cress) PE=1 SV=1 |
|  | Vf04G0546 | Protein ECERIFERUM 1 GN=CER1 OS=Arabidopsis thaliana (Mouse-ear cress) PE=1 SV=1 |
| Sub-functionalization | Vf06G2858 | Protein ECERIFERUM 1 GN=CER1 OS=Arabidopsis thaliana (Mouse-ear cress) PE=1 SV=1 |
|  | Vf06G2857 | Protein ECERIFERUM 1 GN=CER1 OS=Arabidopsis thaliana (Mouse-ear cress) PE=1 SV=1 |
|  | Vf04G0305 | Purple acid phosphatase 18 (Precursor) GN=K10D20.4 OS=Arabidopsis thaliana (Mouse-ear cress) PE=2 SV=1 |
| Sub/neo-functionalization | Vf04G0306 | Probable purple acid phosphatase 20 (Precursor) GN=F3C22.180 OS=Arabidopsis thaliana (Mouse-ear cress) PE=2 SV=1 |
|  | Vf11G0977 | Purple acid phosphatase 22 (Precursor) GN=F3C22.220 OS=Arabidopsis thaliana (Mouse-ear cress) PE=2 SV=1 |
|  | Vf09G1183 | Protein LYK5 (Precursor) GN=LYK5 OS=Arabidopsis thaliana (Mouse-ear cress) PE=2 SV=1 |
| No-functionalization | Vf03G0089 | Protein LYK5 (Precursor) GN=LYK5 OS=Arabidopsis thaliana (Mouse-ear cress) PE=2 SV=1 |
|  | Vf09G0959 | Protein LYK5 (Precursor) GN=LYK5 OS=Arabidopsis thaliana (Mouse-ear cress) PE=2 SV=1 |
