## Supplementary material for "The Tung Tree (*Vernicia Fordii*) Genome Provides A Resource for Understanding Genome Evolution and Oil Improvement": Table S52

**Supplementary Table S52. Cross-species comparison of NBS-encoding gene family number**

| Type | *V.fordii* | *S.indicum* | *R.communis* | *M.esculenta* | *J.curcas* | *H.brasiliensis* | *V. vinifera* | *A. thaliana* | *P. trichocarpa* | *Z. mays* | *O. sativa* |
| --- | --- | --- | --- | --- | --- | --- | --- | --- | --- | --- | --- |
| TIR-NBS | 0 | 0 | 7 | 3 | 15 | 4 | 3 | 17 | 13 | 0 | 0 |
| TIR-NBS-LRR | 0 | 0 | 25 | 27 | 40 | 15 | 17 | 79 | 78 | 0 | 0 |
| CC-NBS | 23 | 25 | 17 | 15 | 6 | 35 | 18 | 8 | 19 | 11 | 53 |
| CC-NBS-LRR | 7 | 5 | 21 | 40 | 19 | 45 | 28 | 17 | 119 | 58 | 402 |
| NBS-LRR | 16 | 23 | 67 | 124 | 91 | 186 | 121 | 20 | 120 | 31 | 74 |
| NBS | 42 | 118 | 95 | 103 | 104 | 198 | 129 | 26 | 53 | 7 | 16 |
| Total | 88 | 171 | 232 | 312 | 275 | 483 | 316 | 167 | 402 | 107 | 543 |
