## Supplementary material for "The Tung Tree (*Vernicia Fordii*) Genome Provides A Resource for Understanding Genome Evolution and Oil Improvement": Table S54

**Supplementary Table S54. Cross-species comparison of oil-related gene family number**

| Gene | *V.fordii* | *J.curcas* | *R.communis* | *A.thaliana* | *S.indicum* | *G.max* |
| --- | --- | --- | --- | --- | --- | --- |
| ACCase | 9 | 6 | 6 | 7 | 6 | 10 |
| MAT | 2 | 1 | 1 | 1 | 1 | 2 |
| KAS | 5 | 6 | 9 | 4 | 10 | 14 |
| KAR | 3 | 3 | 3 | 6 | 7 | 7 |
| HAD | 1 | 2 | 2 | 2 | 1 | 5 |
| EAR | 1 | 2 | 2 | 1 | 4 | 5 |
| FATA | 1 | 2 | 2 | 2 | 2 | 6 |
| FATB | 5 | 4 | 3 | 1 | 1 | 6 |
| SAD | 4 | 7 | 5 | 7 | 7 | 5 |
| FAD2 | 1 | 1 | 1 | 1 | 1 | 5 |
| FAD3/7 | 4 | 9 | 4 | 4 | 3 | 11 |
| FADX/FAH12 | 2 | 2 | 1 | 0 | 1 | 2 |
| LACS | 8 | 2 | 2 | 2 | 9 | 22 |
| DAG-CPT | 1 | 1 | 1 | 2 | 2 | 2 |
| PDCT | 1 | 1 | 1 | 2 | 1 | 2 |
| PP | 6 | 8 | 6 | 6 | 7 | 19 |
| LPAT | 7 | 5 | 6 | 5 | 7 | 11 |
| GPAT | 9 | 8 | 9 | 7 | 15 | 27 |
| PDAT | 6 | 7 | 5 | 3 | 6 | 8 |
| DGAT | 4 | 4 | 3 | 3 | 3 | 11 |
| OLE | 5 | 6 | 5 | 17 | 7 | 14 |
| PEPC | 3 | 4 | 7 | 4 | 4 | 16 |
| Total | 88 | 91 | 84 | 87 | 105 | 210 |
