## Supplementary material for "The Tung Tree (*Vernicia Fordii*) Genome Provides A Resource for Understanding Genome Evolution and Oil Improvement": Table S56

**Supplementary Table S56. Colinear gene pairs in poplar (*P. trichocarpa*)**

| BLOCK_NO | BLOCK_SCORE | E_VALUE | LOCUS_1 | LOCUS_2 | Ka | Ks |
| --- | --- | --- | --- | --- | --- | --- |
| 37 | 486 | 8.00E-15 | Potri.001G013700 | Potri.006G191700 | 0.7001 | 3.2078 |
| 37 | 486 | 2.00E-128 | Potri.001G013800 | Potri.006G191400 | 0.3004 | 1.5427 |
| 37 | 486 | 2.00E-51 | Potri.001G013200 | Potri.006G191900 | 0.312 | 1.0684 |
| 37 | 486 | 2.00E-177 | Potri.001G012500 (FADx) | Potri.006G192000 (FAD2) | 0.1704 | 2.2608 |
| 37 | 486 | 1.00E-179 | Potri.001G010900 | Potri.006G193900 | 0.2506 | 1.5559 |
| 37 | 486 | 2.00E-161 | Potri.001G010500 | Potri.006G194100 | 0.2015 | 2.7668 |
| 37 | 486 | 4.00E-69 | Potri.001G009100 | Potri.006G195300 | 0.3406 | 1.7346 |
| 37 | 486 | 6.00E-45 | Potri.001G008500 | Potri.006G195700 | 0.6179 | 3.8136 |
| 37 | 486 | 0 | Potri.001G007900 | Potri.006G196000 | 0.1915 | 1.2513 |
| 37 | 486 | 0 | Potri.001G007800 | Potri.006G196100 | 0.0579 | 1.2053 |
| 37 | 486 | 1.00E-160 | Potri.001G007000 | Potri.006G196900 | 0.2857 | 1.6713 |
| 37 | 486 | 1.00E-52 | Potri.001G006400 | Potri.006G197000 | 0.3281 | 2.0095 |
| 104 | 286 | 2.00E-132 | Potri.001G018900 | Potri.016G052700 | 0.2472 | 1.018 |
| 104 | 286 | 6.00E-180 | Potri.001G018700 | Potri.016G051900 | 0.3734 | 1.3881 |
| 104 | 286 | 2.00E-79 | Potri.001G018600 | Potri.016G051700 | 0.1742 | 1.2215 |
| 104 | 286 | 0 | Potri.001G015700 | Potri.016G049100 | 0.0993 | 1.6092 |
| 104 | 286 | 9.00E-137 | Potri.001G013800 | Potri.016G047600 | 0.2824 | 1.5449 |
| 104 | 286 | 0 | Potri.001G013500 | Potri.016G046700 | 0.1247 | 1.3067 |
| 104 | 286 | 2.00E-119 | Potri.001G013200 | Potri.016G046600 | 0.2814 | 1.1618 |
| 104 | 286 | 0 | Potri.001G012500 (FADx) | Potri.016G046200 (FAD2) | 0.1753 | 3.1035 |
