## Supplementary material for "The Tung Tree (*Vernicia Fordii*) Genome Provides A Resource for Understanding Genome Evolution and Oil Improvement": Figure S4

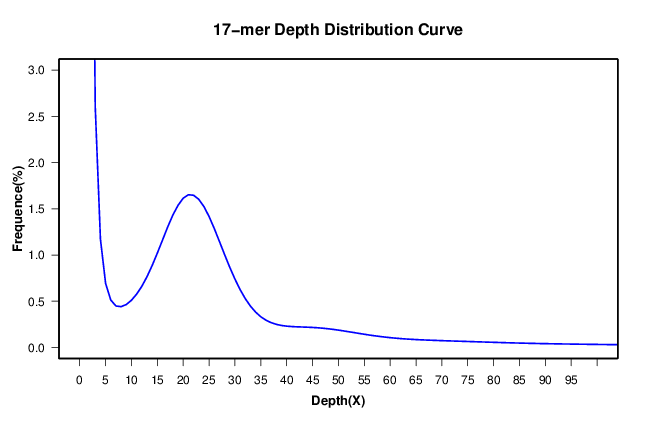

**Supplementary Figure S4. The k-mer analysis to estimate the tung tree genome size.** The figure shows frequency of 17 k-mers which are 17 bp sequences from the reads (after filtering) of short-insert size libraries. We identified 30,007,782,380 k-mers using 36.51 Gb data. The tung tree genome size was estimated to be 1.31 Gb by (total k-mer number) / (the volume peak).
