## Supplementary figures and images for "The Tung Tree (*Vernicia Fordii*) Genome Provides A Resource for Understanding Genome Evolution and Oil Improvement"

### Figure S5

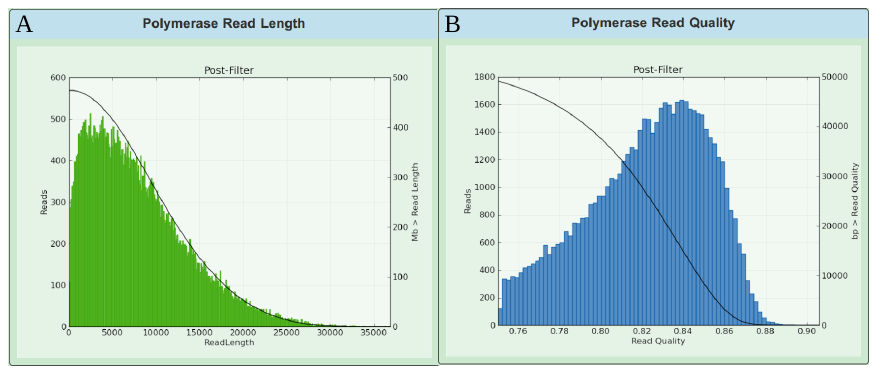

**Supplementary Figure S5. Distribution of length (A) and quality (B) of Pacbio raw reads.**

### Figure S6

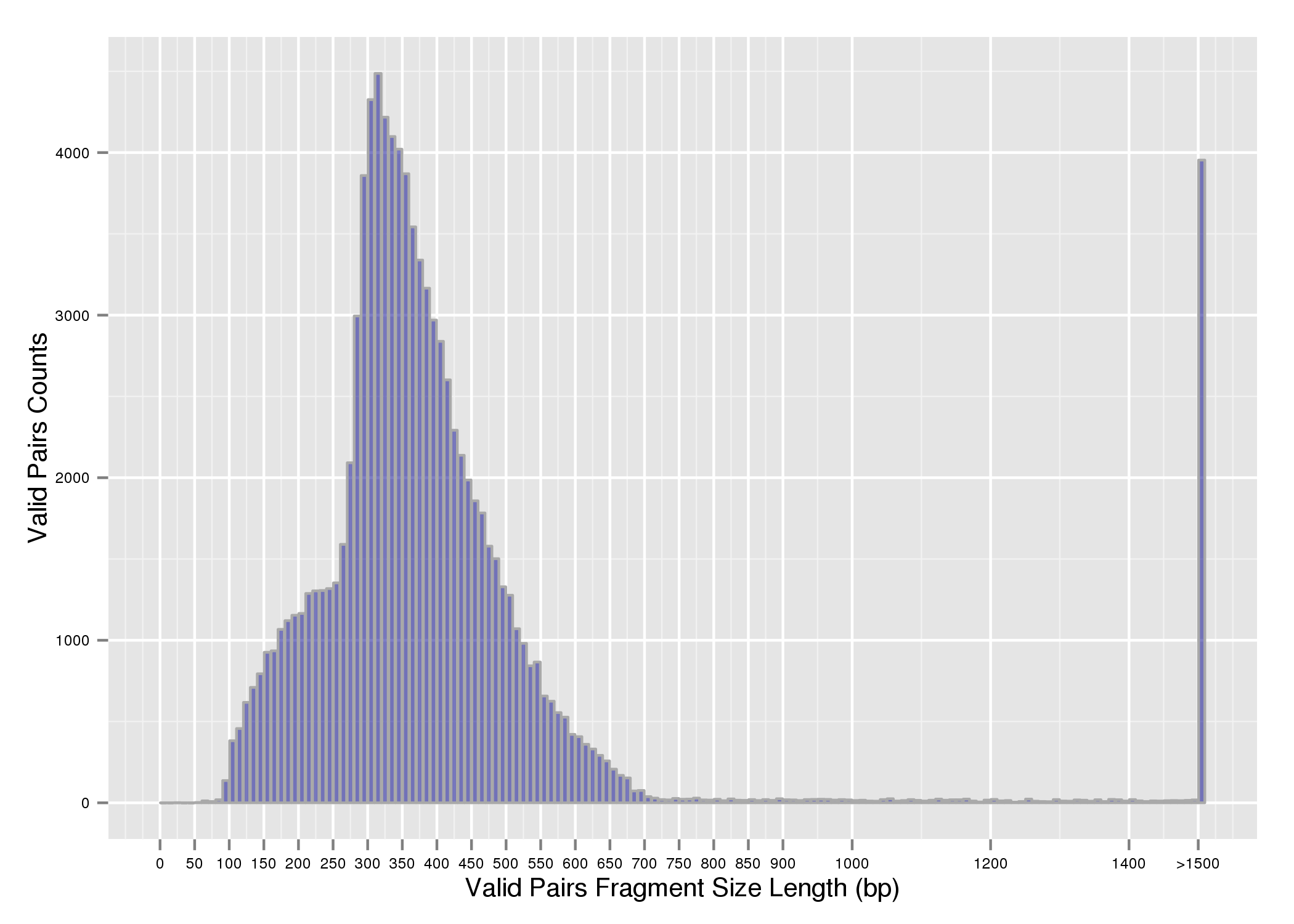

**Supplementary Figure S6. Distribution of inserted fragment length for Hi-C library.**

### Figure S10

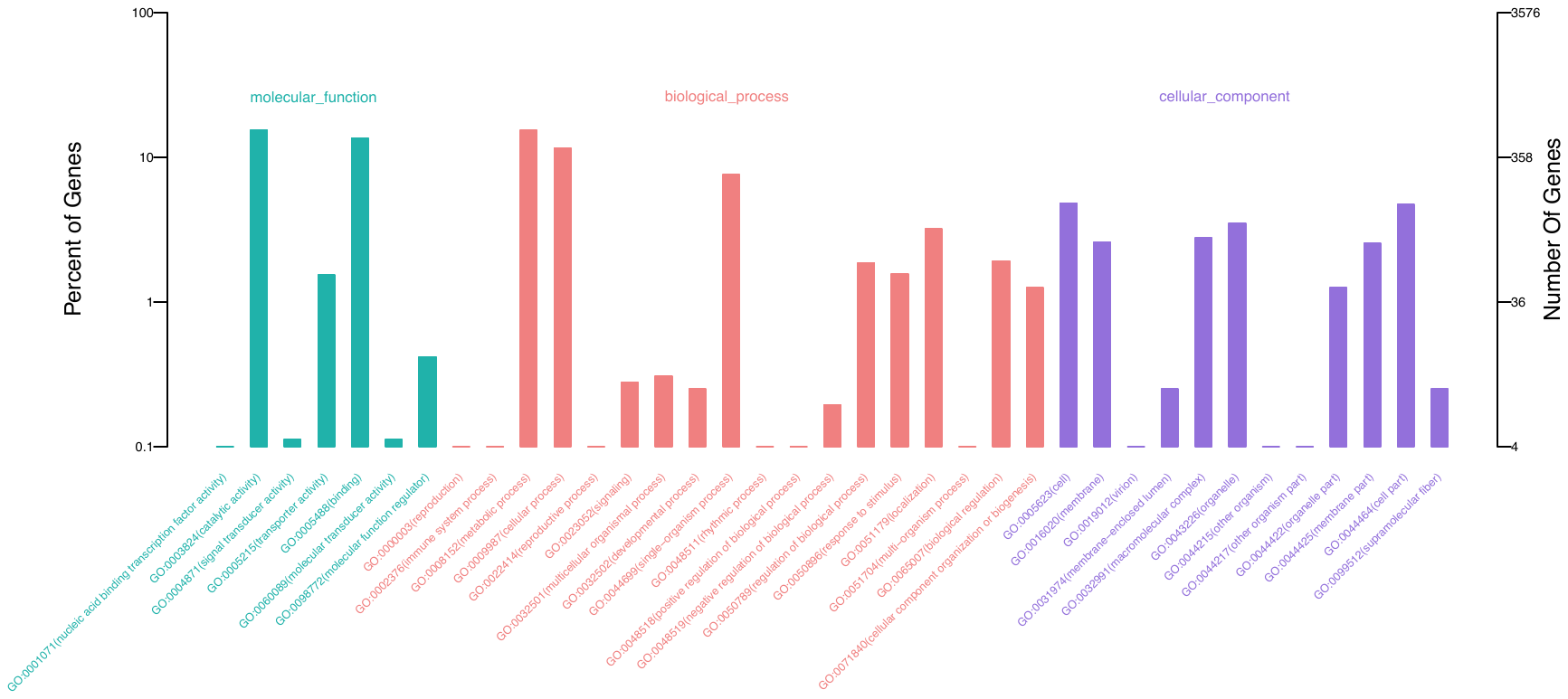

**Supplementary Figure S10. GO classification of tung tree-specific gene families.**

### Figure S12

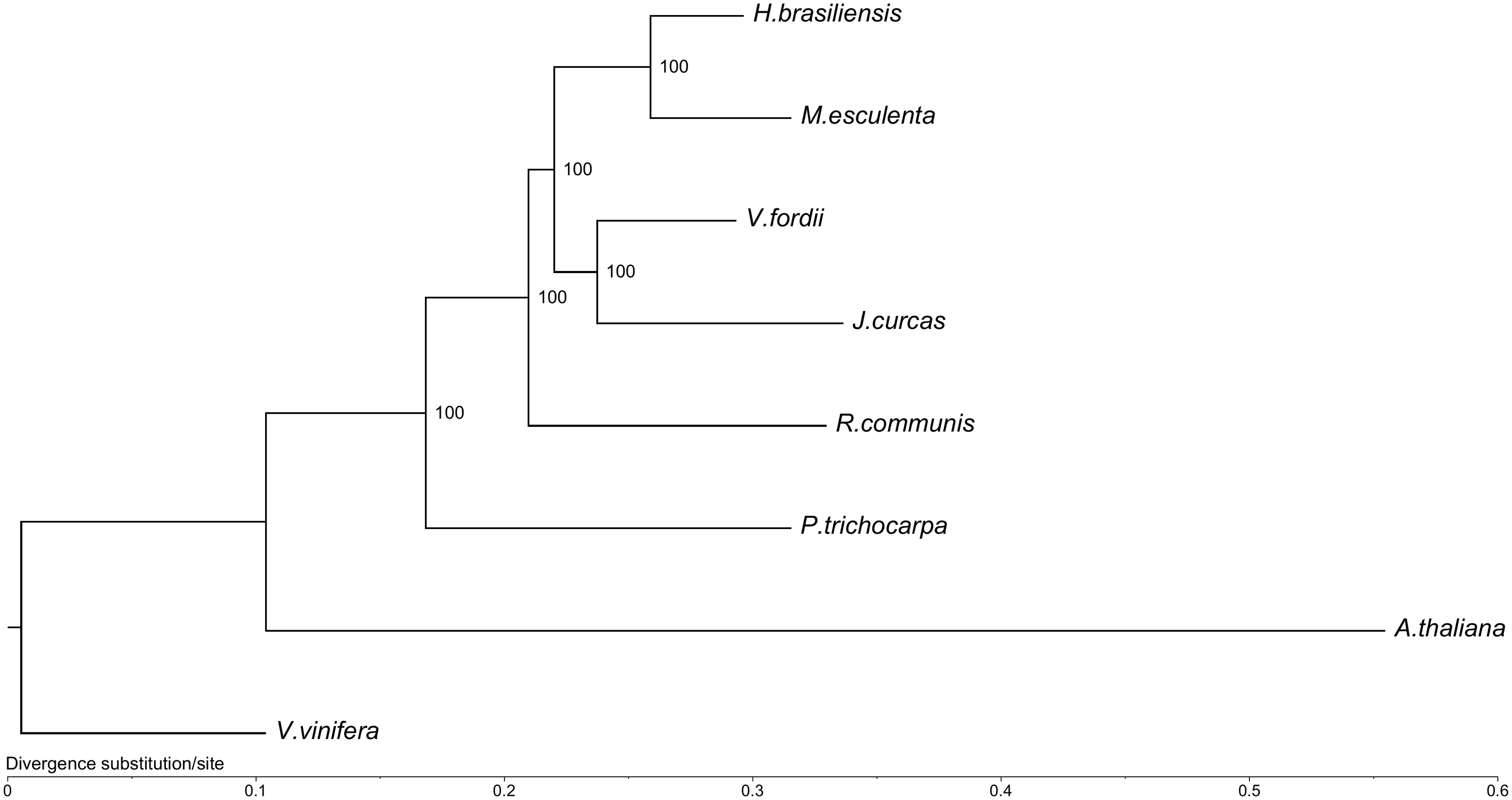
**Supplementary Figure S12. Phylogenetic tree of tung tree and seven other plant species.**

### Figure S13

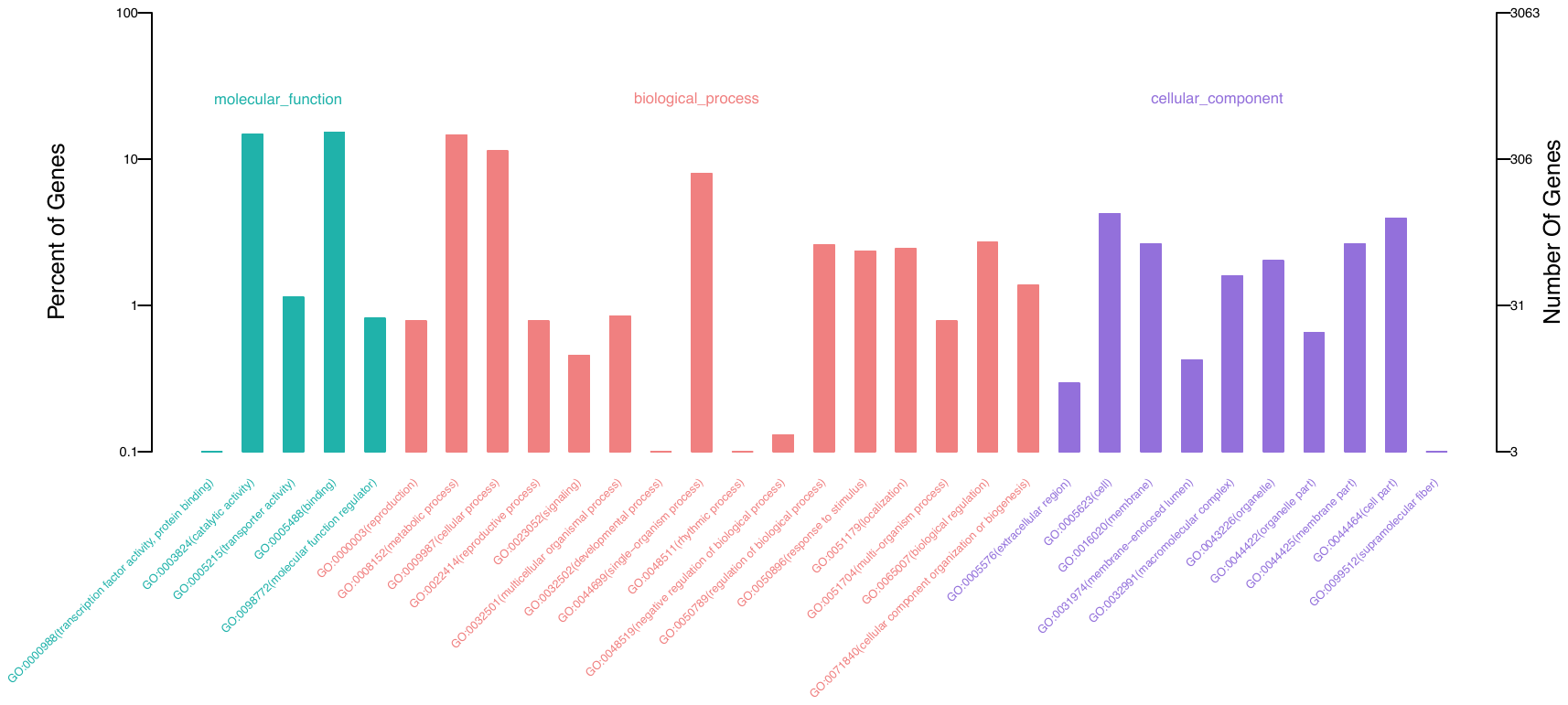

**Supplementary Figure S13. GO classification of tung tree expanded gene families.**

### Figure S14

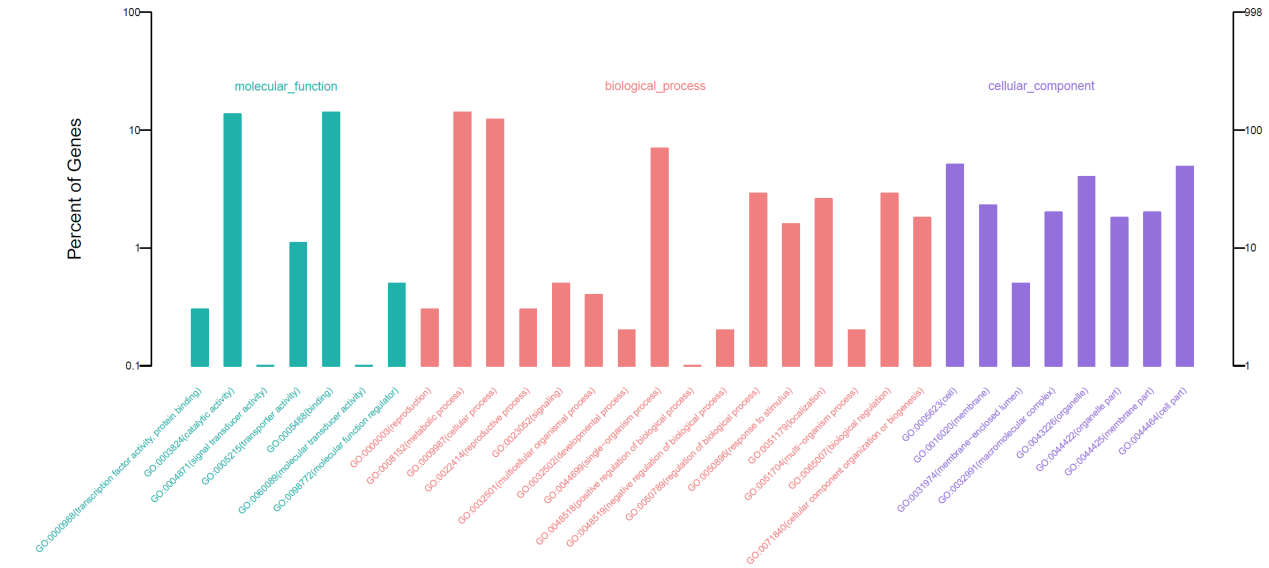

**Supplementary Figure S14. GO classification of tung tree positively selected genes (PSGs).**

### Figure S15

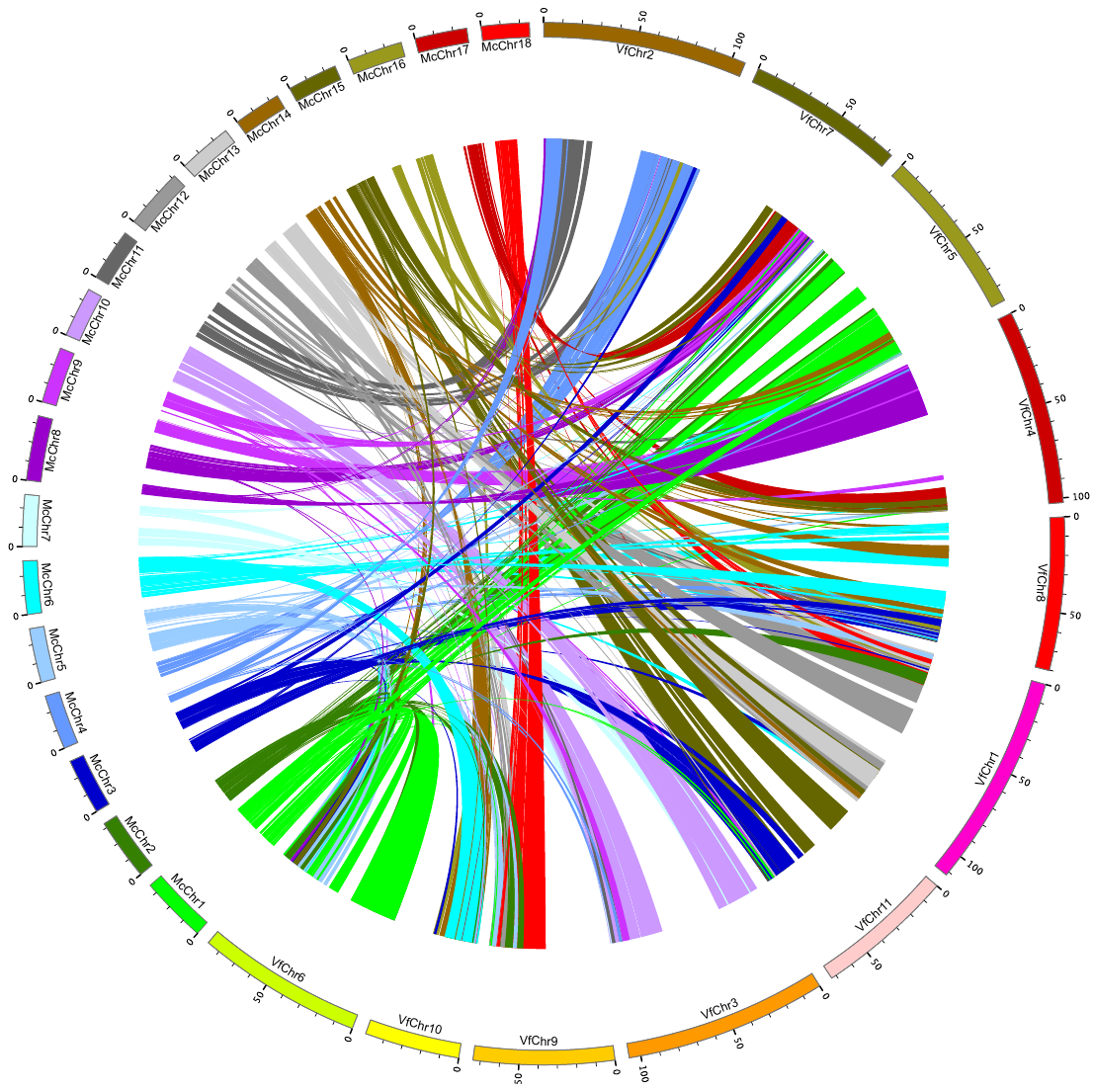

**Supplementary Figure S15. Synteny analysis between *V. fordii* and *M. esculenta*.**

### Figure S16

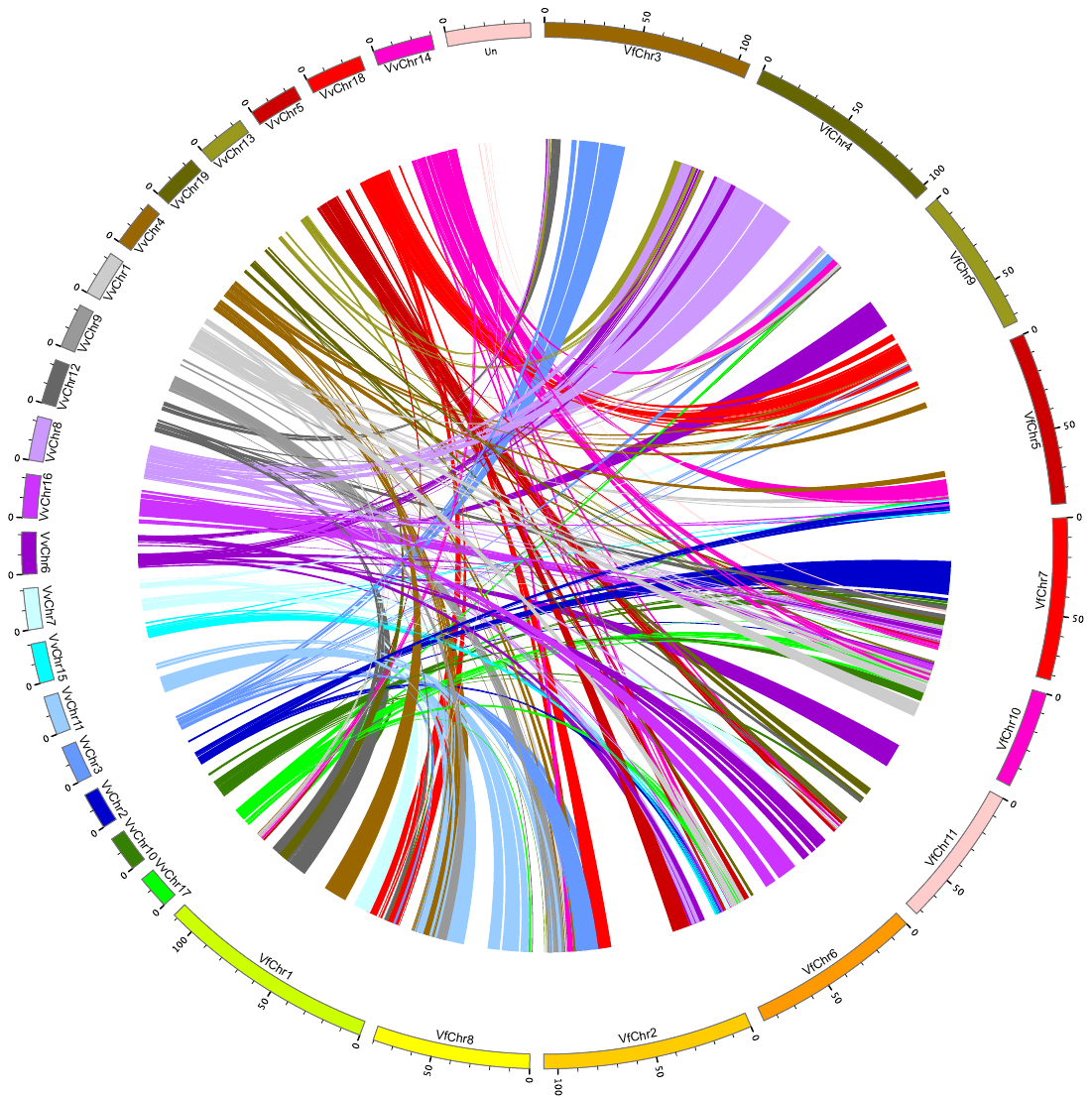

**Supplementary Figure S16. Synteny analysis between *V. fordii* and *V. vinifera*.**
