## Supplementary material for "The Tung Tree (*Vernicia Fordii*) Genome Provides A Resource for Understanding Genome Evolution and Oil Improvement": Figure S7

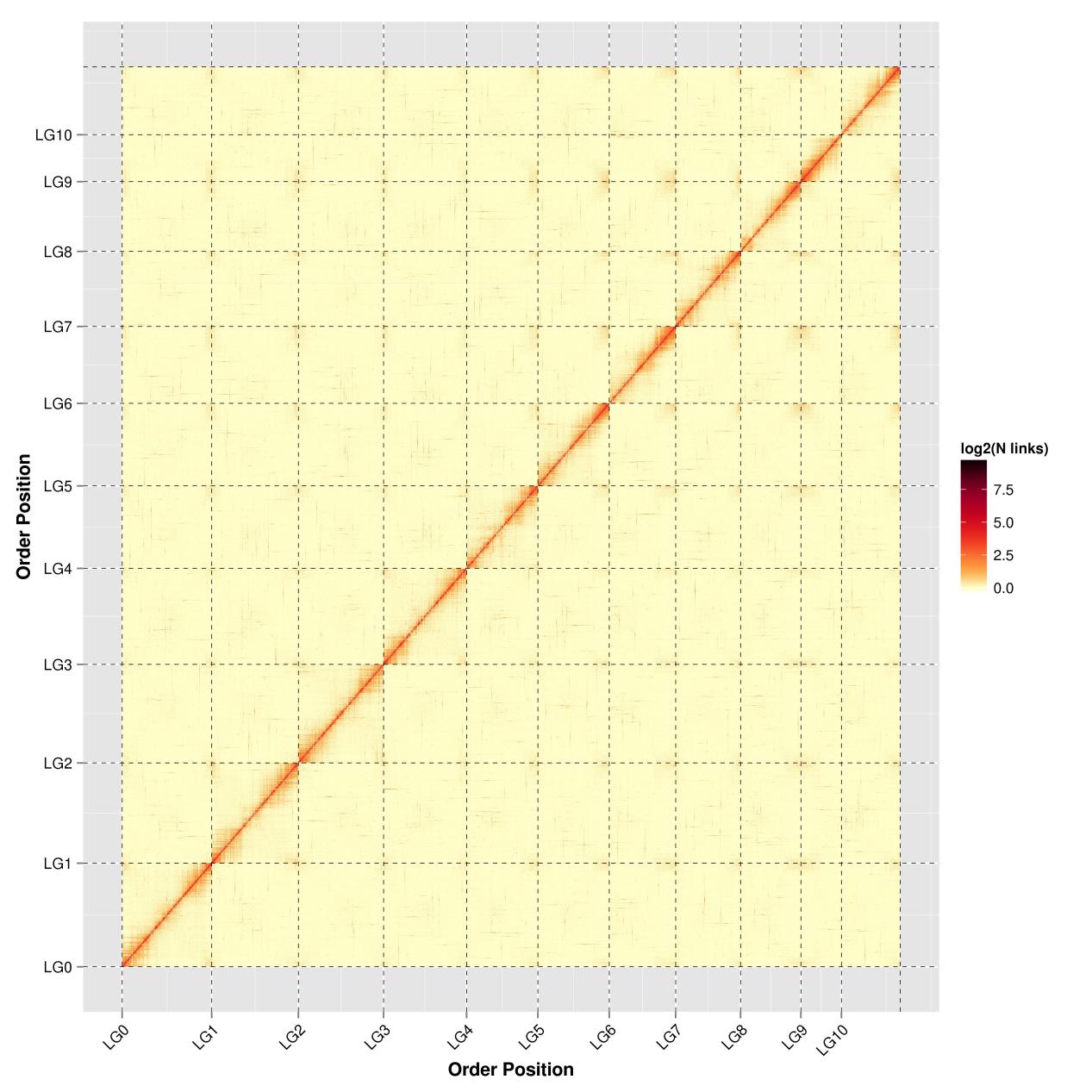

**Supplementary Figure S7. Hi-C linkage density heat map of assembled contigs.** LG0-LG10 represents Lachesis Group0-10, respectively. The x-axis and y-axis indicate the order position in the corresponding linkage group.
