## Supplementary material for "The Tung Tree (*Vernicia Fordii*) Genome Provides A Resource for Understanding Genome Evolution and Oil Improvement": Figure S8

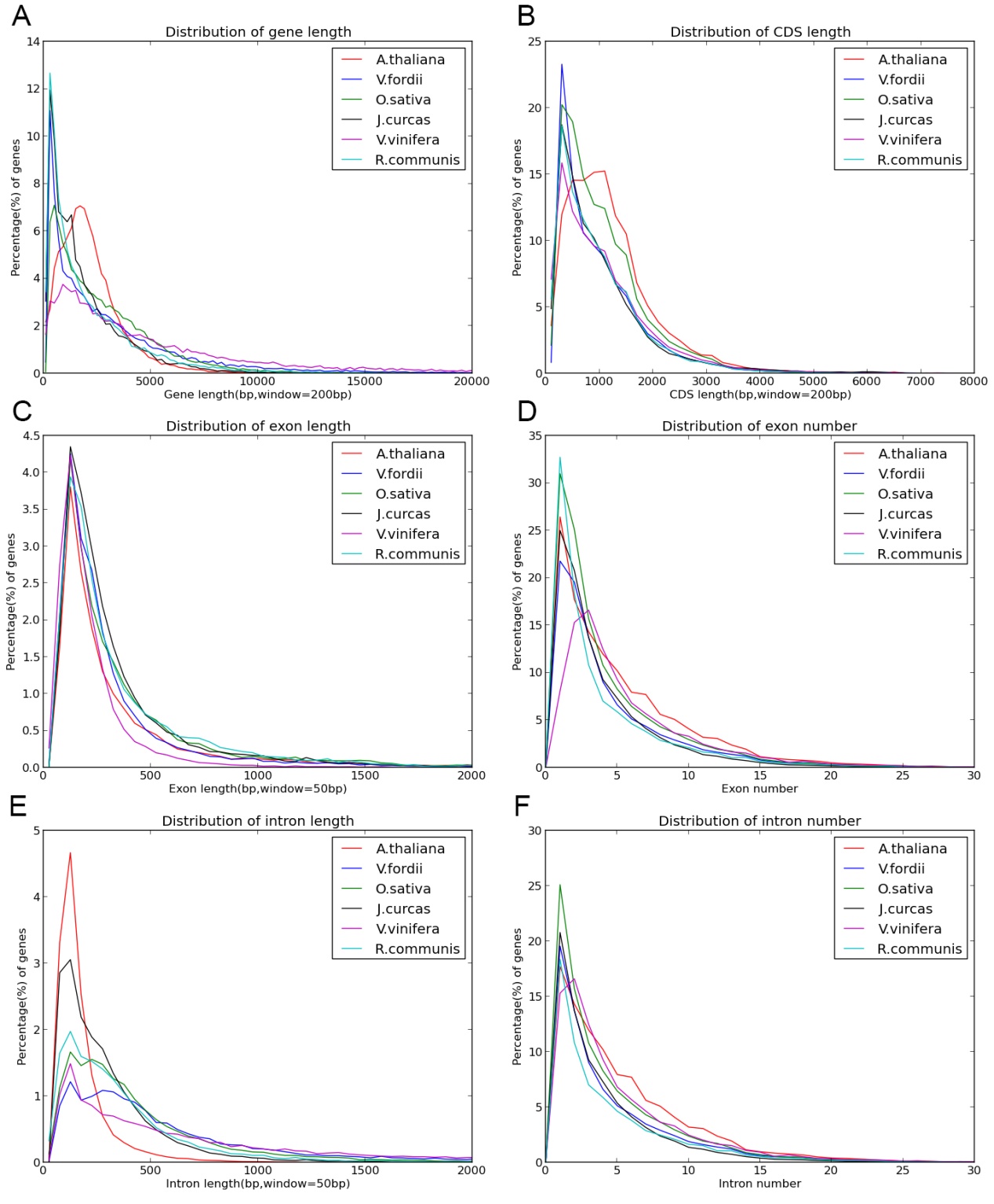

**Supplementary Figure S8. Cross-species comparison of gene elements between tung tree and other five species.** A: Distribution of gene length; B: Distribution of CDS length; C: Distribution of exon length; D: Distribution of exon number; E: Distribution of intron length; F: Distribution of intron number.
