## Supplementary material for "The Tung Tree (*Vernicia Fordii*) Genome Provides A Resource for Understanding Genome Evolution and Oil Improvement": Figure S9

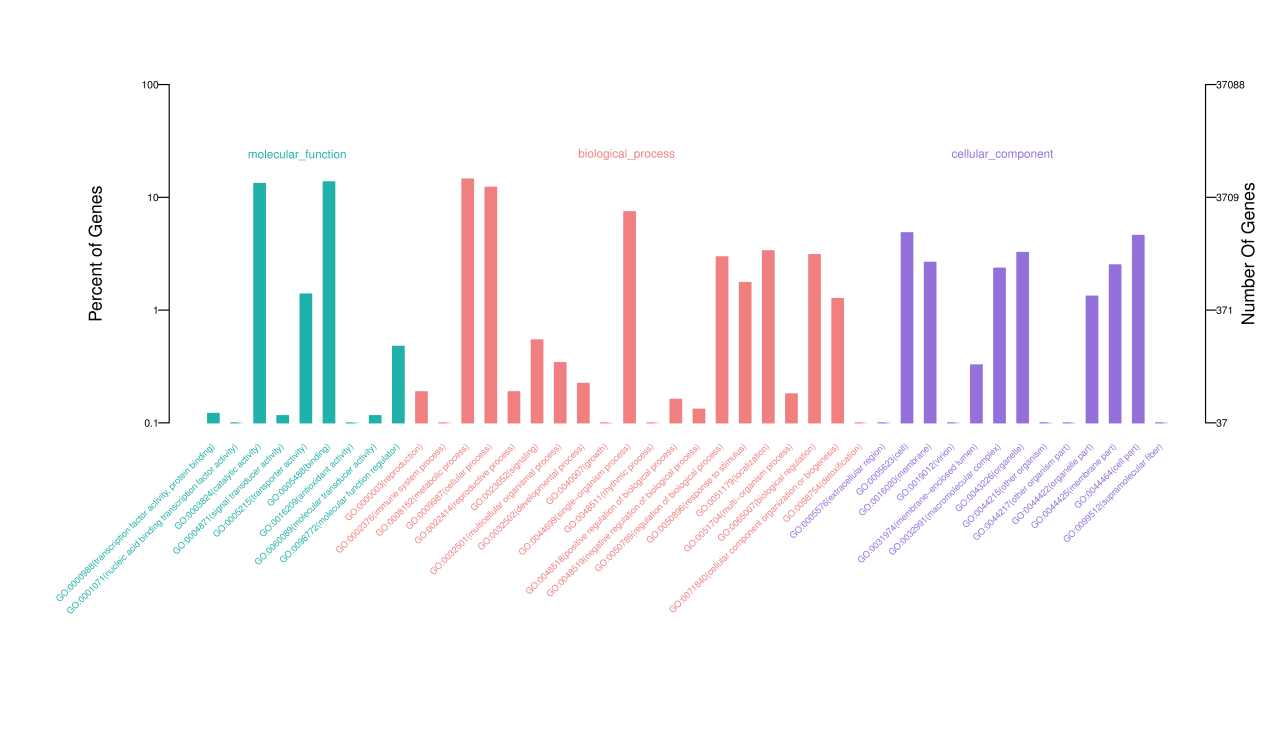
 **Supplementary Figure S9. Gene GO classification of tung tree genome.**

The results are summarized in three categories: molecular function, biological process and cellular component.
