## Supplementary material for "The Tung Tree (*Vernicia Fordii*) Genome Provides A Resource for Understanding Genome Evolution and Oil Improvement": Figure S11

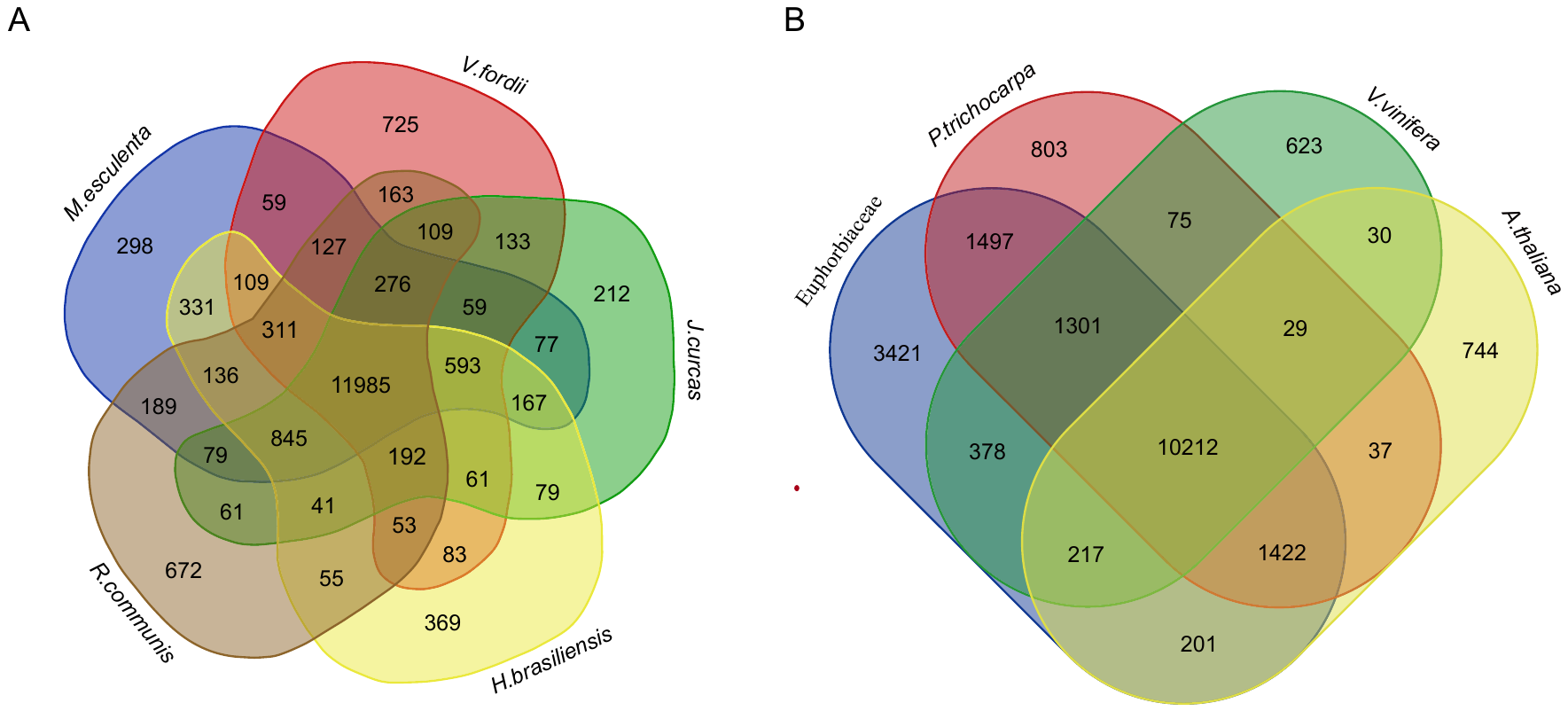

**Supplementary Figure S11. Venn diagrams of cross-species gene family comparisons.** A: The shared orthologues among five species in Euphorbiaceae; B: The shared orthologues among Euphorbiaceae, *A. thaliana*, *P. trichocarpa*, and *V. vinifera*. Each number represents a gene family number.
