## Supplementary material for "The Tung Tree (*Vernicia Fordii*) Genome Provides A Resource for Understanding Genome Evolution and Oil Improvement": Figure S17

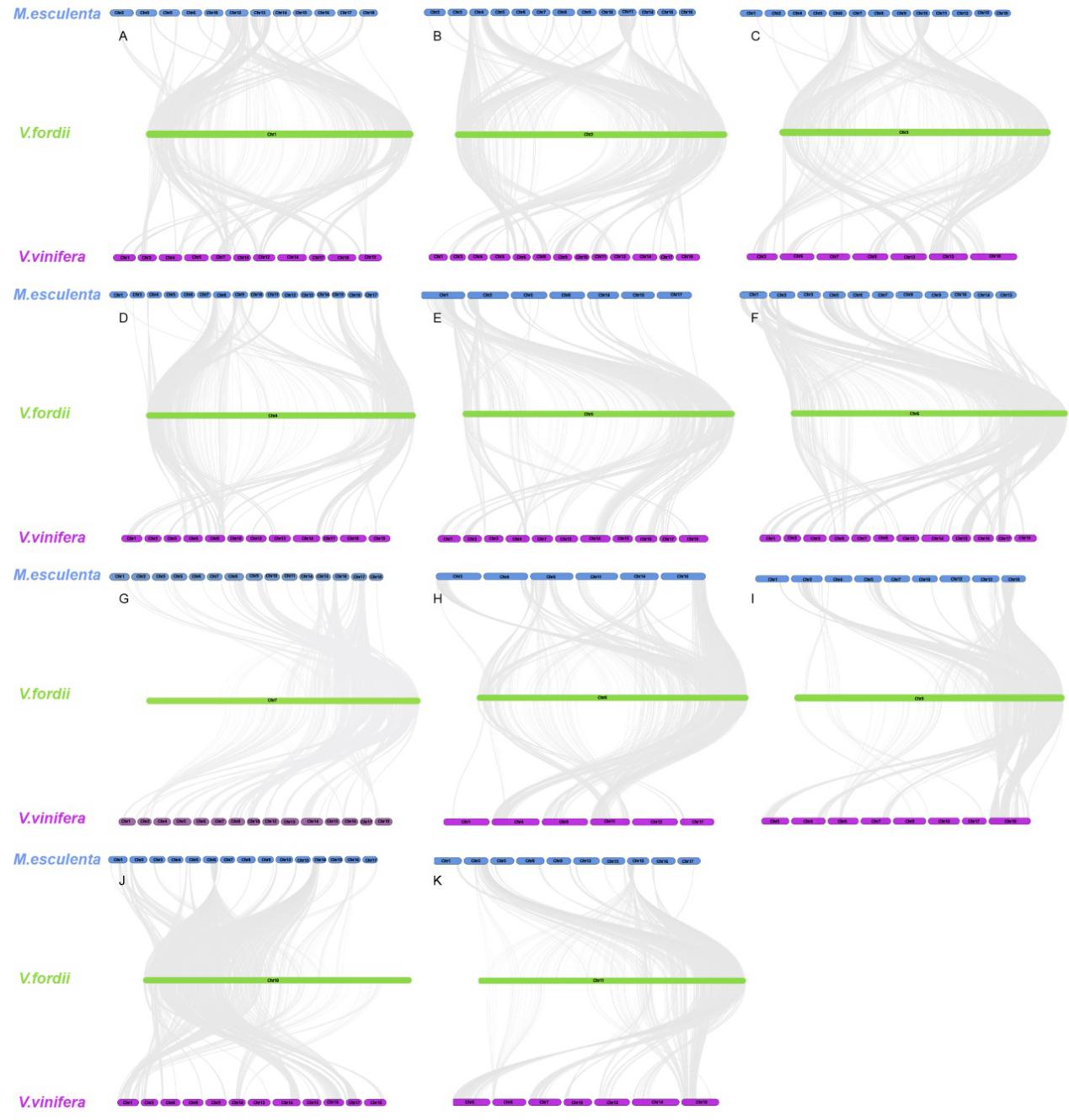

**Supplementary Figure S17. Collinear relationship of *V. fordii*, *M. esculenta* and *V. vinifera*.** A-K indicates each of the 11 tung tree chromosomes showing collinear relationship with *M. esculenta* and *V. vinifera*, respectively.
