## Supplementary material for "The Tung Tree (*Vernicia Fordii*) Genome Provides A Resource for Understanding Genome Evolution and Oil Improvement": Figure S18

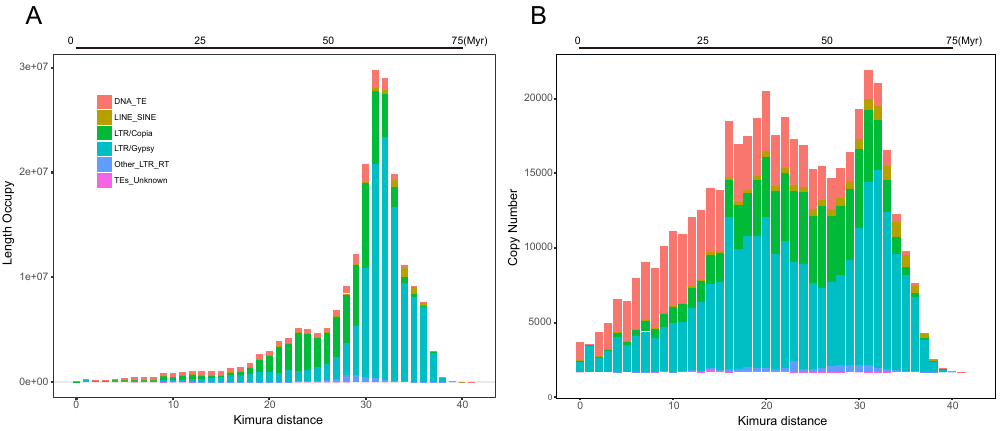

**Supplementary Figure S18. Evolutionary history of TE super-families in tung tree genome.** A: The occupied TE lengths; B: copy number of TEs. Kimura distances stand for the percentages of substitutions in matching regions compared to the consensus sequences to classify TEs.
