## Supplementary material for "The Tung Tree (*Vernicia Fordii*) Genome Provides A Resource for Understanding Genome Evolution and Oil Improvement": Figure S19

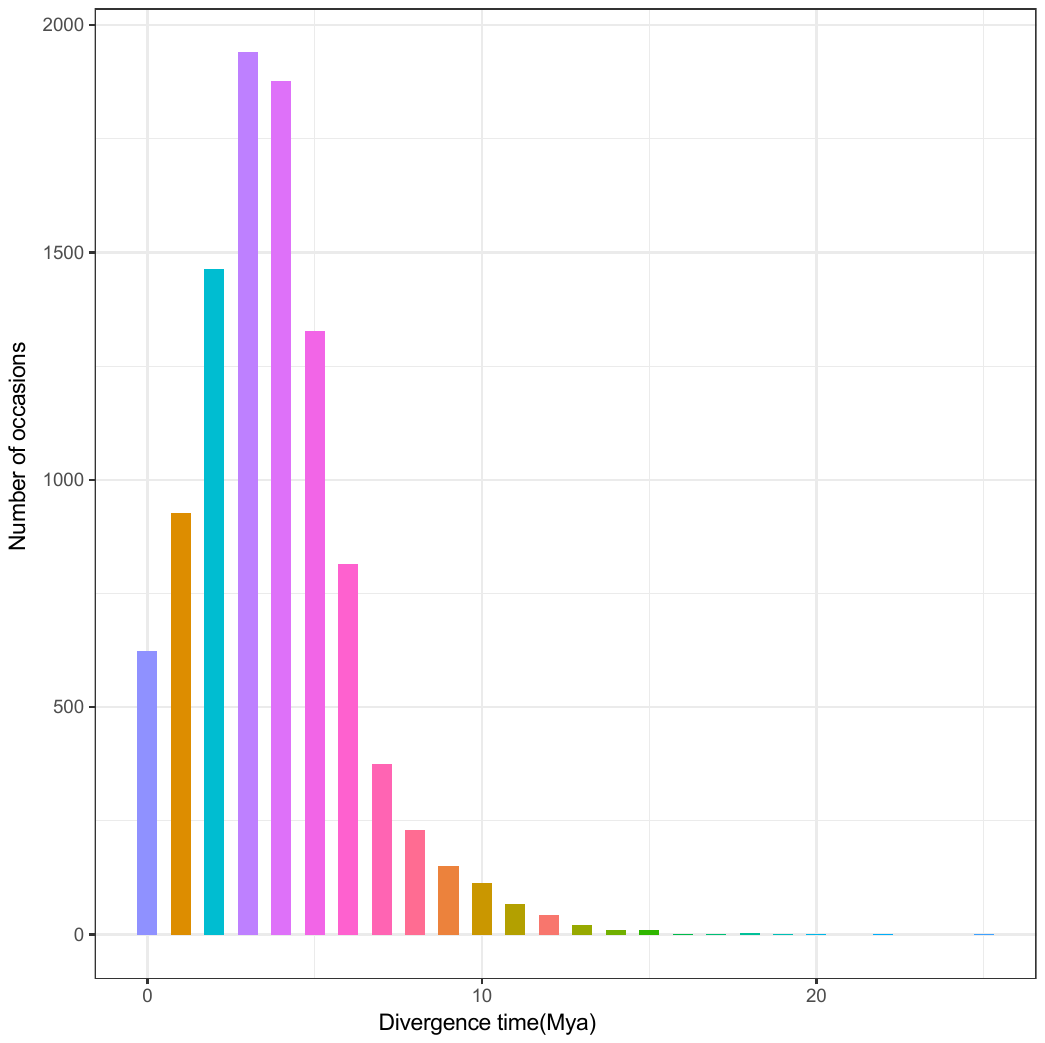

**Supplementary Figure S19. The insertion times for intact LTR retrotransposons in tung tree genome.**
