## Supplementary material for "The Tung Tree (*Vernicia Fordii*) Genome Provides A Resource for Understanding Genome Evolution and Oil Improvement": Figure S20

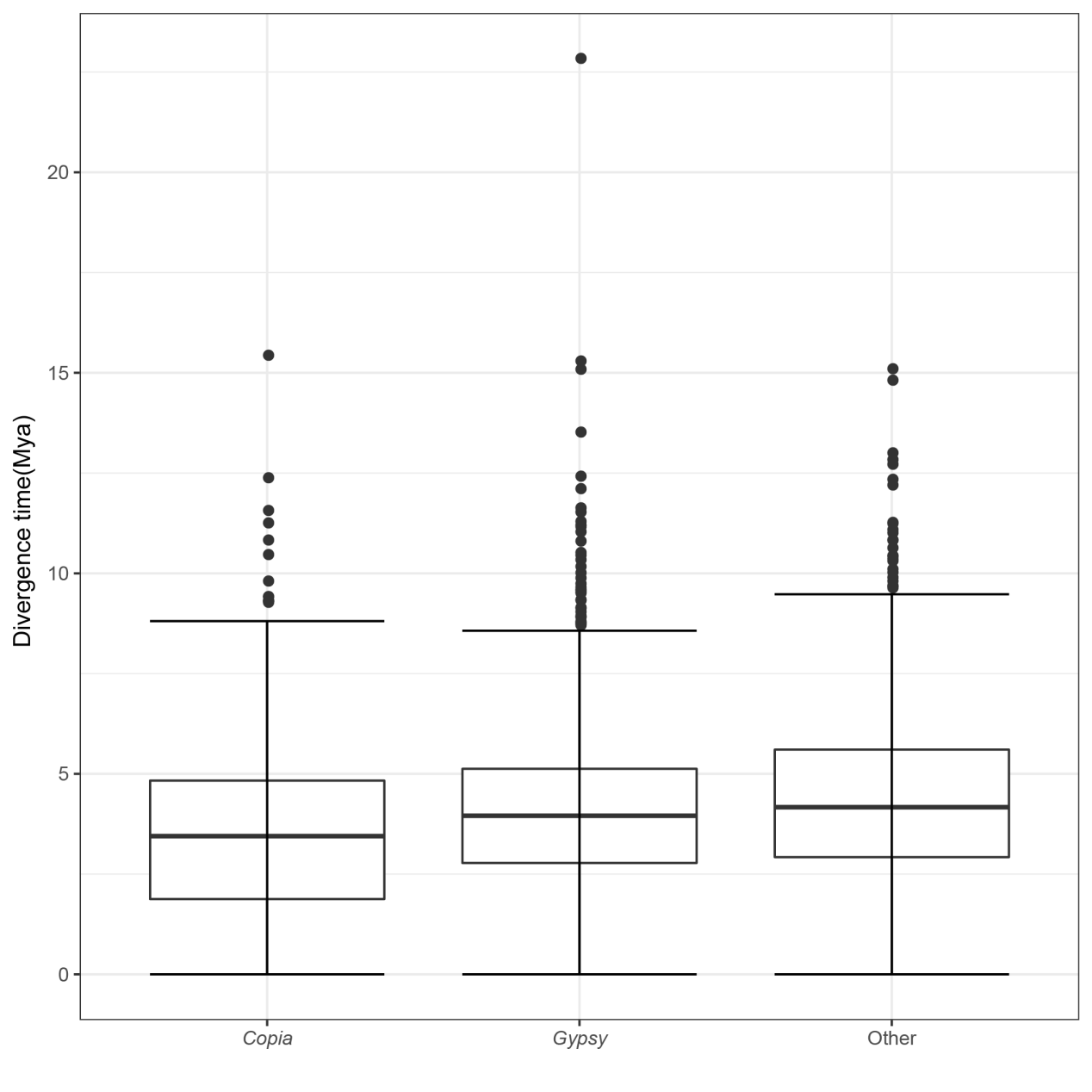

**Supplementary Figure S20. Insertion times of Ty1/Copia, Ty3/Gypsy and other LTR retrotransposon families in tung tree genome.**
