## Supplementary material for "The Tung Tree (*Vernicia Fordii*) Genome Provides A Resource for Understanding Genome Evolution and Oil Improvement": Figure S21

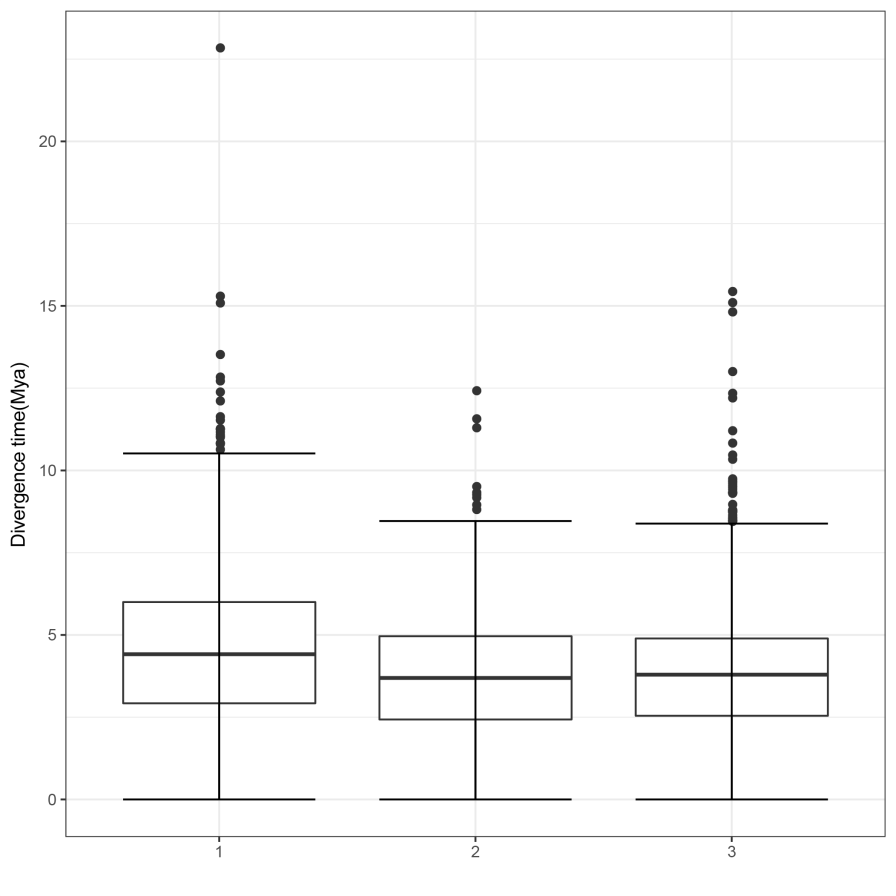

**Supplementary Figure S21. Insertion times of various copy number of retrotransposon families in tung tree genome.** Three types of families are shown, including single-copy families (1), median-copy families with 2-4 intact member (2) and high-copy families with >= 5 intact member (3).
