## Supplementary material for "The Tung Tree (*Vernicia Fordii*) Genome Provides A Resource for Understanding Genome Evolution and Oil Improvement": Figure S23

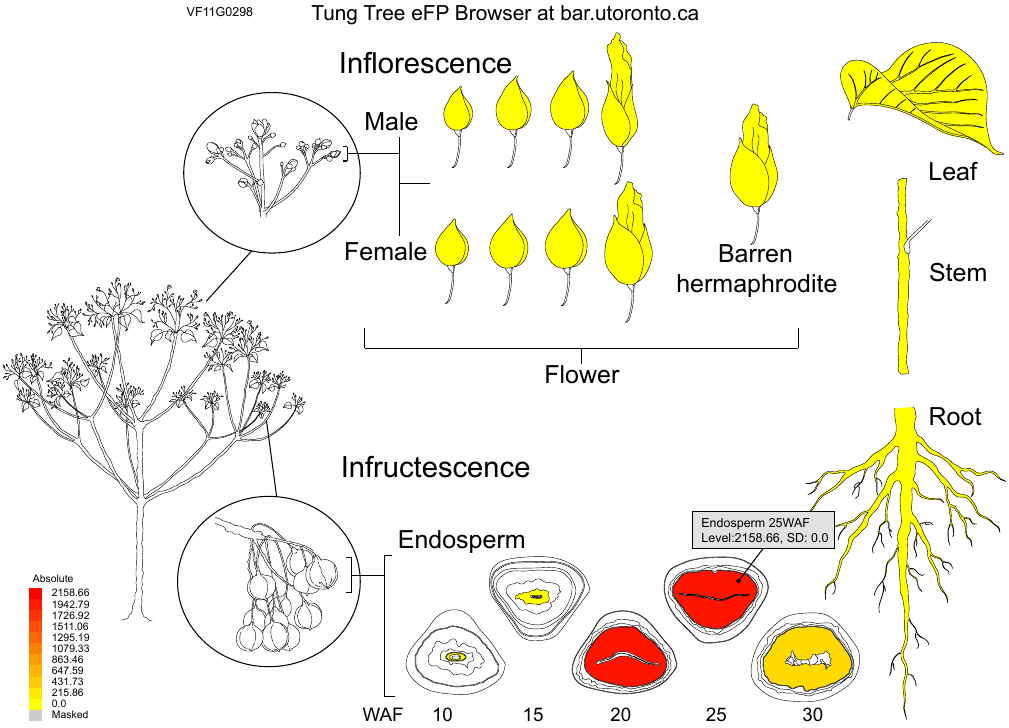

**Supplementary Figure S23. eFP browser view of gene expression pattern in tung tree.** A tung tree eFP browser output image showing expression values for the VfFADx-1 gene (Vf11G0298) is shown. Red indicates higher levels of transcript accumulation and yellow indicates a lower level of transcript accumulation.
