## Supplementary material for "The Tung Tree (*Vernicia Fordii*) Genome Provides A Resource for Understanding Genome Evolution and Oil Improvement": Figure S24

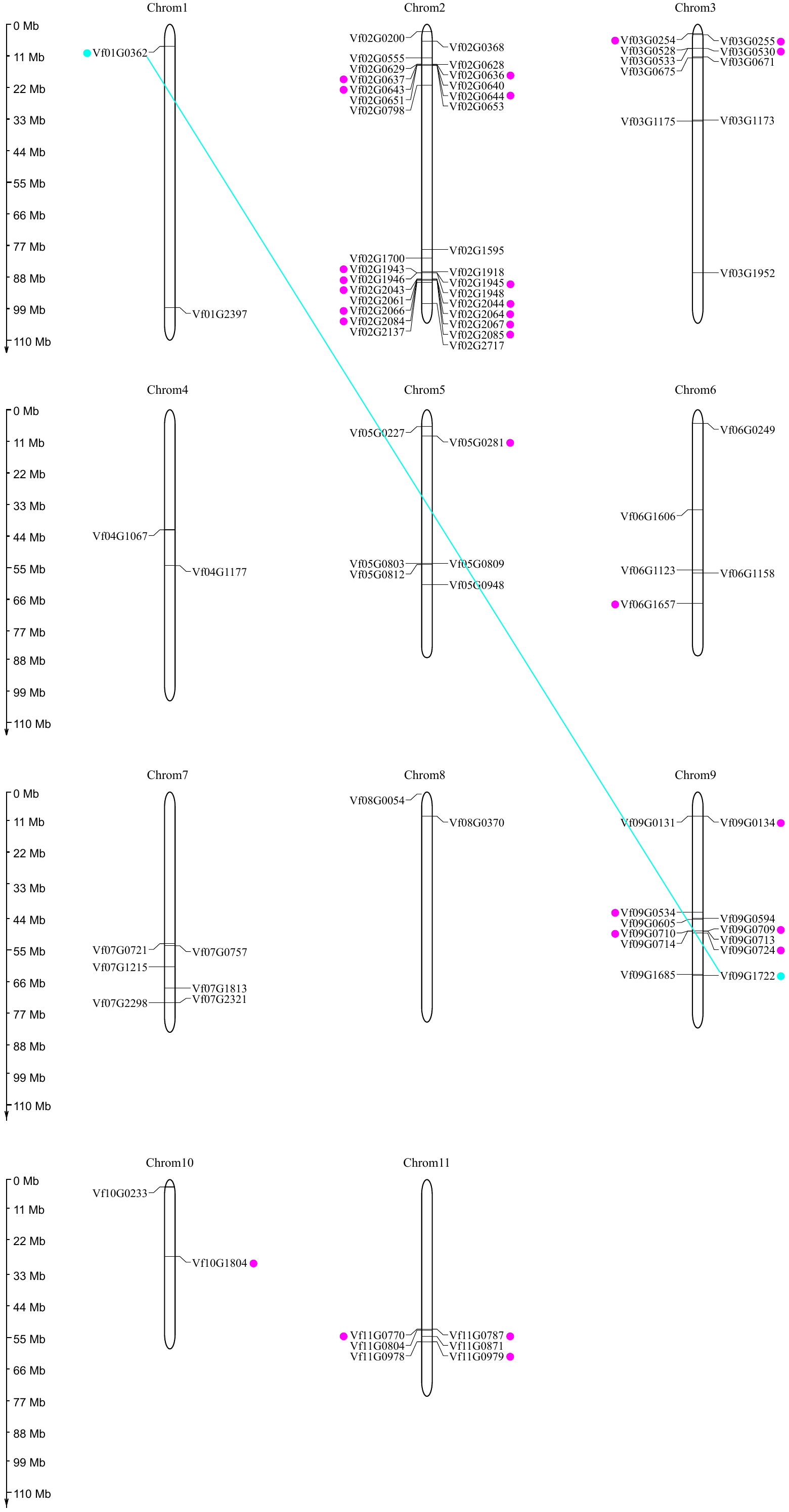

**Supplementary Figure S24. Chromosomal locations and region duplication for tung tree NBS genes.** The Chromosomal position of each NBS gene was mapped according to the tung tree genome. The chromosome number is indicated at the top of each chromosome. The scale is in mega bases (Mb). The segmental duplicated genes are indicated by blue dots and linked by blue line. The tandemly duplicated genes are indicated by pink dots.
