## Supplementary material for "The Tung Tree (*Vernicia Fordii*) Genome Provides A Resource for Understanding Genome Evolution and Oil Improvement": Figure S25

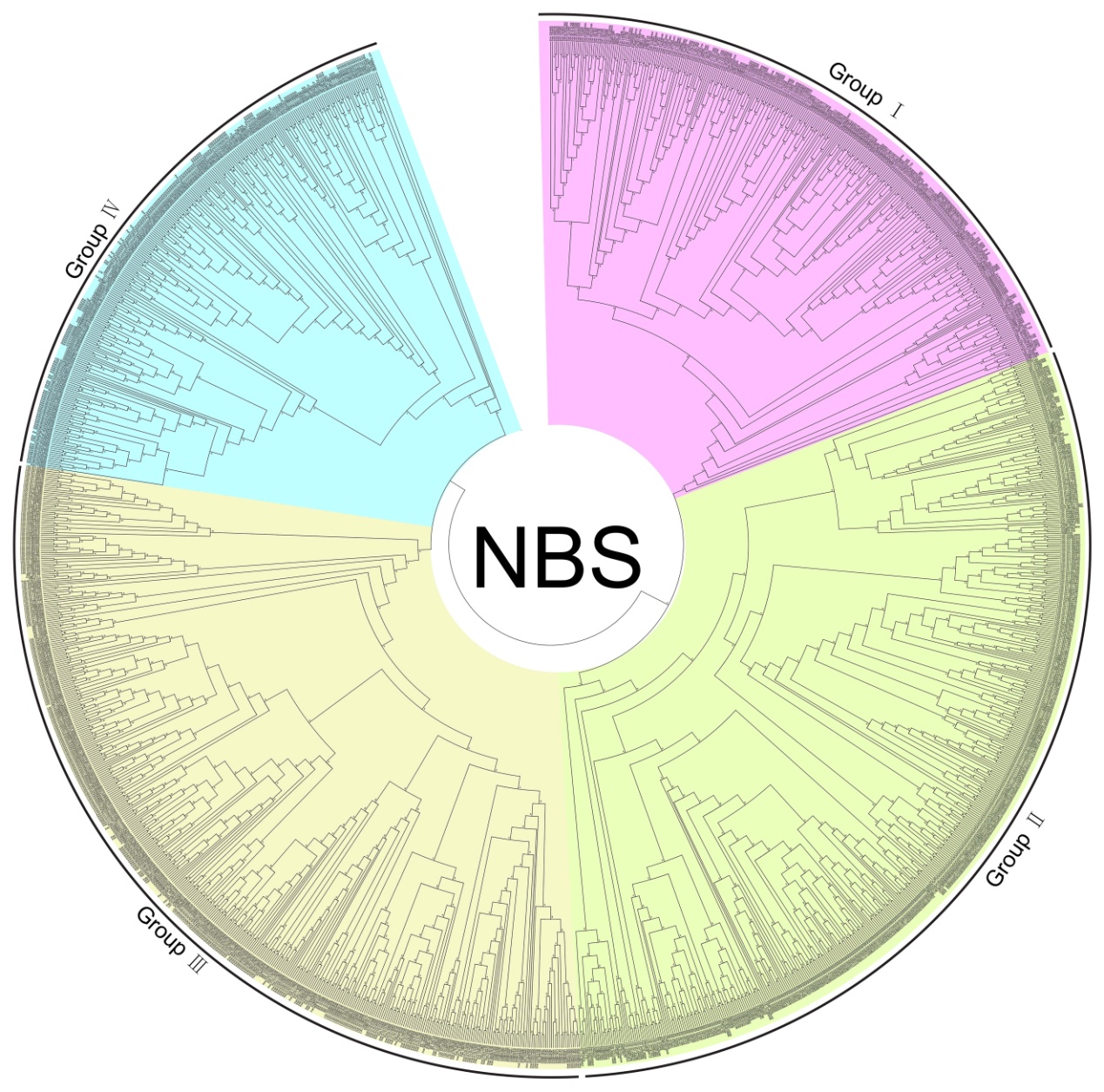

**Supplementary Figure S25. Phylogenetic analysis of NBS-encoding genes.**

A maximum-likelihood phylogenetic tree constructed from 1,530 protein sequences from *V. fordii*, *J. curcas*, *S. indicum*, *O. sativa*, and *P. trichocarpa*. The taxon names in the phylogenetic tree are indicated after gene ID. The clades are marked by four different block colours in the tree. The last one (yellow), a basal angiosperm, *A. trichopoda*, used as an outgroup; the monocot FAD2, eudicot FAD2 and udicot FADx clades are marked in red, blue and green, respectively.
