## Supplementary material for "The Tung Tree (*Vernicia Fordii*) Genome Provides A Resource for Understanding Genome Evolution and Oil Improvement": Figure S31

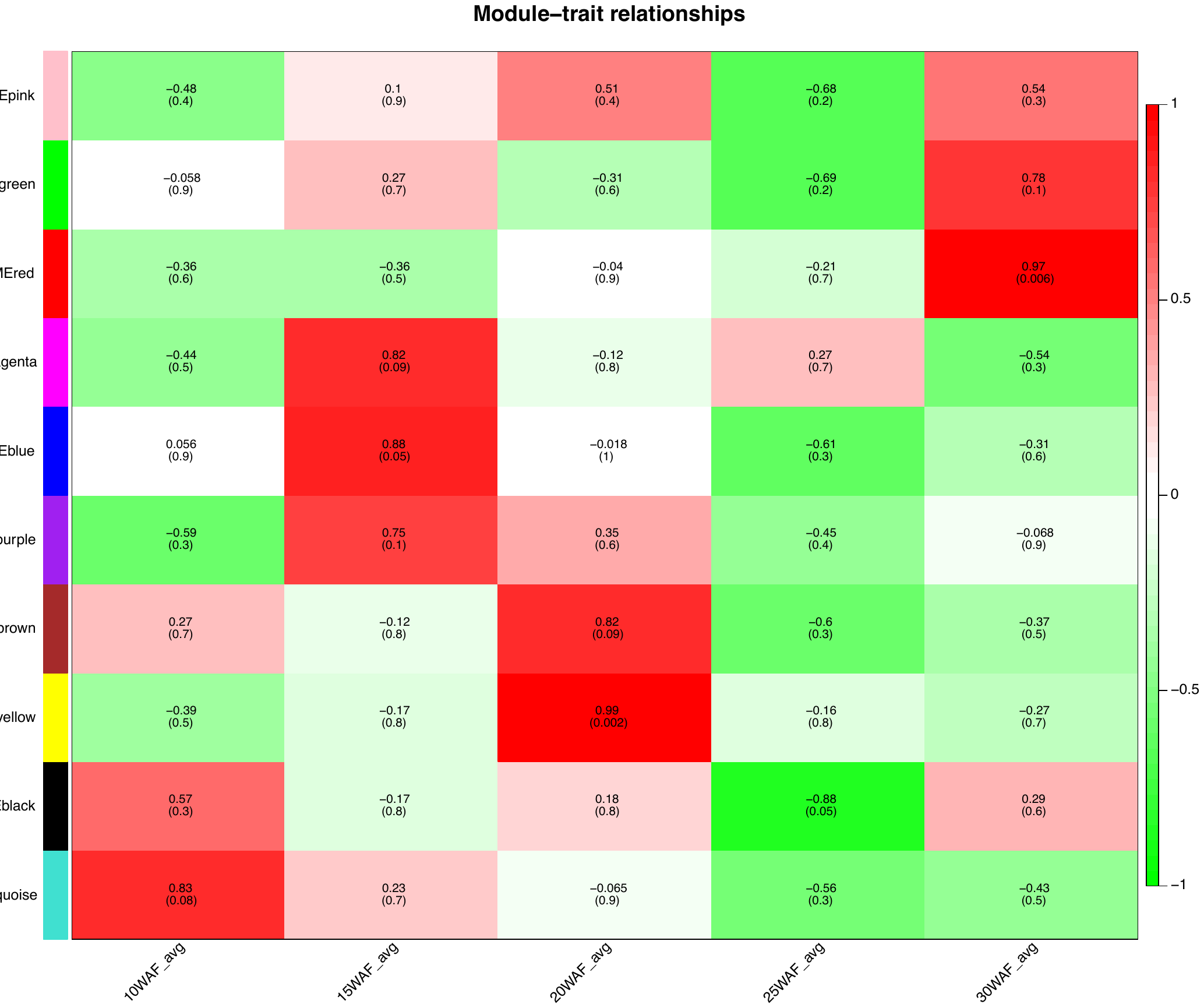

**Supplementary Figure S31. The relationship between co-expression module and trait in tung tree.** The color blocks on the left side are the co-expression modules constructed from RNA-seq data of five developing seed samples. The numbers underneath the module names are the number of genes in each module. The number represents the correlation between co-expression module and trait. The number in parentheses represents the p-value (Student's t-test) of the correlation.
