## Supplementary material for "The Tung Tree (*Vernicia Fordii*) Genome Provides A Resource for Understanding Genome Evolution and Oil Improvement": Figure S32

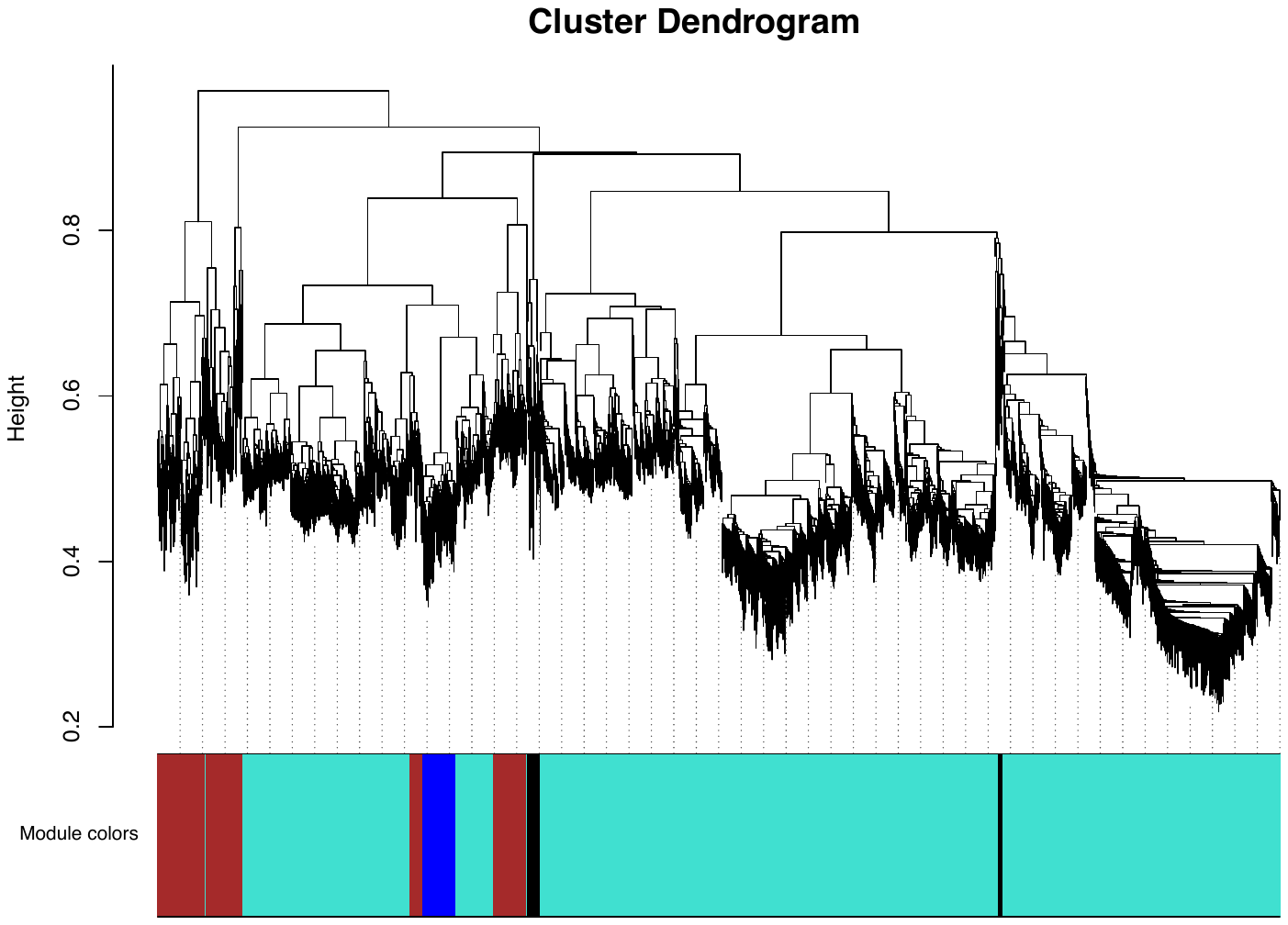

**Supplementary Figure S32. Co-expression network analysis of genes in developing seeds in tung tree.** Hierarchical cluster tree showing co-expression modules identified by weighted gene co-expression network analysis of 16,732 genes whose FPKM values≥1 in each seed sample. The major tree branches constitute 10 distinct co-expression modules labelled by different colors as indicated by a color band underneath the tree.
