## Supplementary material for "The Tung Tree (*Vernicia Fordii*) Genome Provides A Resource for Understanding Genome Evolution and Oil Improvement": Figure S33

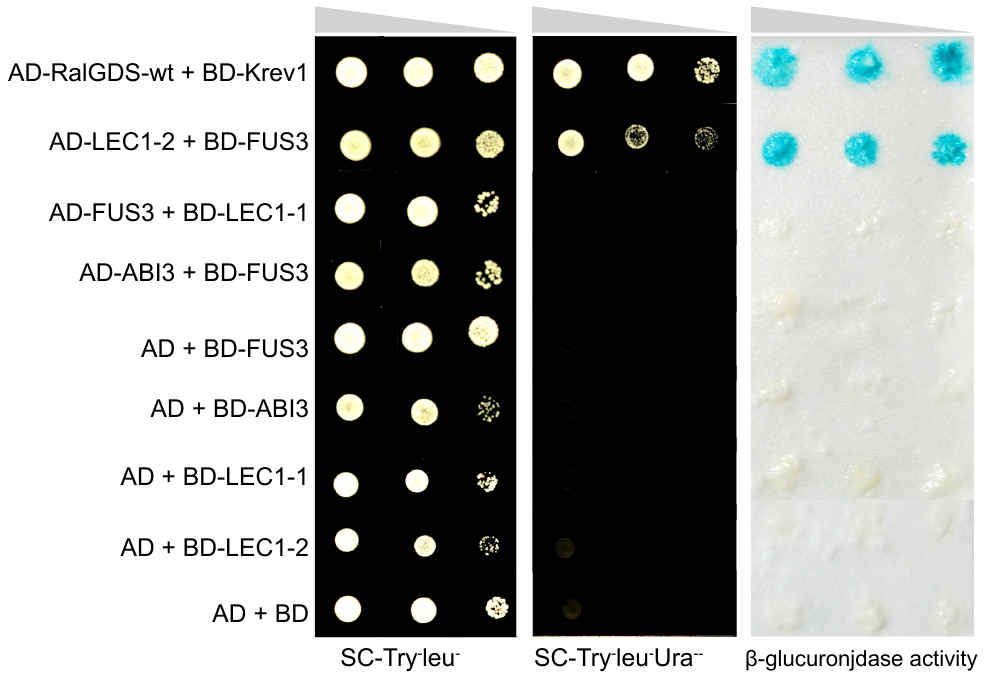

**Supplementary Figure S33. Yeast two-hybrid assay of transcription factors.**

AD + BD: negative control plasmids; BD-Krev1 + AD-RalGDS-wt: positive control plasmids;
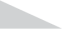
 indicates yeast concentrations from high to low.
